# A single cell atlas of sexual development in *Plasmodium falciparum*

**DOI:** 10.1101/2023.07.16.547819

**Authors:** Sunil Kumar Dogga, Jesse C. Rop, Juliana Cudini, Elias Farr, Antoine Dara, Dinkorma Ouologuem, Abdoulaye A. Djimde, Arthur M. Talman, Mara K.N. Lawniczak

## Abstract

The developmental decision made by malaria parasites to become sexual underlies all malaria transmission. Here, we describe a rich atlas of short and long-read single-cell transcriptomes of over 37,000 *Plasmodium falciparum* cells across intraerythrocytic asexual and sexual development. We used the atlas to explore transcriptional modules and exon usage along sexual development, and expanded it to include malaria parasites collected from a Malian individual naturally infected with multiple *P. falciparum* strains. We investigated genotypic and transcriptional heterogeneity within and among these wild strains at a single-cell level for the first time, finding considerable differential expression between different strains even within the same host. This work is a key addition to the Malaria Cell Atlas, enabling a deeper understanding of the biology and diversity of transmission stages.

**One sentence summary:** This addition to the Malaria Cell Atlas presents an analysis of sexual development and uses it to explore a natural infection.

## Introduction

Malaria parasites undergo asexual proliferation in the human host and this causes disease, however transmission from human to human can only occur when parasites successfully reproduce sexually in the mosquito vector. Sexual reproduction is initiated in the human host and depends on the facultative decision to commit to sex. This decision is controlled by the epigenetically-regulated transcription factor AP2-G (*1*, *2*) and depends on environmental factors, including competition, drug treatment, and host metabolic factors (*3–8*). Once sexually committed, human malaria parasites develop into male and female gametocytes that circulate in the bloodstream awaiting uptake by an *Anopheles* mosquito. Preventing sexual reproduction by stopping the parasite’s development at any point along its sexual cycle is critical for malaria control, as it breaks the cycle of transmission. Therefore, a deep understanding of how malaria parasites make decisions that lead to transmission is of great relevance to global health.

### Patterns of gene expression following sexual commitment to development

To fully describe the transcriptomic patterns during sexual commitment and development for *Plasmodium falciparum*, we used both short Illumina reads and full-length PacBio IsoSeq to carry out single-cell RNA sequencing along sexual commitment, development, and differentiation. After QC and data integration of short-read data, we retained 37,624 cells covering the asexual cycle and sexual differentiation and development (Fig. 1A, fig. S1, Materials and Methods). Although parasite differentiation and development are continuous processes, researchers have defined discrete stages based on morphology and gene expression. We deployed a consensus stage prediction method using a combination of clustering, reference mapping to both single-cell and bulk data sets, and identification of marker genes to label cells and annotate stages of asexual and sexual development (fig. S2-S4). UMAP visualisation portrays the asexual cycling of parasites as a circle and the differentiation into sexual forms as a stalk and two branches.

#### Gene expression signatures in sexually committed parasites

Our primary focus is on the developmental decisions that drive sexual commitment and sex determination, therefore, we subsetted and more finely labelled 8958 sexual cells to explore these key cell fate transitions (Fig. 1B, figs. S2 and S4). We observe a cluster at the base of the stalk emerging from the asexual trophozoites that is transcriptionally distinct from trophozoites and marked by expression of early gametocyte markers such as AP2-G, Pfs16, G27/25 and Pfg14-748, as well as several previously reported markers of sexually committed rings and schizonts such as HDA1, LSD2, REX4 and SURF8.2 among others (Fig. 1B, C, data S1) (*10*, *11*). We conclude that this cluster corresponds to the earliest sexually committed state sampled in our data, and termed it “committed”. Committed cells show a reduced repertoire of several genes associated with knobs and Maurer’s cleft that are involved in sequestering the asexual stages from the peripheral circulation by adherence to host endothelium, as previously reported (*12*) – among the 76 genes downregulated, 11 overlap with those reported to localise to these morphological features (*9*). Committed cells also express a set of ETRAMPs and GEXPs that may assist in homing and retaining these early stages in the extravascular spaces of the bone marrow and other tissues in natural infections (data S1).

**Fig. 1.**
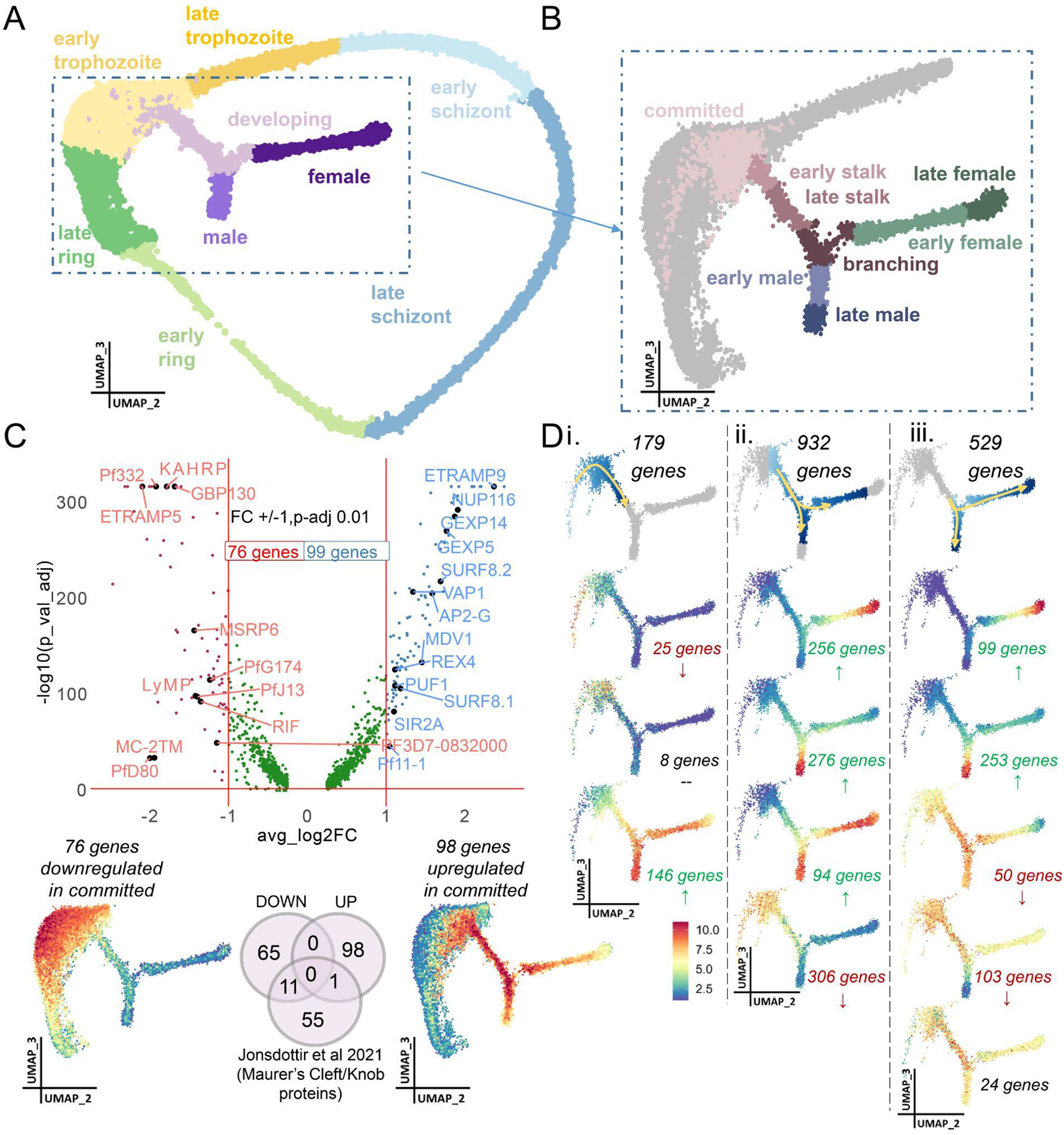
An atlas of gametocyte development. **(A)** UMAP of 37,624 single cells coloured by their assigned life cycle stage, and **(B)** the subset of 8958 differentiating sexual cells. **(C)** Expression in the committed cells was explored in comparison to the early trophozoites from which the sexual stages project. The volcano plot shows the genes identified via MAST (presented in data S1). The panel below shows averaged expression of upregulated and downregulated genes on the cell UMAP. Comparison of differentially expressed candidates with proteins localised to Maurer’s Cleft/Knobs (9) shows that committed cells downregulate 11 of these genes, as depicted in the Venn diagram. **(D)** Patterns of gene expression from commitment to differentiated sexes are explored and categorized into modules (i) following commitment, (ii) as the stalk gametocytes develop and branch into males or females, and (iii) following bifurcation, where males and females progress along their distinct sexual developmental trajectories. Modules are determined based on gene expression increases or decreases along the sexual lineages. Top UMAPs display the cells included in each comparison and the other UMAPs show the mean expression of all genes in each module. Complete results are presented in data S2.

#### Modules of expression from commitment to differentiated sexes

After sexually committed cells have departed the asexual cycle, we observed cells following a shared developmental path – the “stalk” – that are neither clearly male nor female followed by cells that are “branching” into either male or female developmental paths (Fig. 1A, B, figs. S2 and S4). To understand the developmental decisions that lead to these transitions, we performed a trajectory analysis and examined transcriptional changes along three paths: first, the stalk, then along the branching clusters leading to male and to female, and finally the progression through male and female development (Fig. 1D). For each of these three trajectories, we defined expression modules as described below and explored their underlying genetic programs.

#### Gene expression patterns within the stalk gametocytes

First, examining the stalk cells as they depart from the asexual cycle, we identify distinct gene expression patterns that we classify as modules S1-S3 (Fig. 1D, data S2, fig. S5). In module S1, we find several Maurer’s cleft proteins implicated in cytoadherence (CBP2/GEXP07, REX1-3, MAHRP1) (*13*) that show transcriptional decrease following commitment, as observed above in committed cells, as do AP2-G and GEXP02, an early target of AP2-G (*14*, *15*). Module S2 shows transient expression of exported proteins (ETRAMP5, ETRAMP9, MESA, GEXP14, GBP130, ACBP2) as well as two “Plasmodium exported proteins” of unknown function, perhaps indicating a role in gametocyte sequestration (*16–19*). This is followed by a transcriptional increase in genes affecting chromosome organisation (H3, H2A, H3 variant, H2B) and upregulation of several transcription factors (including c-myc, HDP1 and ApiAp2 family proteins) in module S3 (fig. S5, data S2). HDP1 is a key player for gametocyte maturation, facilitating transcription of genes essential for gametocyte development, including genes that are critical for the inner membrane complex and thereby gametocyte morphology (*5*, *20*). Md1, which has been recently characterised as a key player in sex determination and male development, is present in module S3 (*21*). Other uncharacterised nucleotide binding proteins within this module might similarly play roles in modulating gene expression at this critical juncture of sexual development.

#### Gene expression patterns as stalk gametocytes branch into female or male gametocytes

Second, we examine the stalk gametocytes as they branch into males or into females and increase sex-specific (modules E1 and E2) and sex-shared (module E3) transcripts (Fig. 1D, data S2, fig. S6). Male gametocytes begin to accumulate factors necessary for DNA replication, flagellar motility and chromatin condensation (module E2), while female gametocytes accumulate mRNAs that become translationally repressed as they are later necessary for zygote and ookinete development in the mosquito vector (module E1) (*22*) (fig. S6, data S2). We also find transient expression in exported proteins (module E4) putatively involved in erythrocyte remodelling in earlier gametocyte stages (*16*), suggesting that their continued expression may not be required for continued gametocyte sequestration (fig. S6, data S2).

#### Gene expression patterns along maturation of female and male gametocytes

Finally, we examine transcriptional patterns as the sexes mature after bifurcation is complete in the cell UMAP (Fig. 1D, data S2, fig. S7). Sex-specific genes in modules L1 and L2, the majority of which are also represented in modules E1 and E2, appear to be initially transcribed within the branching gametocytes and continue to increase in expression through maturation. Modules L3 and L4 include genes that were shared between the developing gametocytes prior to sexual bifurcation but appear to maintain their expression only along male (L3) or female (L4) lineages following bifurcation (fig. S7, data S2). Recent studies uncovered the role of an AT-rich interaction domain containing protein, PfARID, in regulating male gametogenesis and female fertility (*23*), while the knockdown of its *P. berghei* ortholog (*md4*) affects male gene expression (*24*). Interestingly, genes in modules E4, L3, and L4 that show transient expression along either or both sexual lineages are distinguished by a higher AT content in their putative promoter regions, suggesting a possible shared mode of transcriptional regulation of these modules (fig. S8).

#### Investigating male-ness or female-ness within the stalk gametocytes

The developmental stage at which a parasite commits to become either a male or female gametocyte remains unclear – e.g. the decision of whether to become male or female could be established upon sexual commitment or could take place as a second step along the route to becoming a mature gametocyte (*25–27*). Additionally, one sex could be the default sex with additional factors required to push the default into the other sexual state. While the cells sampled here reveal a developmental stalk in which gametocytes appear to be transcriptionally homogeneous before branching into male or female lineages, the UMAP is just a projection and it can fail to reveal subtle signatures of maleness or femaleness among these stalk cells. Therefore, we explored this late stalk to understand if there were any transcriptional patterns consistent with sex already being determined in the stalk cells. We selected 10 genes from module S3 that either show increased expression in the late stalk in the male lineage compared to the female lineage or vice versa, before bifurcation between the sexes is observable in the cell UMAP (figs. S5 and S9). We used the expression of these 10 genes to rank and group cells in the late stalk into putative male-like and female-like developing gametocytes (fig. S9). Differential expression analysis between these male-like and female-like gametocytes identified 44 genes with likely sex-dependent increases in expression well before bifurcation is observed in the cell UMAP. These include several known male and female marker genes (fig. S9, data S3 and S4). The increase in transcription of these sex-specific genes in the stalk stages suggests that gametocytes become transcriptionally dimorphic and sexually determined earlier than at the point of bifurcation in the cell UMAP.

### Global view of gene expression based on coexpression patterns

#### Identification of gene coexpression patterns across intraerythrocytic development using UMAP reduction

Functionally unannotated genes still remain the majority in the *P. falciparum* genome, and classifying them based on their shared expression patterns with functionally annotated genes can reveal something about their functional relevance. To explore this further, we used UMAP to collapse a normalised, transposed cell-by-gene matrix and used k-means clustering to group these genes into 15 clusters (Fig. 2A, data S5, interactive version malaricacellatlas.org).

**Fig. 2.**
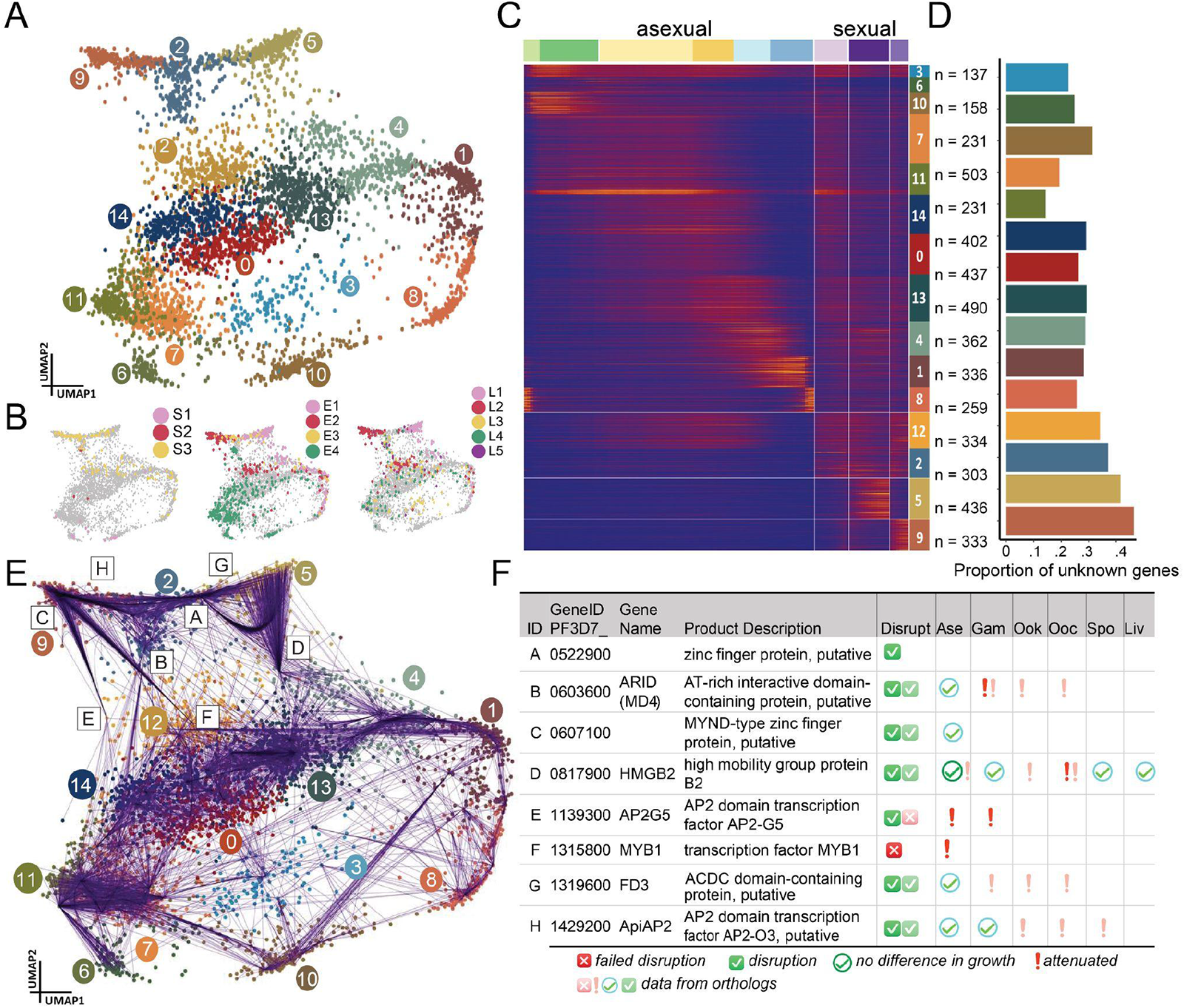
Patterns of gene co-expression across the life cycle stages. **(A)** UMAP of 5087 genes (each dot is a gene) based on their expression in each cell. Genes are coloured by their cluster assignment, using k-means (k=15) clustering on the predicted UMAP dimensions. **(B)** Transcriptional modules described in Fig. 1D show considerable overlap with sex specific gene clusters 2, 5, and 9. **(C)** Heatmap of gene expression clusters, where cells were ordered after pseudotime and assembled to 300 cell pseudobulks. **(D)** Proportion of genes in each cluster that have no known function – this is higher for the sex specific clusters. **(E)** GENIE3-based gene regulatory network to infer potential regulators of the sexual gene clusters, showing the strongest regulatory network connections. **(F)** Top 8 candidates that appear to regulate genes in the sexually associated clusters, along with phenotypic information from Phenoplasm (28). X – refractory to deletion, boxed tick – successfully disrupted, circled tick – no difference in asexual growth, and ! – attenuated phenotype in the life cycle stages (Ase – Asexual, Gam – Gametocyte, Ook – Ookinete, Ooc – Oocyst, Spo – Sporozoite, Liv – Liver). Lighter coloured icons indicate data from orthologs. Full list of genes ranked by number of connections is presented in data S6.

The gene graph revealed a striking structure with most genes clustering according to their use across the life stages (Fig. 2A, data S5). The gene graph displays three gene clusters specifically associated with sexual development and differentiation (2, 5, and 9) that also share genes with many of the modules described above (Fig. 2B). In some cases, these modules also contain genes found in other clusters, perhaps indicating recycled gene usage in asexual and sexual stages, or indicating genes that play a role in pushing asexual cells onto the sexual trajectory (Fig. 2B). Together, these three clusters contain the majority of sex-specific genes including 40 of 41 previously identified ‘gold standard’ sexual markers (*29*) (Fig. 2C). Importantly, these three clusters have a higher proportion of genes of unknown function than all other clusters, and genes contained within them should be explored as potential transmission blocking targets (Fig. 2D).

Gene Ontology (GO) analysis reveals similarly informative functional associations, as illustrated by the clusters 1 and 8 (top GO Cellular Component terms, ‘apical part of cell’ and ‘rhoptry’) comprising a significant number of proteins used in invasion and egress of the intraerythrocytic forms (data S5). Genes in cluster 9 predominantly exhibit expression in male gametocytes, aligning with the presence of gold standard sexual markers (PF3D7_0115100, PfsMR5, PF3D7_0219100, PF3D7_1325200, PUF1), and are enriched in GO terms related to cytoskeleton and motility. Interestingly, cluster 2, which exhibits consistent expression throughout all sexual stages (consisting of gametocyte markers P48/45, Pfs230, Pfg27, MDV1, and Pfs16) and cluster 5 with a predominant expression in female gametocytes (consisting of female gametocyte markers Pfs25, pCCp5, PFs47, NEK2, and NEK4) present GO terms related to monocarboxylic and fatty acid biosynthetic processes as top terms, warranting further exploration towards possible metabolic targets for transmission-blocking interventions (data S5).

#### Inferring interactions of transcription factors with sexual gene clusters using Gene Regulatory Networks

We next employed an unbiased gene regulatory network (GRN) inference approach to infer the interactions of 96 experimentally validated and manually curated transcription factors (*30*) in regulating these 15 clusters using GENIE3 (*31*). We illustrate the top eight transcription factors that have multiple regulatory connections to the sex clusters (Fig. 2E, F, data S6). FD3 and MD4/PfARID show a high degree of connectivity to the female and male clusters respectively, and play a role in gametocyte fertility in *P. berghei* and *P. falciparum* (*23*, *24*, *32*). Disruption of HMGB2, AP2-O3 and PF3D7_0522900 exhibit reduced growth in the ookinete or oocyst stages, indicating that downstream players of these factors may affect transmission (*33–37*). Finally, we find an association of the male cluster and AP2-G5, which is known to regulate AP2-G and is essential for gametocyte maturation (*1*, *38*). Although GRN inferences have limitations (*39*), these are promising candidates for potential roles in sexual determination and development.

### Interrogating sexual development using single-cell IsoSeq

#### Identification and classification of stage-specific isoforms by long read sequencing

Distinct isoform usage can drive developmental decision making and alternative splicing often plays an important role in sex determination and differentiation (*21*, *40*). To explore whether stage and sex specific isoforms might underlie critical decision points in the parasite, we generated single-cell IsoSeq data to capture full length transcripts for each of the five samples. After QC, for each sample we recovered more than 1 million unique transcripts with an average length of 1 kb (table S1, fig. S10). Reads from the five IsoSeq runs were merged and 37,614 cells were assigned stage labels based on short read data. For each stage, we typically obtained between ∼200K – 1M total reads and between ∼50 -230 reads per cell depending on the stage (fig. S10). We observed a good concordance of gene/UMI detection between long-read and short-read data (fig. S10). Gene expression signatures and cell type clusters were also consistent (Fig. 3A, B). Following QC and chaining of stage-specific isoforms to obtain unique isoforms (Methods), isoform diversity was explored within each stage and across all stages using SQANTI3 (figs. S10-S12, Methods) and presented as a data resource (data S7).

#### Revealing stage-specific exon usage, isoforms and novel splicing events using long read sequencing

The generation of cDNA in the 10X protocols should be full length but the relatively short length of our long-read data (average =1203 bp) compared to the expectation from gene annotations (average = 3022 bp transcript length on PlasmoDB) indicated that many reads do not fully span genes (explored in Supplementary Text). To account for this technical challenge of incomplete full length data, we used DEXseq (*41*) to investigate differential exon usage rather than isoform usage within and between the sexual and asexual stages.

We find differential exon usage in 6 genes between committed and asexual trophozoites, in 24 genes between males and females, in 39 genes between females and asexuals, and in 17 genes between males and asexuals (Fig. 3C, fig. S13, data S8). We present several representative illustrations of observed patterns (Fig. 3D-F, fig. S14). For example, GEXP13, an exported protein potentially in the PHIST family (*42*), shows a novel isoform appearing as the asexual stages commit to and proceed into sexual stages (Fig. 3D). This alternative isoform usage could influence sequestration or cell remodelling differences between sexuals and asexuals. We also observe distinct exon usage in Md1 (Fig. 3E), as recently reported (*21*). Md1 plays important and distinct roles in sex determination and development; female gametocytes lack exon1 expression of Md1 altogether and instead show antisense transcription driven by a specific alternative bidirectional promoter (*21*). Such antisense expression, associated with decreased exon usage, was also observed in other examples (fig. S15, S16) suggesting a common mode of regulation of stage specific processes. Another interesting candidate belonging to the Jumonji-type demethylase family, jmjc3, shows distinct exon usage profiles between gametocytes and asexuals, with the exon encoding its cupin-like domain not expressed in the sexual stages (Fig. 3F). Jumonji Inhibitors have been shown to disrupt gametocyte development and formation in *P. falciparum* (*43*). Among the candidate genes explored by IGV visualization, we noticed several cases where current gene models in the annotation appear to be inaccurate or incomplete (fig. S16). Conclusions from the long read data need to be carefully evaluated as we did detect some differentially expressed exons that are likely the result of spurious reverse transcription due to polyA stretches (fig. S17). DEXseq applied to the asexual stages also revealed several genes showing differential exon usage between rings, trophozoites, and schizonts, which should be further explored for their influence on asexual development (fig. S18). It is clear that single-cell long-read data reveal novel and dynamic isoform usage that may be important in developmental decisions, and that improvements to the annotation and to the technical ability to represent complete full length cDNA in single cells will enhance our understanding of the prevalence and importance of isoforms in parasite development.

**Fig 3.**
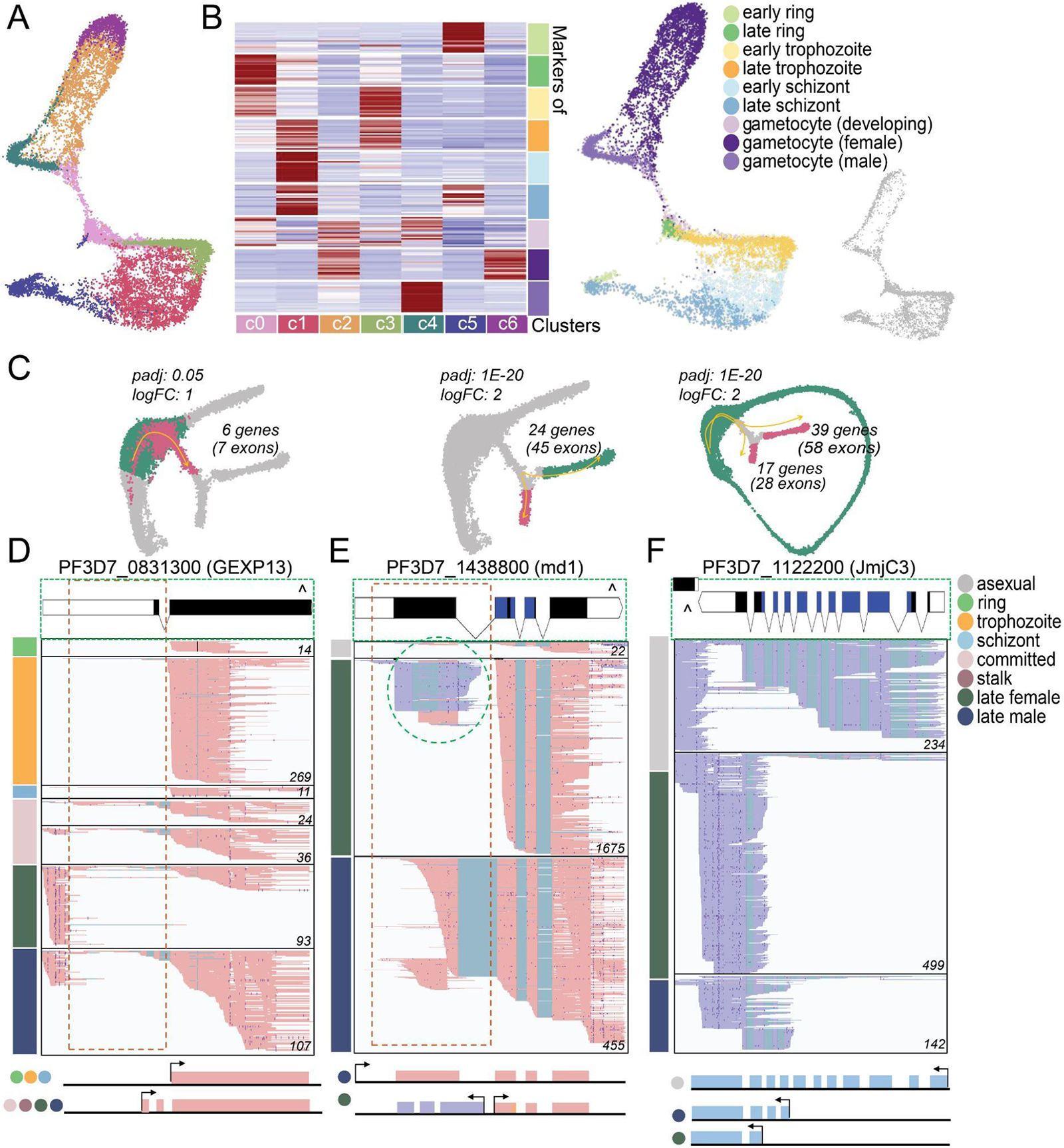
Long-read data reveals stage specific exon usage. **(A)** UMAP embedding of single-cell gene expression using long-read data echoes the topology inferred using short-read data. 17067 cells, with number of reads and genes > 20, are coloured according to the graph-based clusters determined using Seurat. **(B)** Identity of the above clusters is explored using the top 20 stage-specific markers from short read data. The heatmap in the left panel shows the summed expression across all the cells of each cluster for each stage-specific marker gene present. UMAP plots in the right panel show cells (12183 cells) in A coloured by stage labels assigned using short-read data. Cells not present in the filtered short read dataset (4884 cells) are displayed in a separate panel in grey. **(C)** Differential exon usage (DEU) between early trophozoite vs committed (left); female vs male (middle); and asexual vs female and asexual vs male (right) was completed. In D-F, long-read data is shown using IGV for three example genes, with PlasmoDB gene model on the top and inferred exon usage models on the bottom. PFAM domains in the gene model are displayed in blue. Summed coverage from IGV tracks is presented in the lower right corner. **(D)** Committed gametocytes and subsequent sexual stages show a previously undescribed isoform in the GEXP13 locus with an additional intron in exon 1. **(E)** In md1, female gametocytes lack exon1 expression, and display antisense expression of a lncRNA (highlighted by the oval dashed line). **(F)** jmjc3 shows usage of all annotated exons in the asexual stages while altering its profile in both the males and females.

### High-resolution stage and strain delineation in a natural infection using short and long read single cell transcriptomics

The complete blood stage developmental time course presented above can now be used as a reference atlas to deeply investigate natural malaria infections at the single cell level, which we do for the first time. We generated short and long-read scRNAseq data from asexual and sexual stage parasites collected from a 10-year-old naturally infected asymptomatic female Malian donor (fig. S19). In areas of high endemicity like Mali, people can carry different strains in a single infection and one of our aims is to better understand whether each strain contributes to the transmissible reservoir. We used the short-read scRNAseq genotypes and transcriptomes to first perform quality control (QC) and to assign a strain (*44*) and a lifecycle stage (*45*) to each parasite (fig. S20, S21, Methods). After QC we retained 1011 parasites comprising late rings, early trophozoites, males, and females as expected in an asymptomatic natural infection (Fig. 4A, fig. S21) (*46*). These were distributed across 8 souporcell assigned strains (Fig. 4B, fig. S22).

#### Exploring relatedness between strains

Two strains (SC2 & 6) had at least 30 asexual parasites and 30 gametocytes present (Fig. 4B, fig. S22). Other strains appeared to have only one of these stages represented (SC3 [asexual] and SC4 [gametocytes]) or were predominantly asexual (SC1) or sexual (SC5 & 7) (Fig. 4B, fig. S22). The imbalanced distribution of strains in the asexual and sexual compartments could be due to incomplete sampling, clearance of specific fractions by the immune system, differential propensities to sexually commit, or asynchronous sequestration. To validate souporcell findings and perform further relatedness analysis, we genotyped each strain’s asexual and/or gametocyte clusters, considering any cluster that contained at least 30 cells using a custom pipeline (fig. S22, S23, data S9, Methods). For SC2 and SC6, the asexual and sexual genotypes were in agreement with souporcell assignments confirming asexual and gametocytes assigned to the same strain were indeed genotypically identical, despite clustering separately on the genotype PCA plot (Fig. 4B, fig S24). We then called genotypes on strain clusters, using the same pipeline above, to estimate the genome proportions and locations identical by descent (IBD) between every pair of strains. Several strains share considerable fractions of their genome in IBD tracts suggesting they have very recent shared ancestry: SC1 shares 39% of its genome with SC2; SC2 shares 42% with SC6; and SC6 shares 45% with SC7 (fig. S24, S25). In spite of this, SC7 shares no recent ancestry with SC2 or indeed any other strain apart from SC6 (fig. S24, data S10). Other pairs had IBD ranging from 0-0.25 (fig S24, data S10). IBD results reflected the souporcell PCA clustering: SC7 was separated from all strains along PC1, apart from the SC6 cluster, with which it shares IBD (Fig. 4B). And SC1,2, and 5 are separated along PC2 reflecting their levels of IBD. Given the complex IBD patterns along the genome with no strain showing unrelatedness from all other strains, these likely represent recombinants from a single mosquito that ingested at least 5 genetically distinct gametocyte-stage parents from the previous host (fig. S24, S25, data S10).

**Fig 4.**
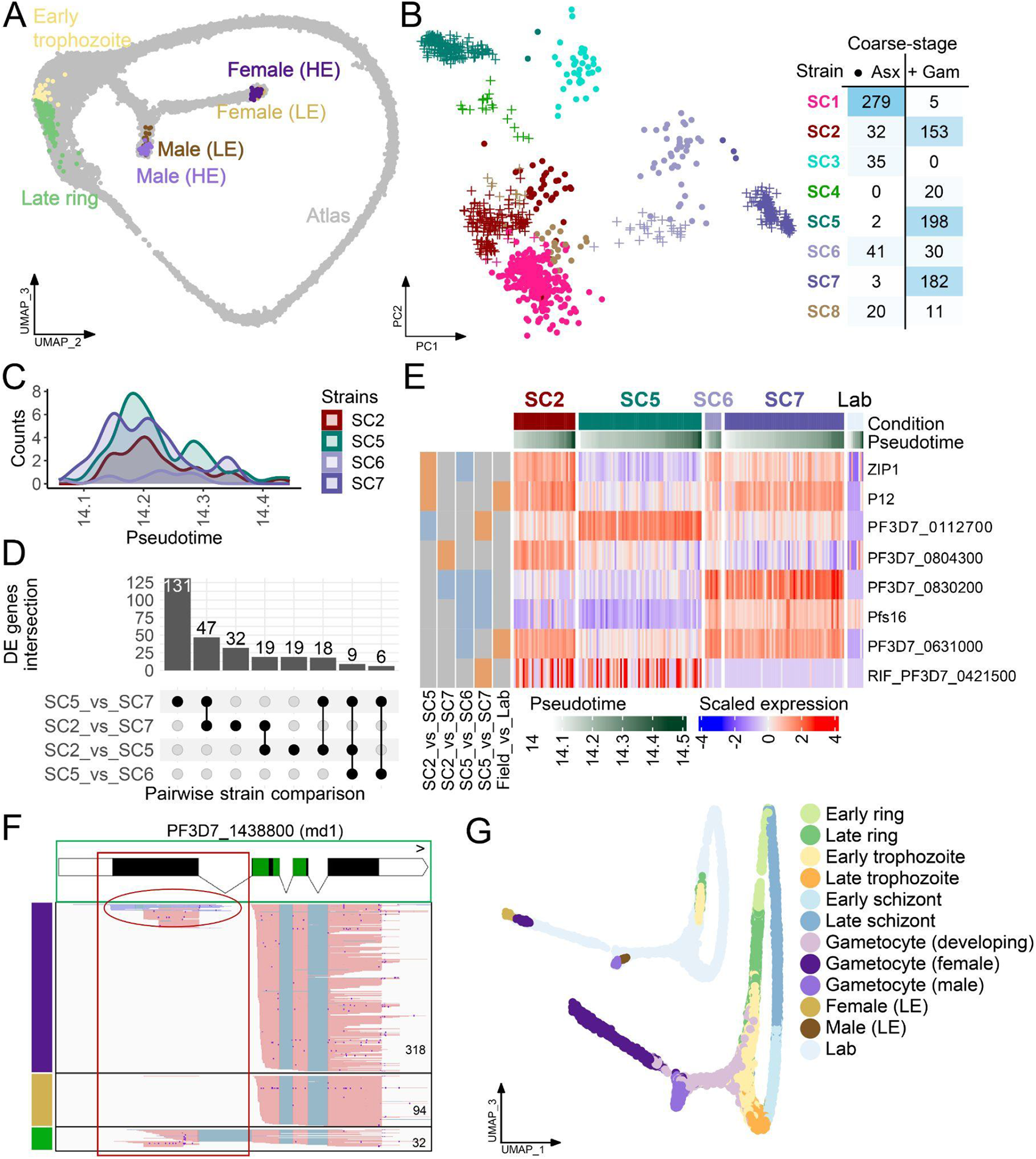
Strain and stage delineation of a natural infection. **(A)** scmap label assignment UMAP of the MCA lab atlas coloured based on the mapping of Malian donor parasites. **(B)** PCA on donor parasite genotypes, with strain represented by color and coarse-stage represented by shape (Asexual (•) or Gametocyte (+)). Total cell counts are in the table. **(C)** Distribution of female (HE) donor parasites coloured by strain along pseudotime. **(D)** Counts of differentially expressed (DE) genes (MAST p-value <0.05 & tradeseq waldStat >10) for each pairwise comparison of the strains containing sufficient numbers of females (HE) (SC2, SC5, SC6, and SC7). **(E)** Heatmap of top DE genes (average log2 fold change >1.5; MAST p-value <1e-10, & tradeseq waldStat >10) between strains in females (HE), with cells arranged according to pseudotime. The bars on the left are coloured orange if the gene is highly expressed in the first listed strain and blue if vice versa. The field_vs_lab bar shows the pairwise DE of all donor parasite strains combined versus the combined lab strains. **(F)** Differential exon usage between males and females of donor parasites in exon 1 of md1. **(G)** Integrated UMAP containing all lab and field parasites. LE gametocytes have been labelled separately because they may reflect new biology and are highlighted together with other field stages in the smaller top left UMAP.

#### Investigating the biology of cell types within the natural infection

Transcriptomic clustering revealed five cell clusters. Three clusters were confidently assigned as asexual, male, or female and these assignments were strongly supported by marker expression (fig. S21, fig. S26, data S11). Interestingly, the two remaining clusters had convincing expression of known male and female markers but weak assignment scores to gametocytes in the atlas presented above (*29*) (fig. S21). These two clusters expressed fewer genes overall and these genes were expressed at greatly reduced levels compared to the male and female clusters (fig. S26, S27). We therefore labelled the weakly assigned clusters as low-expression (LE) and the strongly assigned clusters as high-expression (HE) and explored these further (Fig. 4A, fig. S21). The LE males and females clearly express a subset of the genes expressed in their HE counterparts but they do not express any unique genes, which is not true of any other life stage (fig. S26, data S11). For each stage we defined a gene as being core if it was detected in at least 25% of the cells for GO analysis. GO analysis on core genes exclusive to LE and HE females shows an enrichment for the crystalloid component among other GO terms. Crystalloids are limited to the ookinete and oocyst but some genes encoding their proteins are expressed in female gametocytes (*47*). Although much more lowly expressed in LE females, genes and orthologs of genes whose disruption leads to a defective phenotype in mosquito stages (*28*) are detected in both LE and HE females (fig. S26, data S11). This includes genes that are known to be expressed in gametocytes and translationally repressed and may indicate that LE females are viable although this is subject to more investigation (fig. S26, data S11). Core genes unique to HE females and not LE females or any other stage are enriched in apicoplast and mitochondrial components, as well as RNA associated processes consistent with ongoing active transcription (*48*) (fig. S26, data S11). A higher ratio of unspliced (pre-mRNAs) to spliced transcripts (mature mRNAs) in the HE as compared to the LE females is also supportive of more active transcription in the former (fig. S26). For the unique male (HE) core genes, enriched GO terms include cilium and microtubule components essential for male gamete function (*49*) (fig. S26, data S11). We further explore the biology of the LE gametocytes using long read sequencing below.

#### Striking differential expression between strains within the same host

To ensure the developmental atlas best represents the biology of natural infections, we integrated the parasites from the natural infection into the lab-based reference atlas (fig. S27). We then explored DE between the lab and donor parasites while controlling for stage and pseudotime, finding many DEGs (836-2838 depending on stage) (fig. S28-29, data S12). We report what we have found together with GSEA analysis but we cannot determine if these expression differences are driven by environment (e.g. lab culture vs natural host), genotype, or batch effects (fig. S29, data S12). Therefore, we focused on exploring DE between the different parasite strains observed within the natural infection (fig. S30, S31). We found between 6 and 319 DEGs between strains depending on life stage (fig. S31, data S13). For each pairwise strain comparison the number of DEGs was negatively correlated with the degree of genetic relatedness as demonstrated by the example of HE and LE females, where we had balanced field strain distribution in sufficient numbers (fig. S30, fig S32). HE females also had the highest gene expression levels giving the greatest power to detect differential expression (319 unique DEGs in total across pairwise comparisons between four strains) (Fig. 4C-E, fig. S31, data S13). Interesting genes showing genotype-specific patterns of expression among females include *Pfs16* and *Zip1*, both of which are implicated in parasite development within the mosquito (Fig. 4E, data S13) (*50*, *51*). Still focusing on female (HE), where we also had the most balanced distribution of lab strains in addition to that of donor strains (fig S30), we further investigated whether field strains within an infection display comparable levels of DE to *P .falciparum* lab adapted strains. The lab strains included a South American isolate (7G8) and a West African isolate (NF54), which were cultured independently and subjected to scRNAseq (fig. S31). Using subsamples of approximately 70 cells per strain, we find considerably more DE between field strains (SC5 vs SC7; 225 DEGs) than between the lab strains (7G8 vs NF54; 14 DEGs). Field parasites had more genes detected than lab parasites due to a higher depth of sequencing, nevertheless, restricting the analysis to only genes expressed in both lab and field parasites did not alter this finding (SC5 vs SC7; 132 DEGs, 7G8 vs NF54; 11 DEGs) (fig. S31). It is striking that an order of magnitude more genes show differential expression between strains within a single host than between lab strains originating from different continents and cultured separately.

#### Exploring stage-specific isoforms and exon usage within field parasites

We also carried out single cell IsoSeq on the parasites from this donor, resulting in 800 k unique long transcripts, most of which came from HE male and HE female cells, as well as SQANTI3-based isoforms classification of these transcripts (fig. S33, fig S34, data S14). Over 50 genes showed differential exon usage (DEU) between HE males and HE females, 37 of which are present within the lab sex stages comparison using similar thresholds, and we present a few of these examples (Fig. 4F, fig. S35, data S15). Visual exploration of DEU between the HE and LE females revealed that LE females tend to harbour longer transcripts while HE females are more variable in their transcript lengths, with some showing more 3’ degradation. This is also reflected in the length distribution of isoforms and proportion of full splice and incomplete splice match isoforms generated by SQANTI3 (fig. S36, data S14). Interestingly, 39 of 71 (55%) genes detected by long reads in LE females are among those known to be translationally repressed, compared to 121 of the 316 (38%) genes detected in HE females (fig. S36). This difference is more pronounced in short read data (fig. S36). The observations above may suggest that the HE gametocytes are undergoing active transcription, also supported by higher unspliced/spliced ratios relative to LE gametocytes as described earlier (fig. S26), and 3’ degradation to maintain RNA homeostasis. On the other hand, the LE gametocytes might be fully mature or dying female gametocytes that still retain the longer transcripts in messenger ribonucleoprotein (mRNP) particles, macromolecular complexes that harbour non-translating mRNAs bound by various RNA-binding proteins regulating translation, localization, and turnover (*22*, *52*). Given testing the transmissibility of LE gametocytes (which have never been seen in lab culture) will be impossible unless they are found to exist without HE gametocytes or as distinct strains in natural infections, further work to explore these unusual stages should entail longitudinal sampling from donors to better understand their dynamics, such as when they appear and how long they remain in circulation. Intriguingly, the LE parasites in this sample do not appear to be strain specific as they are observed for all strains that have any sexual stages.

### Integrated Malaria Cell Atlas as a reference for natural infections

While using single-cell approaches to study parasites derived from natural infections is in its infancy (*53*, *54*), unexpected observations underscore the importance of exploring natural infections at single cell resolution. Here, we find male and female stages not previously observed in lab culture. We also observe much greater transcriptional variation between genotypically distinct strains within the same host than between lab strains cultured in separate flasks. It remains to be determined whether differential expression and/or unusual life stages are driven by intra-host dynamics or other factors but it is clear that natural infections are revealing new biology that is likely to be relevant for persistence, pathology, and transmission. We have therefore integrated this first natural infection scRNAseq dataset into the reference atlas to create a combined resource that supports mapping of both donor and laboratory *P. falciparum* parasites together with a more complete view on interrogation of gene expression (Fig. 4G). The integrated object comprising 38,748 cells from laboratory and natural infections covering asexual and sexual differentiation is presented as a new chapter to the interactive Malaria Cell Atlas data resource (malariacellatlas.org). Single-cell approaches now enable us to study strain composition and expression behaviour including differential expression and isoform usage in natural infections, and have enormous potential in improving our understanding of malaria parasite biology and transmission in natural infections.

## Acknowledgments

This work was supported by a Medical Research Council research grant (MR/S02445X/1) to MKNL, which supports SD. This work was also supported by Wellcome core funding to the Wellcome Sanger Institute (Grant 206194/Z/17/Z), which supports MKNL and JCR. Part of the field work in Mali was supported by the BMGF (INV-001927) through RCA No 20_356 between Sanger and USTTB. We thank Matthew Jones and the staff of the Wellcome Sanger Institute Scientific Operations for their contribution to library preparation and sequencing, Yong Gu for help with running cellranger on raw data, David Serre, Brittany Hazzard, Lia Chappell, and Liz Tseng who advised on single-cell IsoSeq, Haynes Heaton who advised on strain analysis, and Virginia Howick, who assisted with one of the sampling days. We thank James Smith and the Core Software Team at Wellcome Sanger Institute for their contribution to the MCA web resource. We also thank the team that runs the clinic in Faladie, Mali for their contributions to sampling natural infections as well as the study population.

## Author contributions

Conceptualization: MKNL, AMT, AAD

Data Curation: SKD, JCR, JC, EF, AD

Formal Analysis: SKD, JCR, JC, EF

Methodology: AMT, SKD, JCR

Investigation: SKD, JCR, AD, EF, JC, DO

Visualisation: SKD, JCR, EF

Funding acquisition: MKNL, AAD, AMT

Project administration: MKNL, AAD

Supervision: MKNL, AMT, AAD

Writing – original draft: SKD, JCR, EF, JC

Writing – review & editing: SKD, JCR, EF, JC, AD, DO, AMT, AAD, MKNL

## Competing interests

Authors declare that they have no competing interests.

Data and materials availability: All raw sequencing data have been deposited in the European Nucleotide Archive at European Molecular Biology Laboratory European Bioinformatics Institute (www.ebi.ac.uk/ena/) under accession numbers PRJEB58790 and PRJEB55754. Stage and expression data across the life cycle stages profiled can be visualised at the Malaria Cell Atlas website (www.malariacellatlas.org). Metadata, raw counts and normalised counts for the cells are also available at the same website for download. Expression matrices, analysis code and supplementary files are archived on Zenodo (DOI:10.5281/zenodo.8139823).

## Supplementary Materials

Materials and Methods

Supplementary Text

Figs. S1 to S36

Tables S1 to S3

References (*55*–102)

Data S1 to S15

