## Supplementary Data data S4 for "A single cell atlas of sexual development in *Plasmodium falciparum*"

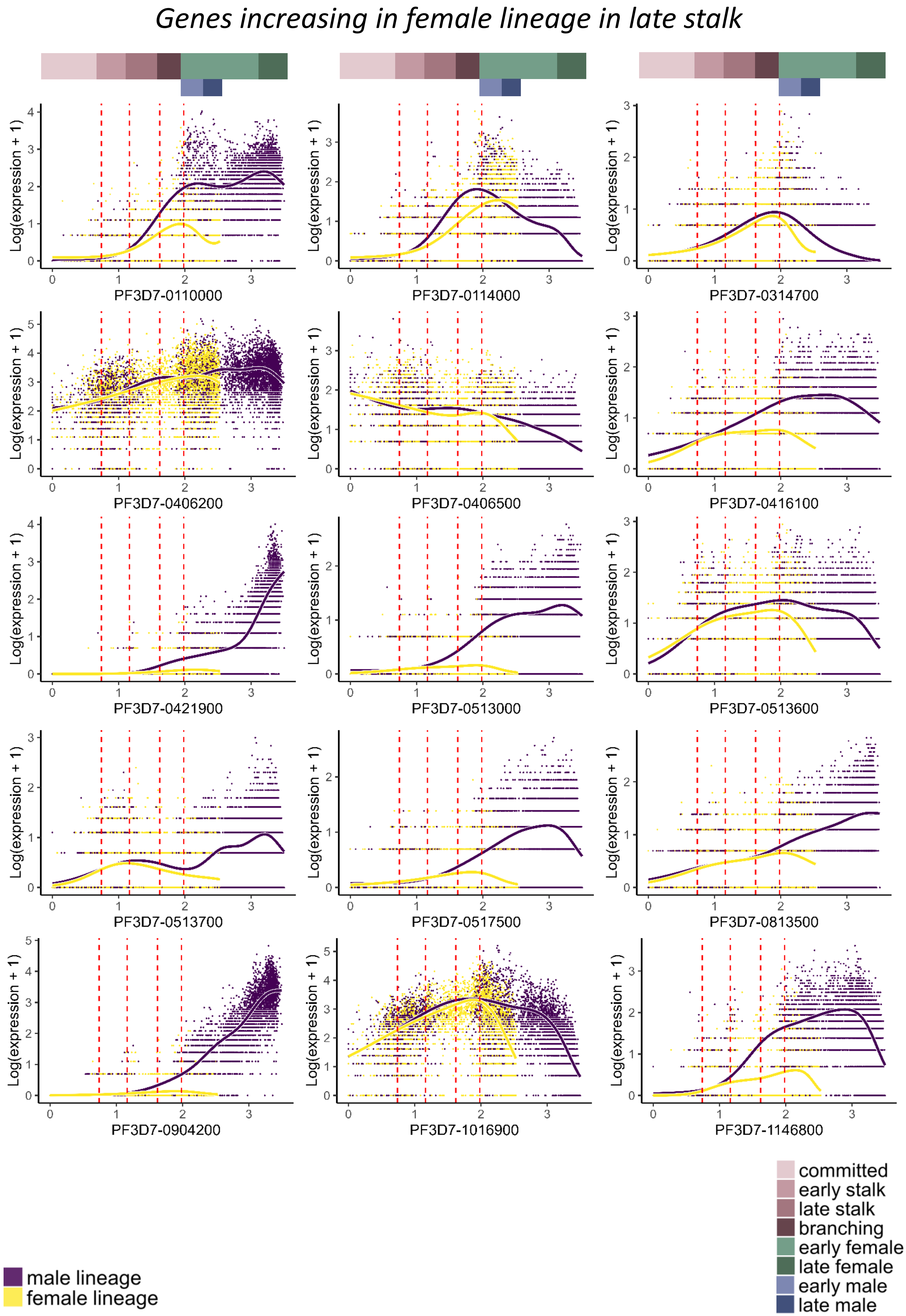

Genes increasing in female lineage in late stalk
Log(expression + 1)
PF3D7-0110000
PF3D7-0114000
PF3D7-0314700
PF3D7-0406200
PF3D7-0406500
PF3D7-0416100
PF3D7-0421900
PF3D7-0513000
PF3D7-0513600
PF3D7-0513700
PF3D7-0517500
PF3D7-0813500
PF3D7-0904200
PF3D7-1016900
PF3D7-1146800
male lineage
female lineage
committed
early stalk
late stalk
branching
early female
late female
early male
late male

### *Genes increasing in female lineage in late stalk*

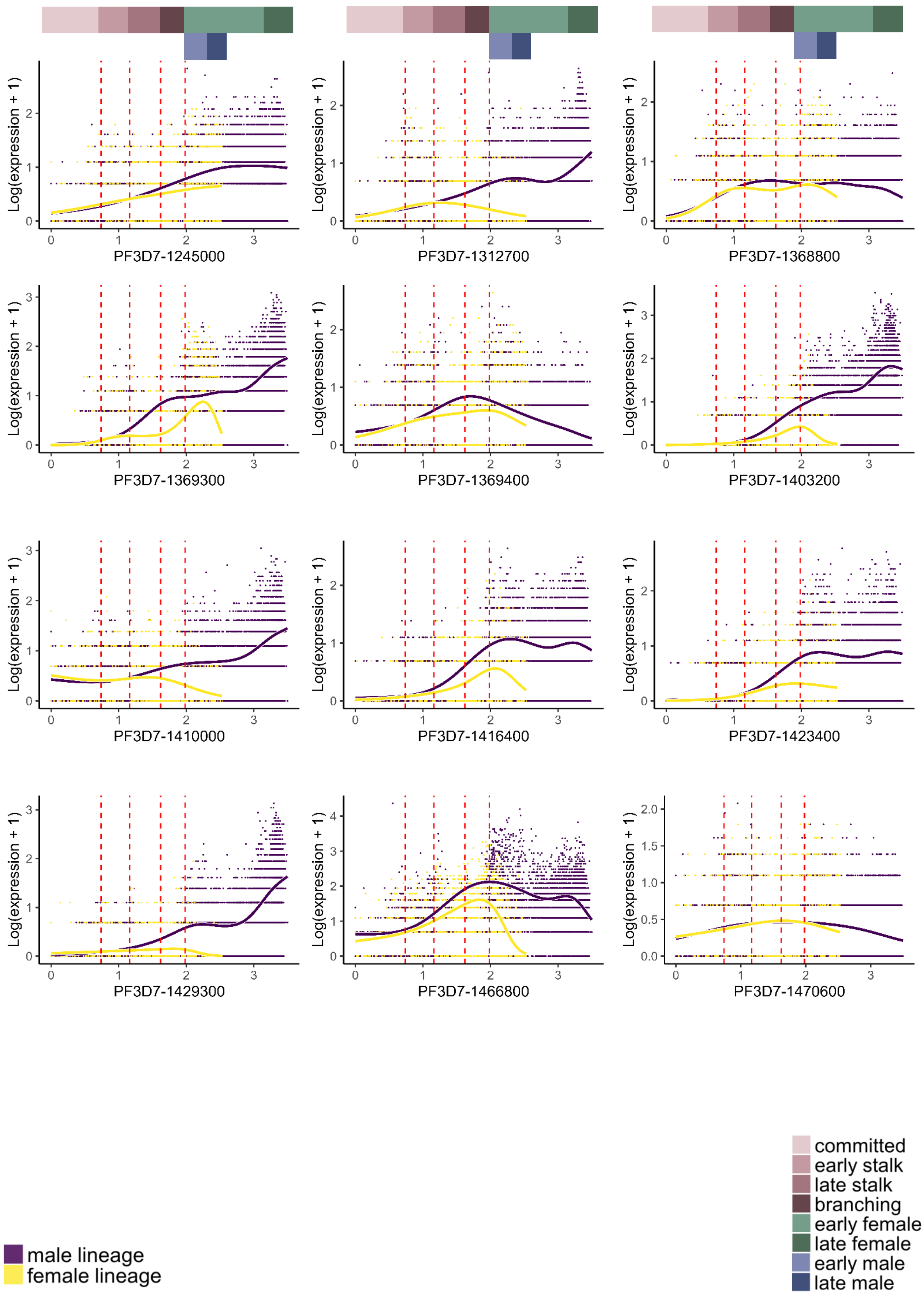

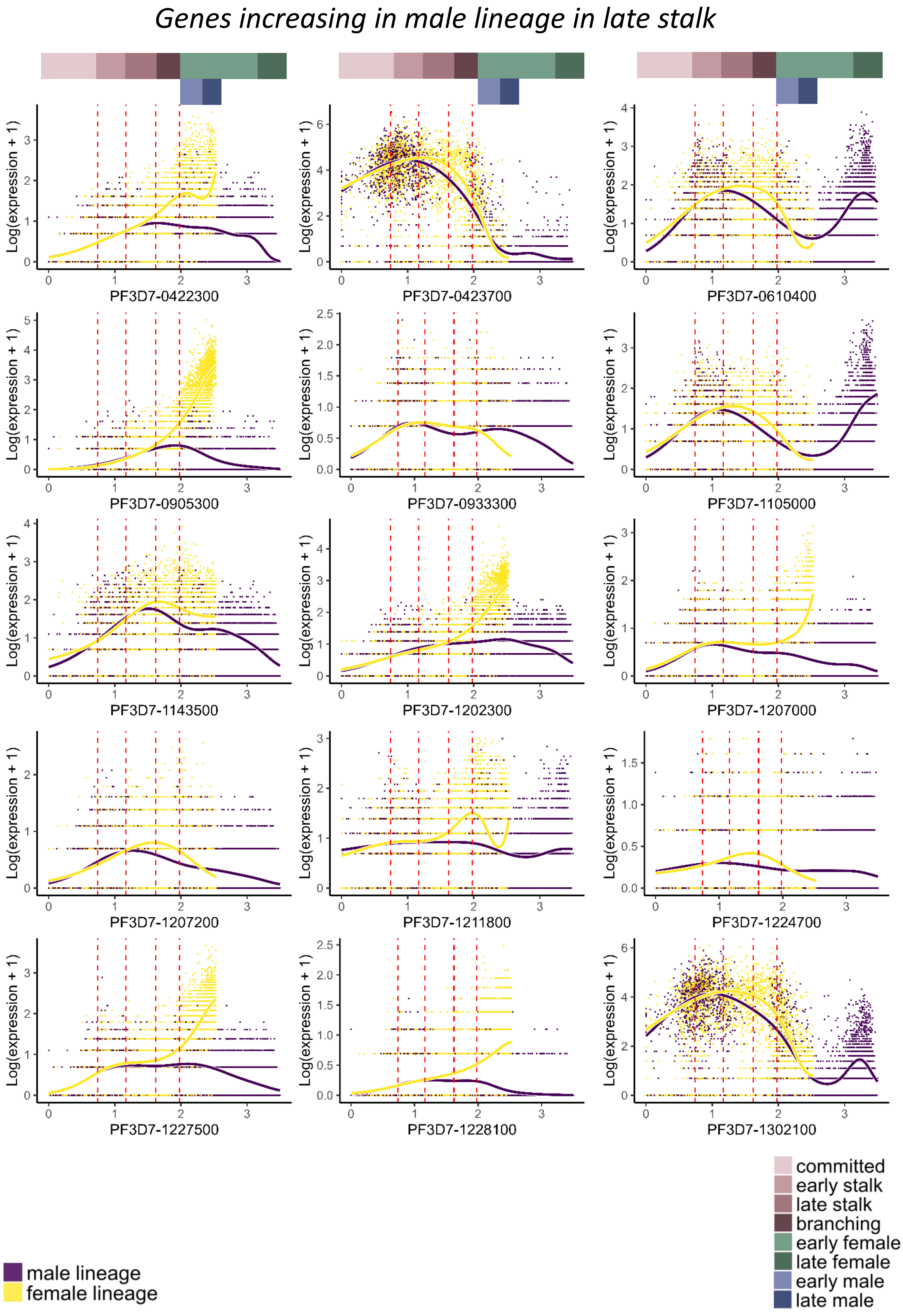
Genes increasing in male lineage in late stalk
Log(expression + 1)
PF3D7-0422300
PF3D7-0423700
PF3D7-0610400
PF3D7-0905300
PF3D7-0933300
PF3D7-1105000
PF3D7-1143500
PF3D7-1202300
PF3D7-1207000
PF3D7-1207200
PF3D7-1211800
PF3D7-1224700
PF3D7-1227500
PF3D7-1228100
PF3D7-1302100
male lineage
female lineage
committed
early stalk
late stalk
branching
early female
late female
early male
late male

*Genes increasing in male lineage in late stalk*

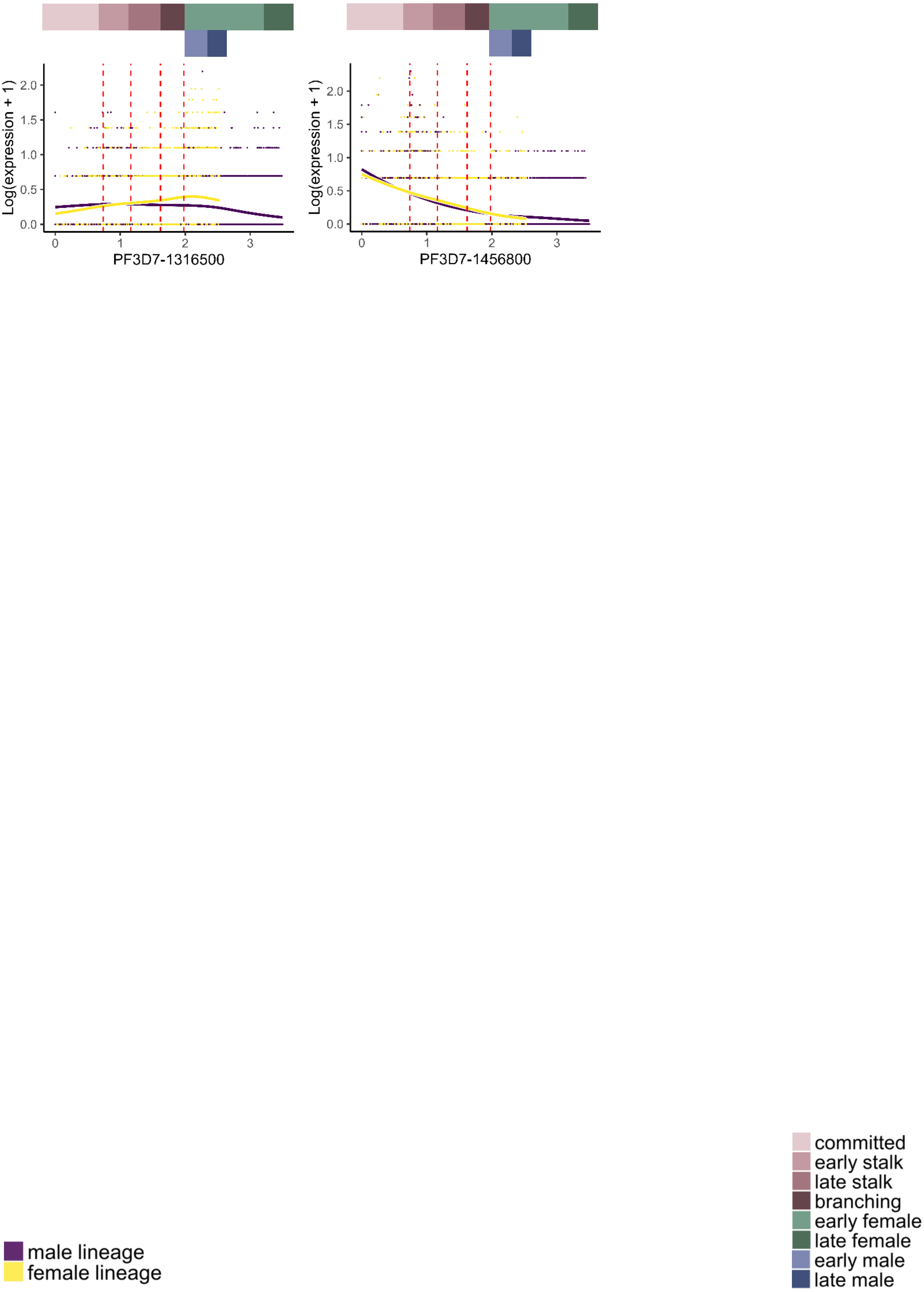
