## Supplementary Material for "A single cell atlas of sexual development in *Plasmodium falciparum*"

#### The PDF file includes:

Materials and Methods
Supplementary Text
Figs. S1 to S36
Tables S1 to S3
Captions for Data S1 to S15
References

#### Other Supplementary Materials for this manuscript include the following:

Data S1 to S15 (Excel format and PDF)

### Materials and Methods

#### *Plasmodium* culture and gametocyte induction

Asexual culturing of *P. falciparum* parasites (NF54) was done with O+ blood at 3% haematocrit in RPMI 1640 culture medium (Gibco) containing 5% Albumax and supplemented with 100x GlutaMAX™ (Gibco) (referred to as Complete Media or CM) in a mix containing 5% O2, 5% CO2, and 90% N2 and maintained at 37 ℃. Human O+ erythrocytes were supplied by NHS Blood and Transplant, Cambridge, UK. All samples were anonymized. Use of erythrocytes from human donors for *Plasmodium* culture was approved by the NHS Cambridgeshire 4 Research Ethics Committee (REC reference 15/EE/0253) and the Wellcome Sanger Institute Human Materials and Data Management Committee.

#### Gametocyte induction and sampling for scRNA-seq

Stage V gametocytes were obtained using a modified standard gametocyte culturing method (*55*). NF54 was thawed and revived in CM at 3% haematocrit. Revived parasites were maintained with a supplement of 5% pooled human serum, obtained locally in accordance with ethically approved protocols. Following gametocyte induction, parasites were sampled on multiple days from two sets of gametocyte inductions, where each induction time consisted of three separate parasite cultures synchronized 12h apart (T12-, T0 and T12+) to help cover more of the life cycle. For induction set 1, sorbitol synchronized parasites were seeded at 3% haematocrit and 1% parasitemia in T75 flasks for gametocyte induction and at 0.5% for asexual maintenance. For induction set 2, another induction flask was set up 2 days later from the asexual culture at 3% haematocrit and 1% parasitemia. Two-thirds of the media was changed everyday for three days until high parasitemia was reached and parasites were stressed. The complete media was changed daily thereafter. The two sets of flasks allowed sampling Day3 and Day5 on the same day along with the uninduced asexual culture (referred to as Day0), and Day10 sampling was done from the first set of induction flasks. Each of these sampling points consisted of a pool of the three T12-, T0 and T12+ flasks covering a 24 hour period in order to capture a wider developmental range. Heparin at 1 $\mu$g/ml was added to the flasks after Day3 to suppress asexual parasite growth. Parasitemia was estimated by Giemsa smears and RBC density was calculated on a hemocytometer. A sample from a second gametocyte induction experiment, consisting of parasites generated using the same gametocyte induction protocol as above and profiled on day 10, was added to the dataset to enrich the representation of the female and male gametocytes in our final dataset for the atlas (here labeled as D10b). This additional dataset consists of NF54 and 7G8 strain parasites that were grown and induced to gametocytogenesis separately prior to pooling them on Day 10 for 10X in a 1:1 ratio.

#### 10X loading, single-cell capture and Illumina library preparation

Cells were loaded directly from culture according to manufacturer's instructions using the Chromium Single Cell 3' GEM, Library, and Gel Bead Kit v3 (10X Genomics; PN-1000075) to recover 10,000 cells per inlet. Illumina libraries were prepared using the 10X Chromium Single Cell 3' Library kit (V3 chemistry) according to manufacturer's instructions. 14, 12, 12, 12, 12 cycles of cDNA amplification were done postGEM-RT for sample D0, D3, D5, D10a and D10b, respectively. 10 µl was used as input for Illumina library prep while the rest was used for PacBio library preparation. 10X input libraries were sequenced on a NovaSeq S4 PE100 with the following read protocol: 28, 8, 0, 91 at 6-plex.

#### QC: Filtering of doublets and low quality cells

Across the 5 samples, low quality cells and empty droplets were sorted out by removing all droplets containing less than 1000 total UMIs and 300 genes (fig. S1). Stage doublets were removed by applying DoubletFinder and scrublet with varying parameters and thresholding after visual inspection (fig. S1) (*56*, *57*). Souporcell (*44*) was used to assign a strain to each cell in the D10b sample based on single nucleotide polymorphisms (SNPs) present in the read data, matching variants to those in the genome of 3D7 (a clone of NF54) and 7G8 from the Pf6K data set (*58*). This allowed us to also remove putative inter-strain doublets following annotation.

#### Mapping to single cell and bulk RNAseq datasets

Smart-Seq2 *Pf* single-cell datasets were obtained from Zenodo (https://doi.org/10.5281/zenodo.2843883) and mapped with scmap to the current dataset (fig. S2, S3) (*53*). Four bulk RNAseq datasets covering the entire intraerythrocytic development were taken directly from the publications (*22*, *59*–*61*). Based on pseudotime values, we created 48 mini-bulks of approximately 540 cells each across the asexual cycle. The stalk, male, and female lineages were divided into 33 total pseudobulks each comprising 300 cells. Similarity of the pseudobulks to bulk RNAseq time points was calculated using Pearson correlation (fig. S2, S3).

#### Annotation

Asexual cells were labeled according to their correlation with *P. falciparum* bulk RNAseq samples from the asexual cycle and mapping to SS2 single-cell RNAseq data (*59*, *60*, *62*) (fig. S2). Early gametocytes (early sex bulk 1 & 2) are labeled based on Louvain clustering and correlation with bulk RNAseq time points after gametocyte induction (*61*) (fig. S2, S3). Late-stage gametocytes (female bulk 1 & 2, male bulk 1 & 2) were assigned based on their correlation with FACS-sorted late-stage gametocytes and Louvain clustering (*22*) (fig. S2, S3). Louvain clustering was performed using the Seurat package with a resolution of 0.2 (fig. S2) (*63*). For pseudotime analysis, the Louvain clusters were used as input to Slingshot (fig. S2) (*64*). Analysis scripts and data files associated with annotation are archived on Zenodo (DOI:10.5281/zenodo.8139823).

#### Assigning pseudotime to male and female lineages

Cells were ordered along pseudotime using slingshot version 2.2.1 (*64*). Sexual stage cells assigned as committed were chosen as the starting cluster with start.clus. To obtain pseudotime along male and female lineages, terminal clusters in the male and female stages (male bulk 2, female bulk 2 in fig. S2) were chosen, respectively, as the terminating clusters with end.clus. UMAP embeddings were used as input. Sub-stages (e.g. committed, early stalk, late stalk, branching) were assigned to obtain an equal proportion of early and late developmental stages within the stalk, the male, and the female gametocytes. Final stage assignments of the cells are as follows: early ring, late ring, early trophozoite, late trophozoite, early schizont, late schizont, gametocyte (developing) [committed, early stalk, late stalk, branching], gametocyte (female) [early female, late female], and gametocyte (male) [early male, late male)

#### Modeling gene expression over pseudotime for sexual stages

Gene expression along pseudotime within the sexual stages was estimated using tradeSeq version 1.8.0 (*65*), which uses a negative binomial generalized additive model (NB-GAM) to model expression values across genes in multiple lineages. Raw (non-normalized, non-logged) counts were supplied to tradeSeq along with pseudotime values and cell weights generated by slingshot. *evaluateK* function was used on a sub-sample of 200 genes to obtain an estimation of the optimal number of knots to provide as input to *fitGAM* function. The *fitGAM* command was run on the 8958 sexual stage parasites using nknots of 8.

#### Gene expression differences over pseudotime using tradeSeq

tradeSeq's *startVsend* test was used to perform the statistical test to check for DE between starting and end points of the desired lineage. Candidates with a log fold change of 1 and a minimum waldstat value of 60/100 were considered for further analysis and heatmaps.

#### Gene graph and GRN inference

The cell x gene mRNA expression matrix was filtered to include only genes with more than 200 overall counts. This matrix was transposed to a gene x cell matrix containing 5087 genes, which was then used to calculate a two-dimensional embedding using the python library

UMAP (*66*). On the basis of this embedding we classified all genes into 15 clusters using the *scikit-learn* implementation of the k-means algorithm (*67*, *68*). To investigate the gene expression of the gene clusters, cells were ordered according to pseudotime and assembled into 300-cell pseudobulks. A heatmap was then generated displaying the gene expression of the pseudobulks ordered by cell identity in the x - and cluster identity in the y-direction using the ComplexHeatmap package (*69*).

#### Gene regulatory network inference

A machine learning approach was employed to train Gene Regulatory Networks (GRN) using the GENIE3 algorithm (*31*). The input for this training process consisted of pseudo temporal assembled pseudobulks generated from log normalized counts obtained via the Seurat *NormalizeCounts* function (*70*). The pseudobulk sizes used in this study consisted of 150, 300 450 or 540 cells, and two distinct inference algorithms, Extra-Trees and Random Forest, available in GENIE3 were utilized to improve robustness. To further refine our results, analysis was restricted to a pre-selected list of transcription factors from the ApicoTFdb database (*30*). Finally, the weights of 8 GRNs obtained from the four pseudobulk sizes and both inference algorithms were merged to provide robust regulatory relationships. The regulators were ranked after the total number of connections in the top 1.5%.

#### AT% calculation of promoter regions

Sequences of the region 500 bp 5' upstream of the gene body or up to the closest gene were extracted and those genes with at least 400 bp of upstream sequence were considered for plotting AT% across each expression module defined in Fig 1D and supplementary figures S5-7.

#### Single cell IsoSeq library preparation

To prepare single-cell IsoSeq Libraries, 25 µl of cDNA from the postGEM-RT cleanup of the 10X v3.1 3’ runs were amplified further with NEBNext® High-Fidelity 2X PCR Master Mix and cDNA Primers from the Chromium Next GEM Single Cell 3' GEM, Library & Gel Bead Kit v3.1, 16 rxns (PN-2000089). 14, 12, 12, 12, 12 cycles of cDNA amplification were done postGEM-RT and used as input for Illumina library prep for sample D0, D3, D5, D10a and D10b, respectively. Additional 4,3,3,3,6 cycles of amplification were done on the 25 µl of the remaining cDNA to reach the recommended minimum input of 160 ng for IsoSeq library prep, respectively. PacBio’s guidelines were followed to purify this amplified cDNA and generate Single-Cell IsoSeq™ Libraries Using SMRTbell® Express Template Prep Kit 2.0. Each library was sequenced on its own Sequel II SMRTcell.

#### Single cell IsoSeq analysis workflow for lab datasets

Preliminary analysis of the sequencing output from the PacBio reads was done by following the guidelines as recommended here (https://github.com/Magdoll/cDNA_Cupcake/wiki/Iso-Seq-Single-Cell-Analysis:-Recommended-Analysis-Guidelines) to generate a full-length non-concatemer fasta file for each of the samples. The isoseq3 *dedup* step was done on each of the five samples to perform PCR deduplication via clustering by UMI and cell barcodes, generating one consensus sequence per founder molecule.

To generate a Seurat-compatible input encompassing all the cells, deduplicated fasta files from days 0, 3, 5, 10a, and 10b were merged and subjected to the workflow according to the recommended guidelines as above, with the following change in parameters: -c 0.99 -i 0.98 for collapse_isoforms_by_sam.py, filtered to a minimum of 25 FL (Full Length) read counts, mapping with minimap2 using a maximum intron size (-G) of 5000, 0.9 for intrapriming parameter (-a) for sqanti3_RulesFilter.py. A Seurat-compatible input was generated (provided at Zenodo DOI:10.5281/zenodo.8139823) from these SQANTI3 isoforms using *make_seurat_input.py*, which was loaded into Seurat. Metadata was added to the object using the cell barcode assignments from short read data, thereby linking the demultiplexed long reads to the life cycle stages. A centered log-ratio (CLR) normalization was done instead of *LogNormalization* due to the low number of features per cell in IsoSeq data, followed by *ScaleData* and *RunPCA* using all the genes for *VariableFeatures*. Cells with more than 20 reads and genes are used for UMAP dimensionality reduction visualization, presented in Fig 3.

For generating stage-specific and chained isoforms annotations, deduplicated fasta files from days 0, 3, 5, 10a and 10b were merged and stage-specific long read files were generated by deconvolving reads for each life cycle stage (ring, trophozoite, schizont, gametocyte (developing), gametocyte (female) and gametocyte (male)) using single cell barcodes from short read data. Reads from each stage were separately subjected to the workflow according to the recommended guidelines as above, with the following change in parameters: -c 0.99 -i 0.98 for *collapse_isoforms_by_sam.py*, filtered to a minimum of 10 FL read counts, mapping with *minimap2* using a maximum intron size (-G) of 5000, 0.9 for intrapriming parameter (-a) for *sqanti3_RulesFilter.py*.

Across the stages, SQANTI3 generated 59266 transcripts, with the minimum in early gametocytes (2815) and maximum in trophozoite stages (18696). These isoforms were further filtered to remove those with >90% of adenines downstream of the transcription end site to remove transcripts arising from intrapriming (a higher cutoff of 90% AT was used, compared to the recommended 60-70%, due to high AT richness of *P. falciparum*), those with non-canonical splicing junctions, and transcripts with single exons (by using *sqanti3_filter.py*), while retaining transcripts containing full splice matches and novel isoforms.

Filtered isoforms (totalling 18945) from each stage were chained using chain_samples.py to get 13540 non-redundant, collapsed transcripts. These 13540 isoforms were classified among 2724 annotated genes and 748 novel genes, with a mean length of ~1407bp and 938bp respectively, compared to a mean length of 3022bp for current annotated transcripts on PlasmoDB v59. As no 5' end enrichment was performed for these samples, 5' ends were compared to the isoforms identified previously (*71*). Classification of the isoforms following chaining and the annotation file of these isoforms in GTF format is provided in data S7.

The classification of all the SQANTI3 generated transcripts pre-filtering is provided at Zenodo (DOI:10.5281/zenodo.8139823), along with the genome annotation of filtered SQANTI3 identified isoforms for each stage in GFF format. Annotation of the chained isoforms in GTF format merged with *Plasmodium falciparum* 3D7 reference annotation from PlasmoDB v52, is provided at the same repository. In these annotation files, field 2 contains the isoform identifier (PacBio for isoforms identified in this study, and VEuPathDB for annotated genes from PlasmoDB v52). Field 9 of the GTF also includes a color code for the SQANTI3 identified isoforms from this study, where they are colored according to the level of support from long-read RNA-seq data (darker, more support). The color is based on the abundance values calculated from the counts for each isoform using cupcake's *color_bed12_post_sqanti.py* script.

*P. falciparum* was not thought to have much isoform diversity, with only 67 of the annotated 5791 genes characterized as having more than one isoform on PlasmoDB (v56). However, recent RNA-Seq studies exploring 5' capped transcripts in asexual intraerythrocytic developmental stages suggest that the true isoform diversity is much higher (*71*–*73*). The most recent study identified 12,495 novel isoforms among bulk asexual stages alone (*71*).

#### Differential exon usage using DEXseq

A combined BAM file, made by merging the deduplicated reads from across the five samples, was created to include only those with cell barcodes present in the short read data. This BAM was then further split stage-wise using cell barcode based on the stage assignments of each cell from the short read data, for differential exon usage (DEU) analysis and IGV (Integrative Genomics Viewer) visualizations. Each of the stage-wise BAM files were split into two, by random sampling to obtain the two 'bulk' replicates needed for DEXSeq (package version: 1.43.0) to assess differential exon usage between the following conditions: i) early trophozoite vs early gametocyte (committed, stalk), ii) male (early male, late male) vs female (early female, late female)  iii) female (early female, late female) vs asexual (ring, trophozoite, schizont), iv) male (early male, late male) vs asexual (ring, trophozoite, schizont), v) ring vs trophozoite, vi) trophozoite vs schizont, vii) ring vs schizont.

Guidelines from DEXseq vignette were followed to generate count matrices and annotation file (https://bioconductor.org/packages/devel/bioc/vignettes/DEXSeq/inst/doc/DEXSeq.html).

Pairwise comparison was done for the above sets of stage clusters using the default parameters with a simple design of '~ sample + exon + condition:exon'. From the DEXSeqResults object, we asked for exonic regions with a adjusted p-value of < 0.05 and fold change of 1 for i, adjuster p-value of < e-20 and fold change of 2 for ii-v, and adjusted p-value of < e-10 and fold change of 1 for v-vii.

#### Isolation of *Plasmodium* cells from donor blood for 10X scRNA-seq

A detailed protocol is available (*74*). Briefly, a venous blood sample from the donor (Female, age 10 years, Parasitemia 3200/µl, Gametocytemia 104/µl) was collected in a CPDA (citric acid, monobasic sodium phosphate, dextrose, adenine) tube and mixed thoroughly. 3 mL of this blood was resuspended in 5 mL of cold suspended animation (SA) buffer (10 mM Tris-HCL, 166 mM NaCl, 10 mM Glucose, pH 7.37) and passed through a pre-wetted MACS column, and then eluted in ~60 µL SA buffer to obtain gametocytes. 50 µL of the elute was re-pelleted and resuspended in 10 µL SA buffer to concentrate gametocytes for 10X loading. The MACS flowthrough was passed through a Plasmodipur™ filter (Europroxima 8011FILTER) to remove leukocytes, and then treated with Streptolysin O from Streptococcus Pyrogenes (S5265-25KU, from Sigma Aldrich) activated with DTT (P2325, from Life Technologies Ltd) to remove uninfected RBCs (*75*) and obtain RBCs containing circulating asexual stages. Hemocytometers were used to obtain the number of cells and Giemsa smears were made to estimate the proportion of infected RBCs. The sexual and asexual fractions were pooled together for a single 10X run targeting a recovery of 10000 cells according to manufacturer's instructions using the Chromium Single Cell 3' GEM, Library, and Gel Bead Kit v3 (10X Genomics; PN-1000075). Lysis and MACS washes did not remove all the uninfected RBCs - Gametocyte fraction had 7.5% parasites while the SLO fraction had 52% parasites.

#### Single cell Illumina and Iso-Seq library preparation for field sample

Illumina libraries were prepared as described for lab datasets. 14 cycles of cDNA amplification were done postGEM-RT and 10 µl of cDNA was used for preparing Illumina libraries, which were sequenced on a NovaSeq S4 PE100 (1/7th of a lane) with the following read protocol: 28, 8, 0, 91. Single-cell IsoSeq Libraries were prepared as described above for lab samples. An additional 11 cycles of amplification were done on the 25 µl of the remaining cDNA and the library was sequenced on a single Sequel II SMRTcell.

#### Souporcell genotype deconvolution of donor strains and genotype quality control (QC)

To determine the number of strains in the donor sample (hereafter "donor" or "field"), souporcell was run on the filtered cells from cellranger as previously described (*44*). Since souporcell requires the number of strains (k) to be specified, we assumed a range of 1 to 10 strains in the sample based on previous reports on the complexity of infection (COI) in endemic areas (*76*) (*77*) . We then ran souporcell for each of the 10 assumptions and plotted the log-likelihood for all of the iterations (fig. S20). The k just before the log-likelihood plateaus is considered as the most likely number of strains/genotypes in a sample. Because this inflection happened over a range of ks (fig. S20), we considered three values (6,7 and 8) to be likely and proceeded with the higher value (k=8) classification to allow utmost precision. All the cells that were classified as 'Negative' based on souporcell k=6, k=7, and k=8 classifications were removed. 'Negative' cells are possibly barcoded droplets with uncertain assignment that contain ambient RNA and fractions of cells that escaped the cellranger QC. Inter-strain doublets based on souporcell k=6, k=7, and k=8 were retained for later use as ground-truth doublets to facilitate the removal of inter-stage doublets at which point they were also removed as described in the section below. Genotype PCA clustering of the parasites using the k=8 classification was used to visualize the cells (fig. S22). A second round of independent genotype assessment was then done as described further below.

#### Gene expression based clustering and QC of donor parasites

After souporcell strain assignment, we clustered the cells based on short read data using the Seurat v4 (*63*) standard pipeline to get a view of the gene expression clusters. We then proceeded with QC based on gene expression profiles. We removed cells with fewer than 300 genes and 700 UMIs (unique transcripts) (fig. S20) presumed to be of poor quality. We also removed cells where the proportion of transcripts from the mitochondrial genome was higher than 0.45% (fig. S20) presumed to be dying cells. To improve doublet detection, we ran SoupX (*78*) to first correct the UMI counts per cell by adjusting for the soup fraction, i.e ambient RNA from lysed cells that end up in the cell suspension and is included as part of the transcriptome of each cell. We then clustered the cells using Seurat SCTransform (*79*) pipeline which has more resolution in identifying distinct clusters. We then used inter-strain doublets identified above as a ground-truth in the identification of inter-stage doublets within the DoubletFinder (*56*) pipeline (fig. S20). We also used scrublet (*57*) to identify inter-stage doublets without prior specification of a ground-truth set (fig. S20). We removed all the doublets identified by these methods retaining a QCd set of parasites which we reclustered using the standard Seurat v4 (63) pipeline and proceeded with lifecycle stage annotation.

#### Lifecycle stage label assignment of donor parasites

To transfer labels from the MCA V3 lab reference dataset to the query donor parasites, we used scmapCell which clusters reference cells into subcentroids using kmeans and then projects query cells onto this using k-nearest-neighbors (*80*). Donor parasites where the three closest neighboring reference cells had the same stage label, each with a cosine similarity score of >0.4 to the donor parasite, were assigned the label of these reference cells (fig. S21). For those donor parasites where fewer than three of the closest reference parasites had the same stage, we assigned the top stage label as long as this had a cosine similarity score of >0.4 with the donor parasite and all the top 10 most closest reference parasites were similar cell types (fig. S21). We finally assessed expression of known asexual and sexual stage markers to assess concordance with the label transfer method (fig. S21). We retained only those cells that were confidently annotated for the further downstream analysis.

#### Custom pipeline for stringent pseudobulk genotyping for verification of souporcell assignments and relatedness estimations

Once the QC'd parasites had confident stage labels and initial souporcell k=8 genotype assessment, we then did a more stringent second round of strain assessment. The 8 souporcell clusters (SC) were further classified into coarse-stages (asexual, gametocyte) (fig. S22). Strain+coarse-stage clusters with fewer than 30 cells were not considered. The remaining strain+coarse-stage pseudo-bulk clusters were treated as independent samples/groups. The reads from these independent samples were mapped using minimap (*81*) to the *P. falciparum* reference. SNPs differentiating the samples were then called using freebayes (*82*) specifying a ploidy of 1. In the freebayes run, all indels, multi-nucleotide polymorphisms and complex alleles were removed before variant calling. We also only included alleles if the supporting base quality was at least 20, if at least 20% of observations supported the alternate allele in at least one sample, if it was supported by a coverage depth of at least 6, and if it was supported by reads where the alignment mapping quality was greater than 30. All the other settings were left as default. BCFtools was then used to retain only the biallelic SNPs.

For the SNPs identified by freebayes above, we used vartrix (*83*) to count the number of deduplicated transcripts supporting alternative and reference alleles per sample by leveraging 10X UMI barcodes to remove PCR duplicates. Here alleles were counted only if the supporting reads had a mapping quality of at least 30. All the other parameters were left as default.

We then performed further QC on the called SNPs in R. The VCF for biallelic SNPs called by freebayes across the pseudobulk clusters was read into R using seqarray (*84*). We retained SNPs with greater than 75 genotyping QC and with a depth across all the samples of greater than 40. We termed SNPs removed in this step as 'Low quality in later sections in the paper (fig. S23). We then removed 'Mt' (mitochondrial) SNPs (fig. S23). We retained only those SNPs that were also present in the QCd (filter == 'PASS') variants identified from Malian samples included the pf7 *P.falciparum* dataset (*85*). Those removed in this step are labeled as 'Absent in Pf7' in further descriptions below (fig. S23).

We further used the vartrix output for sample specific QC. This was to avoid evaluating SNPs that may have low quality in particular pseudobulk samples/groups. Here we did not remove entire SNPs/variants but only removed the called genotypes in the affected groups. We retained only those genotypes for which either the alternate or reference allele was supported by at least 8 unique transcripts / unique molecular identifiers (UMIs) in the respective sample. Those removed in this step were termed 'Low UMI support' (fig. S23). We then removed SNP calls where there were more than 15 unique transcripts supporting both the alternate and reference alleles, termed as 'Non-distinct' (fig. S23). We then removed calls where either the alternate or reference allele was supported by more than 8 unique transcripts but the difference between the reference and alternate unique transcripts was less than 8, termed as 'Noisy' (fig. S23). We then removed variants with discordance between the vartrix allele counts and freebayes calls. These included 'Alt inverted', those where freebayes called the SNP as being of the alternate allele but vartrix had more UMIs supporting the reference allele and 'Ref inverted' calls where freebayes called the SNP as being of the reference allele but vartrix had UMIs counts supporting the alternate allele (fig. S23). This resulted in a clean set of genotypes that were used for the rest of the analysis (fig. S23).

Using the QCd genotypes we then assessed genotype identity between the samples using snpRelate identity by state (IBS) (*86*) metrics (fig. S24). Samples (asexual and gametocyte pseudobulk groups) with IBS greater than 0.95 were grouped together and classified to be of one strain (fig. S24). We then re-genotyped these collapsed strain pseudobulk groups leveraging on the increased number of cells per group to identify more SNPs. This marked the end of strain classification.

#### Relatedness between donor strains

We performed manual pairwise comparison of alleles between the different strains and plotted sharedness across 100kb windows of the genome (fig. S25). We corroborated the consequent results using hmmIBD (*87*) which also gave values on degree of relatedness between the different strains (fig. S25, S26). We compared the IBD results from the SNPs that passed all the QC steps above to the IBD results from SNPs that passed only those QC filters applied to exclude variants indiscriminate of pseudobulk group i.e removing 'Low quality', 'Mt' and ‘Absent in Pf7’ variants but retaining ‘Non-distinct’, ‘Noisy’, ‘Alt inverted’ and ‘Ref inverted’ genotype calls. This was to assess the effect of the stringent genotyping workflow.

#### Life cycle stage gene overlap in donor parasites

We considered a gene to be present in a stage if it was detected in at least 25% of the parasites of that stage. We then assessed overlap of these genes between all the stages visualizing these using UpSet plots. The genes that were characteristic of particular stages of interest were then subjected to GO enrichment analysis using the clusterprofiler package (*88*) and the latest GO annotation file release from PlasmoDB (*89*). All the ontologies; cellular components, molecular function and biological processes were considered separately and only those terms with an adjusted *p*-value of less than 0.05 were considered to be enriched in the particular list of genes under investigation.

#### Integration of donor and lab parasites, strain DE and GSEA

We used the standard Seurat integration as described in (*90*) to integrate the field and lab datasets (fig. S26). We then rescaled the integrated data and recomputed the PCs and UMAP using the same parameters as those used for the MCA V3 lab reference dataset. We calculate pseudotime along the integrated object using slingshot (*64*). For each field stage that clustered separately in the integrated UMAP we subsetted the part of the UMAP where the parasites from this stage interspersed with lab parasites for differential expression (DE) analysis (fig. S26). We did this by using the minimum and maximum pseudotimes of field parasites within the particular stage to define the boundaries and subsetting all the field and lab parasites within 98.5% of this range. We then categorized the subsetted parasites into chunks based on pseudotime for downsampling. For the field versus lab DE analysis we downsampled the lab parasites to be equal to the field parasites within each pseudotime chunk. For the strain DE, we downsampled each of the 7G8 and NF54 strains independently to be equal to the most abundant field strain within each pseudotime chunk. Field parasites were not downsampled. (fig. S26). We used both seurat MAST (*91*) and tradeSeq (*65*) to perform DE between the donor and lab parasites,between the parasite strains within the donor sample and between lab strains 7G8 and NF54 from D10b. We did the analysis within each stage while adjusting for pseudotime and only considered genes that were present in at least 10% of the parasites in one of the groups being compared. Genes that attained a significance threshold of adjusted *p*-value < 0.05 in seurat MAST and waldStat >10 in tradeSeq were retained as significant. To perform GSEA of the differentially expressed genes, we reran Seurat MAST DE with all the genes detected in the parasite groups under comparison and used the resultant log fold changes to rank the genes. We then used this ranked list together with the latest GO annotation file from PlasmoDB to compute GSEA using the fgsea (*92*) method within clusterprofiler package (*88*).

#### Single cell IsoSeq analysis workflow for donor parasites

Long read output from the field sample was processed as described for the lab datasets, with two minor changes. A lower threshold of 10 FL read counts was used for generating isoforms encompassing all cells, and a threshold of 5 FL counts was used for generating stage-specific isoforms. Cells with more than 5 reads and 5 genes are used for UMAP dimensionality reduction visualization, presented in fig. S33.

For differential exon usage (DEU) analysis and IGV (Integrative Genomics Viewer) visualizations, stage-specific bam files were generated as described above for lab datasets. DEXseq (*41*) was then applied to the long reads to identify differentially expressed exons between all pairwise comparisons of the field stages with sufficient number of parasites. Genes with an adjusted *p*-value of less than 0.01 and log fold-change of more than 1 were considered as having differentially expressed exons. These were visualized by plotting BAM coverage plots on IGV split by stages.

The classification of all the SQANTI3 generated transcripts pre-filtering is provided at Zenodo (DOI:10.5281/zenodo.8139823), along with the genome annotation files of filtered SQANTI3-identified isoforms for each stage. Annotation of the chained isoforms merged with *Plasmodium falciparum* 3D7 reference annotation from PlasmoDB v52, is provided at the same repository. In these annotation files, field 2 contains the isoform identifier (PacBio for isoforms identified in this study, and VEuPathDB for annotated genes from PlasmoDB v52). Field 9 of the GTF also includes a color code for the SQANTI3-identified isoforms from this study, where they are colored according to the level of support from long-read RNA-seq data (darker, more support). The color is based on the abundance values calculated from the counts for each isoform using cupcake's *color_bed12_post_sqanti.py* script.

### Supplementary Text

#### Technical challenges with 10x single cell IsoSeq data

It is difficult or impossible to assign reads to distinct isoforms using 10X 3' short read data alone. Full-length isoform sequencing of 10X generated single cell cDNA using PacBio IsoSeq is a promising approach to explore isoform diversity at single cell resolution. Under ideal circumstances, full length cDNA should result in an accurate sequence of the 5'- and 3'-UTRs and identify novel splicing events. In this study, we investigated whether cell type assignments derived from single-cell transcriptomes can enrich our understanding of isoform diversity across the parasite's intraerythrocytic development, enabling understanding of true isoform diversity present in each cell type.

Long reads were classified with SQANTI3 (*93*), to compare and describe the isoforms based on how they match the current gene annotation available from PlasmoDB (figs. S10 and S11). Stage labels from short read data allowed us to do this for each cell type (ring, trophozoite, schizont, early, female and male gametocytes) (fig. S10). Following initial isoform identification within each cell type by SQANTI3, prior to filtering, most isoforms (~69%) are single-exonic, with less than 1% of them being an exact match to the reference annotation (fig. S11). While a fraction of these reads are likely real novel isoforms, it is likely that majority of them are fragments arising from various artifacts as described below. We also observed isoforms (< 5%) with non-canonical splice junctions, which may be real splice events, though we can't rule out experimental or technical artifacts. Among the 13540 filtered isoforms chained from all the cell types, full splice matches (FSM) and incomplete splice matches (ISM) are the most common structural categories accounting for 4828 (36%) and 3311 (24%) of isoforms, respectively. FSM describes isoforms that match the splicing sites and exons in the reference annotation with allowance for different 5' and 3' ends. Only 64 (1.3%) of the 4828 FSM isoforms matched the existing gene annotations with regards to splice sites and 5'/3' ends, with the rest displaying alternative 3' ends (15%), alternative 5' ends (6.3%) or both (77.4%).

Several recent studies using long read sequencing were instrumental in attempting to improve the *P. falciparum* annotation, especially with regards to updating UTRs in the current PlasmoDB annotation (*71*, *72*, *94*, *95*). In spite of this, we still achieve very few (51) perfect matches between the full length cDNA sequenced here and the annotation. While low reference match and major proportion of alternate end isoforms could indicate poor UTR annotation, there are other caveats to consider as well in our IsoSeq data. These caveats could result in false assignment of isoforms into the various structural categories that SQANTI3 employs. These artifacts could influence differential isoform expression analysis and differential exon usage analysis.

Firstly, the transcript length distribution we observed is low compared to the annotated gene models, which can be apparent from the mean length of raw reads fed into SQANTI3 (fig. S10). This discrepancy could be due to lower processivity of Chromium 10X reverse transcriptase or driven by artifacts caused by the template switching oligo. Synthesis of complete transcripts from long genes may be disadvantaged, giving rise to isoforms mistakenly identified as ISMs or FSMs with alternate ends. While it is possible that some ISM isoforms are in fact real isoforms, as in the case of the female-specific isoform seen in md1 (Fig. 3E), others might be partial fragments arising from incomplete reverse transcription or mRNA decay. Similarly, in addition to PCR or sequencing artifacts generated within a single cell, partially degraded molecules and lack of 5' capping can result in misclassification of isoforms into the various structural categories. As no 5' end enrichment was performed for these samples, 5’ ends were compared to the isoforms identified previously (*71*) revealing ~51% of FSMs fall within 100bp of their TSS. In addition to the lack of 5’ cap enrichment, this could reflect fragmented libraries, poor annotation of UTR ends in the current annotation, and/or the additional representation of isoforms not previously found due to sexual stages present here.

The likely fragmented nature of the data makes differential isoform expression analysis at single-cell level challenging, which is why we focused on exploring exon usage at a cell type level. Using DEXseq, we investigated differential exon usage across cell types and used stringent cutoffs to give confidence to the results, identifying how splice variants are changing across cell types over development as well as capturing novel splice variants. Nevertheless, across the inspected candidates that come up from the various comparisons as probable instances of differential exon usage, we notice instances where the current gene annotation is likely incorrect or incomplete (figs. S15, S16 and S18). We also observe different start sites for transcribing the same gene between as well as within the same cell type. This is observed, for example, in md1, where the female gametocytes appear to show several transcripts arising from each of the exons.

We also observe what appears to be spurious transcription in several genes, for example, in exon 1 of md1 and OAT (PF3D7_0608800) (Fig. 3E, fig. S17). This is likely due to high AT% of *P. falciparum* causing priming of poly-A tracts on first-strand cDNA (fig. S17). This has been reported in 10X libraries (*96*, *97*), and the AT richness of the *P. falciparum* genome makes it more pronounced, especially when looking at short reads (fig. S17). Interestingly, md1 displays anti-sense expression of a lncRNA in this locus as recently reported (*21*) (Fig. 3E). We observe similar examples where antisense expression appears in loci where exon usage is diminished (fig. S15, fig. S16A).

Despite the shortcomings of 10X based full-length RNAsequencing described above, the data helped give confidence to multiple instances of differential isoform or exon usage across cell types, as well as novel splicing events. While it's tempting to infer biological relevance to these splice variants, it should be noted that alternative splicing can be a consequence of stochastic noise in the splicing machinery, and can have no functional significance (*98*).

Our current undertaking of single-cell long read sequencing using the PacBio Sequel II is limited in throughput and cost. A new method relying on concatenation of cDNA molecules into longer fragments called MAS-Seq (Multiplexed Arrays Sequencing) (*99*) results in a 16-fold throughput increase compared to regular single-cell Iso-Seq® libraries. In addition, the new long read sequencing system from PacBio, Revio, provides three-fold increase in HiFi read throughput with a 25M SMRT Cell. The higher throughput with these advances may help overcome some of the problems above, however some are likely driven by the 10X 3' kit itself. Despite the sparsity, long read data accurately managed to cluster cell types and capture the topology of the development as with short read data (Fig. 3). In the near future, it is likely that long read sequencing can generate single-cell transcriptomes comparable to short-read based methods, and effectively replace them, with the additional advantage of exploring isoform variation.

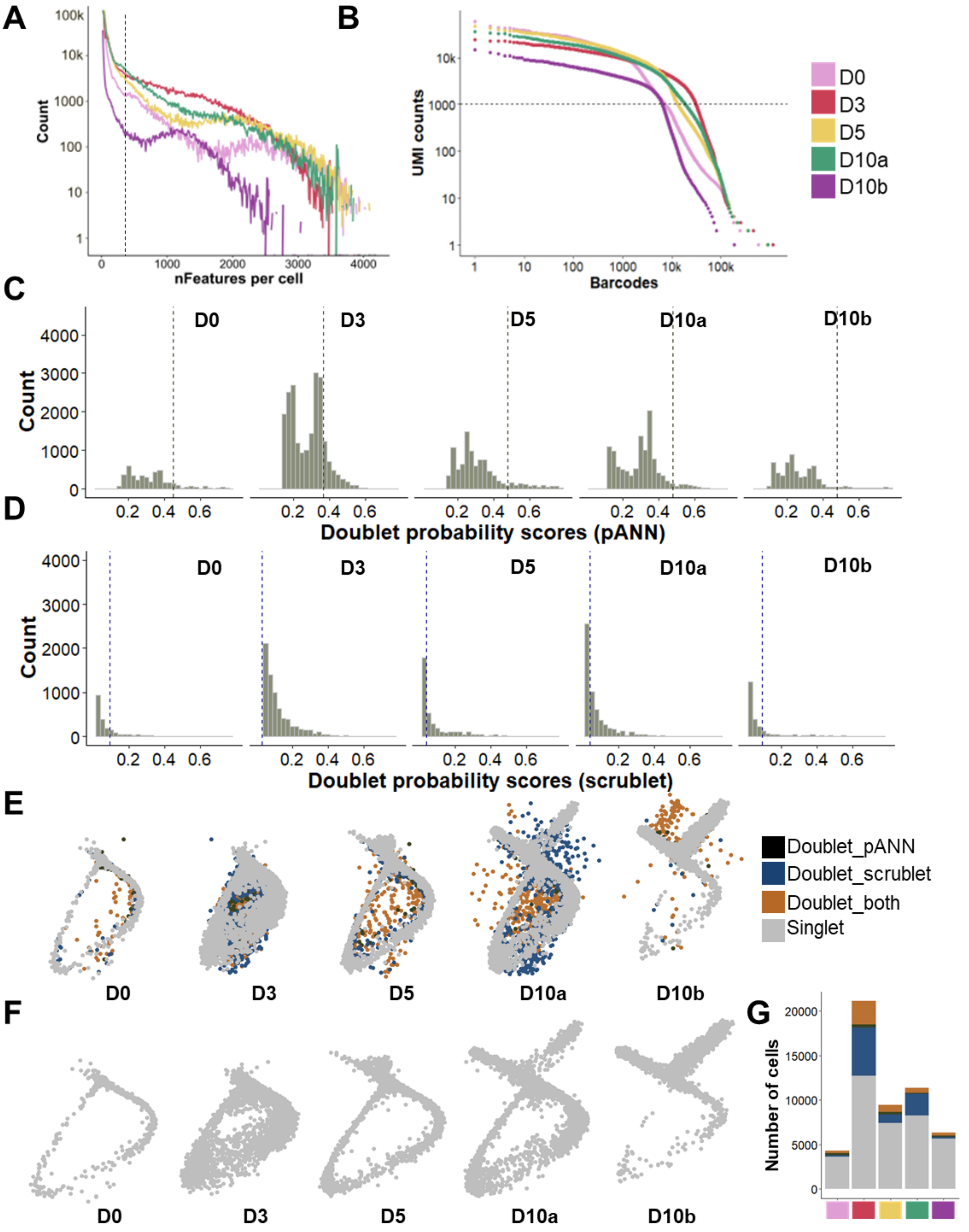


**Fig. S1. Quality control and integration of the datasets.** ***(A)*** *Genes per cell ("nFeatures") before filtering out poor quality cells, grouped by sampling day (D=day). The dotted line shows the*

*threshold of 300 genes required to pass QC.* ***(B)*** *UMI counts per cell before filtering out poor-quality cells, colored by sample. The dotted line shows the threshold of 1000 UMIs required to pass QC.* ***(C)*** *Doublet probability scores of doublet detection algorithms DoubletFinder and (D) scrublet showing cutoff values per sample with dotted lines. Cutoffs of 0.45, 0.37, 0.48, 0.48, 0.48 were used for DoubletFinder, and cutoffs of 0.09, 0.03, 0.04, 0.04, 0.09 were used for scrublet for samples D0, D3, D5, D10a and D10b, respectively.* ***(E)*** *PCA embedding for each sample before filtering for poor-quality cells, highlighting the cells that were excluded as putative doublets by the different detection algorithms.* ***(F)*** *Each sample's PCA plot after filtering out doublets and poor-quality cells.* ***(G)*** *Number of cells per sample detected as doublets*

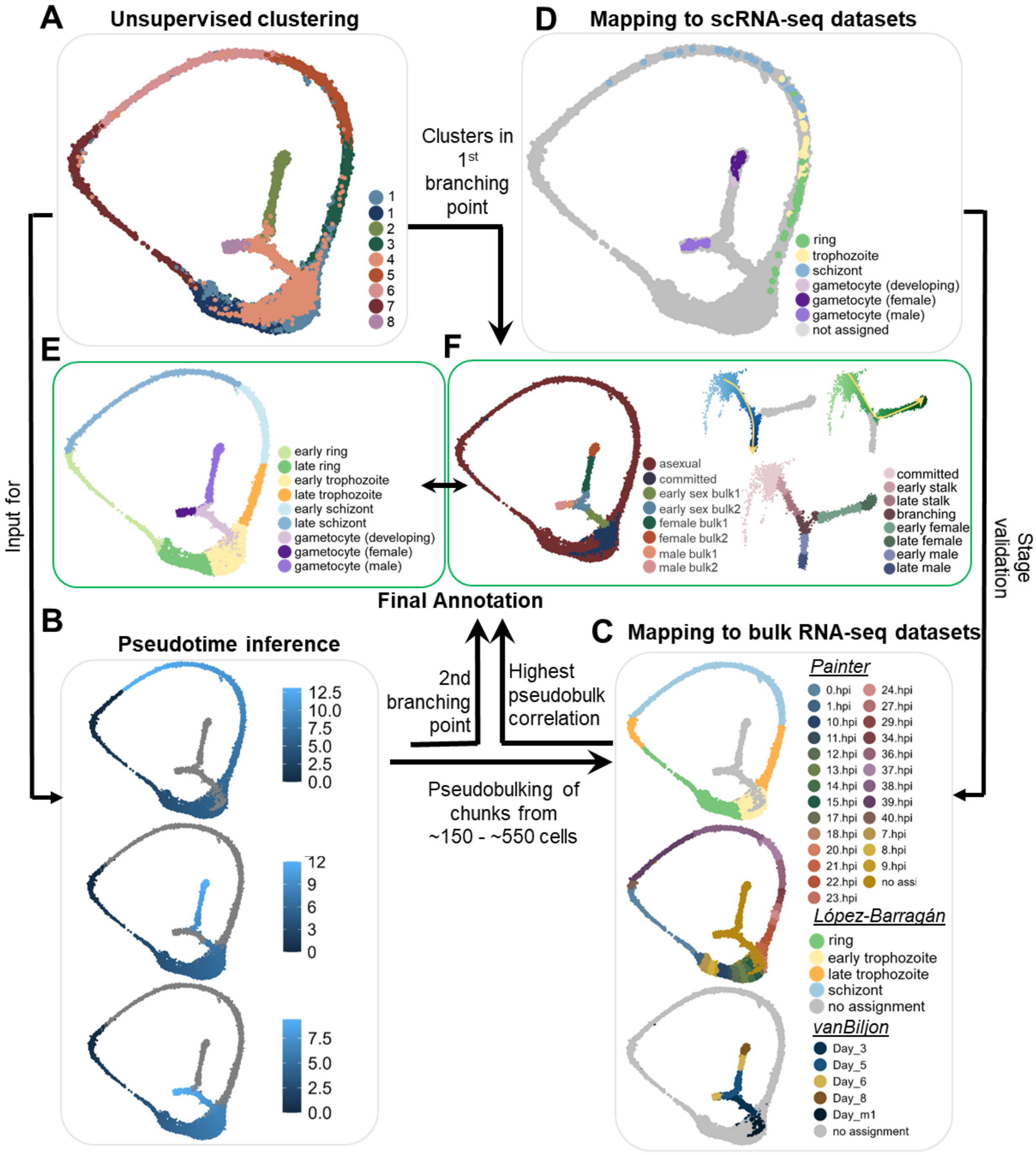


**Fig. S2. Assignment of developmental stages to QCed cells by clustering and mapping to previously published datasets.** *(A) UMAP representation of Louvain clustering with the resolution of 0.2. Cluster 4 in the first branching point (asexual/sexual) includes committed gametocytes and the stalk. (B) Pseudotime inference was performed using slingshot with cluster 7 as starting point and with clusters 2, 6, and 8 as endpoints in order to assign pseudotimes to all asexual cells and*

*both males and females. The pseudotime assignments were the basis for assembling pseudobulks of 150 or 540 cells (540 cells split the asexual cycle in 48 chunks). (C) Pseudobulks were made from the single cell data and then compared to bulk RNAseq datasets including (59) (top), (60) (middle), and (61) (bottom). The strongest correlation of the pseudobulks to these datasets was used to assign asexual stages. (D) Mapping of published P. falciparum SmartSeq2 dataset (62) performed with scmap. The mapping was used to validate the correlation results from the pseudobulking. (E) Final annotations in the assigned stages comprising asexual (early and late rings, trophozoites, and schizonts) and sexual (gametocyte (developing), gametocyte (female), gametocyte (male) stages. (F) Clustering of the sexual stages leading from the committed to the late sexual stages. To explore the transcriptional changes underlying sexual stage commitment and development, the subset of 8958 cells representing the non-asexual stages were analyzed. Slingshot was used to infer cell lineages and pseudotime, fitting a curve to each of these trajectories using the underlying UMAP embedding and annotated cell clusters. Cells were ordered according to their relative position along each developmental trajectory by their pseudotime, leading from the sexually committed cells ('committed') to the male and female bulk 2 clusters. Developing, male, and female gametocytes were re-grouped into different sub-stages (i.e. early and late) by splitting them along pseudotime using values to obtain an equal proportion of early and late stages. The group of cells that precede clear bifurcation of male and female gametocytes from the developing gametocytes are labeled as 'branching'. Arrows represent the progression of pseudotime along the two lineages.*

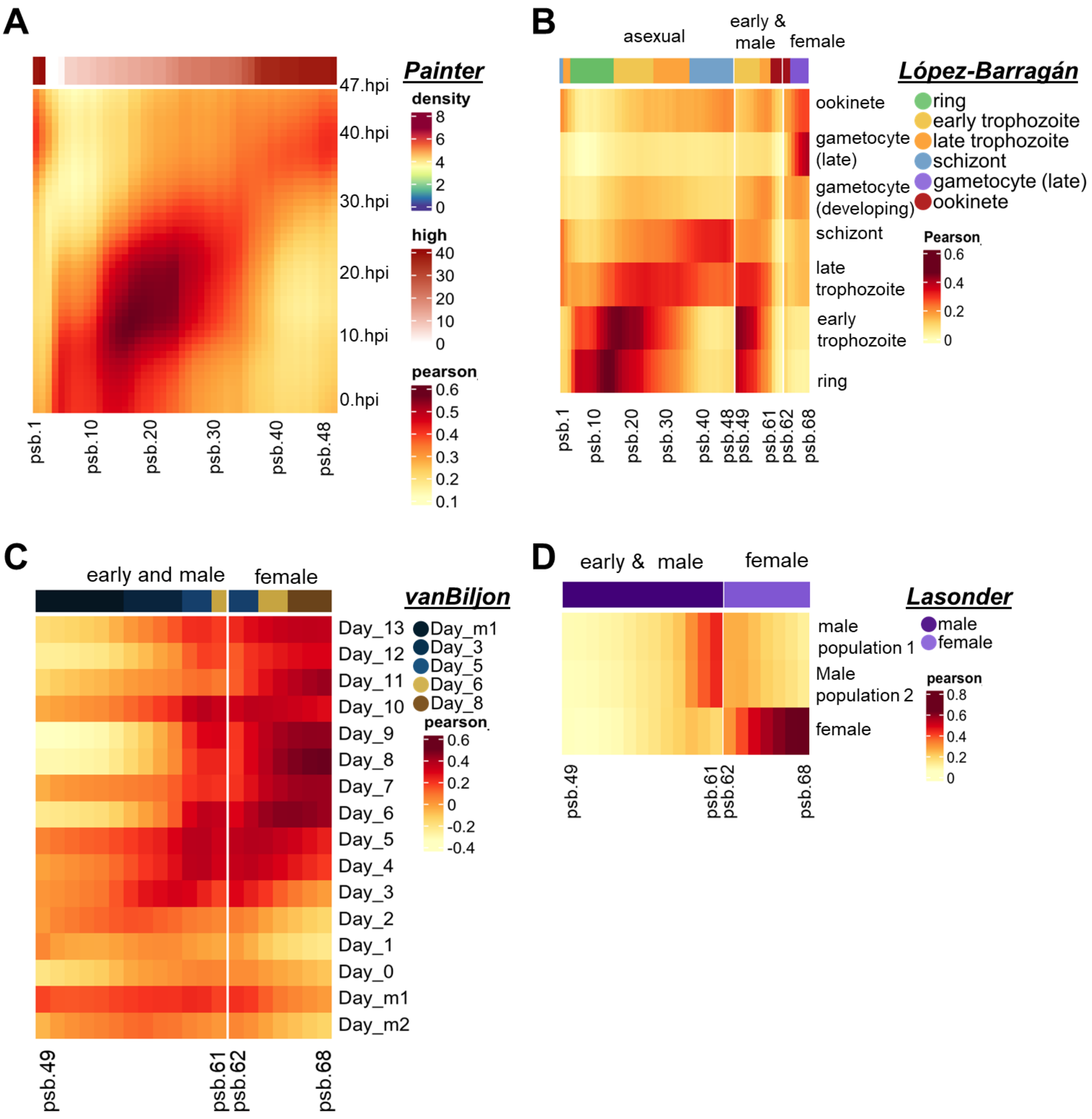


**Fig. S3. Correlation with published bulk RNA-seq datasets.** *Heatmaps are colored by correlation of each pseudobulk (n=48 for asexual stages, each with 540 cells; n=20 for stalk and sexual stages, each with 150 cells) to the respective lifecycle stages of the following four datasets.* ***(A)*** *48 pseudobulks (psb1-48) corresponding to the asexual stages are correlated to a bulk RNAseq 48-hour asexual time course dataset* (59)*, showing a strong correlation.* ***(B)*** *Correlation was performed with bulk RNA-seq data exploring asexual and sexual intraerythrocytic development*

*from (60), again showing a clear correlation of pseudobulk data with the asexual bulk samples.* ***(C)*** *In a bulk RNAseq study examining gametocytogenesis again without separating the sexes (61), our pseudobulks representing early sexual development (starting at psb.49) correlate most strongly with bulk samples prior to gametocyte induction. The later sexual pseudobulks follow the temporal progression of samples while not distinguishing male and female gametocytes.* ***(D)*** *Male and female pseudobulks were compared to the bulk RNA-seq dataset created by (22), where male and female gametocytes were sorted by FACS, showing that the single cell data correlates well with the bulk sex-specific data.*

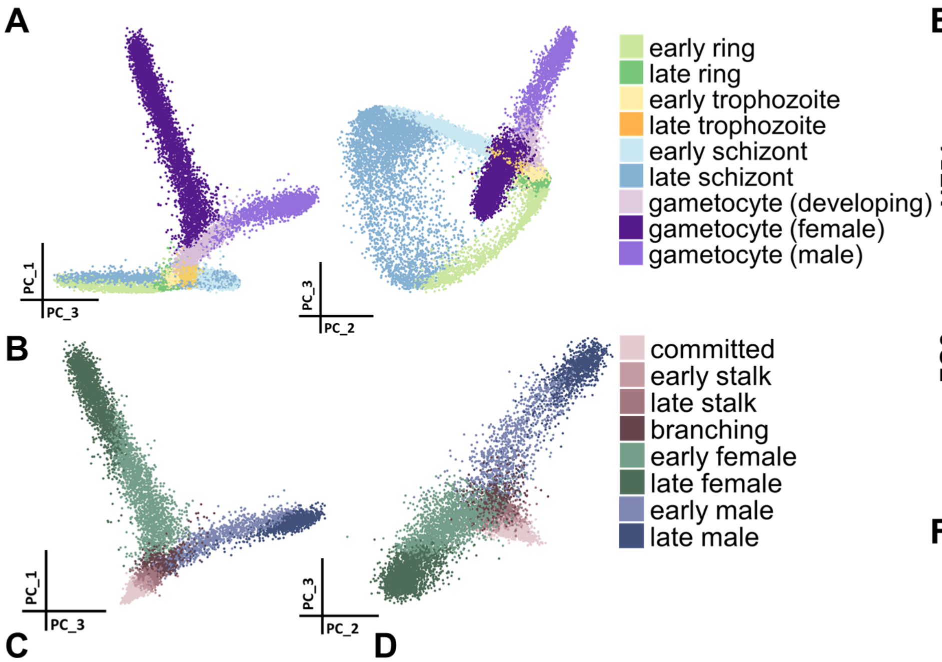


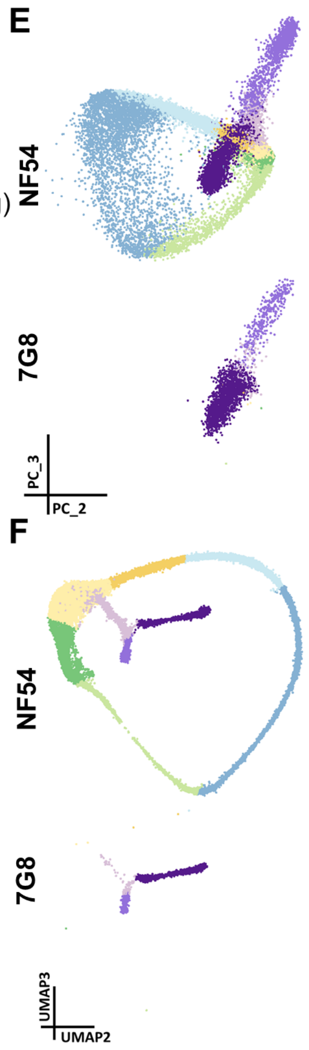


**C**

| Stage | Cells |
|---|---|
| early ring | 1633 |
| late ring | 5438 |
| early trophozoite | 9635 |
| late trophozoite | 3801 |
| early schizont | 3805 |
| late schizont | 4355 |
| gametocyte (developing) | 3180 |
| gametocyte (female) | 4003 |
| gametocyte (male) | 1775 |

**D**

| Stage | female lineage | male lineage | Pseudotime (female/male) |
|---|---|---|---|
| committed | 1315 | 1315 | <=0.74 |
| early stalk | 615 | 615 | >0.74 & <=1.165 |
| late stalk | 610 | 610 | >1.165 & <=1.62 |
| branching | 551 | 551 | >1.62 & <=2/1.98 |
| early female | 1958 | 0 | >2 & <= 3.091 |
| late female | 1945 | 0 | > 3.091 |
| early male | 0 | 985 | >1.98 & <=2.3105 |
| late male | 0 | 979 | >2.3105 |

**Fig. S4. PCA Visualisation and distribution of cells within each assigned stage and strain.** ***(A)*** *PCA scatter plot of the 37624 single cells collected at four sampling timepoints, coloured by assigned life cycle stage based on clustering and reference mapping, visualized for PC1/PC3 and PC2/PC3. The topology observed is similar to that of the UMAP dimensionality reduction in Figure 1A.* ***(B)*** *PCA scatter plot of the 8958 single cells representing the non-asexual stages coloured by the assigned re-grouped life cycle stages, visualized for PC1/PC3 and PC2/PC3. The topology observed is similar to that of the UMAP dimensionality reduction in Fig 1B.* ***(C)*** *Distribution of cells across the assigned stages assigned as in (A).* ***(D)*** *Distribution of cells across the assigned sub-stages and the pseudotime values associated with them, along the female and male lineages.* ***(E)*** *PCA plot of the 37624 single cells displaying the distribution of the 2 strains in the dataset, coloured by assigned life cycle stage based on clustering and reference mapping, visualized for PC2/PC3. (F) UMAP plot of the 37624 single cells displaying the distribution of the 2 strains in the* *dataset, coloured by assigned life cycle stage based on clustering and reference mapping, visualized for UMAP2/UMAP3.*

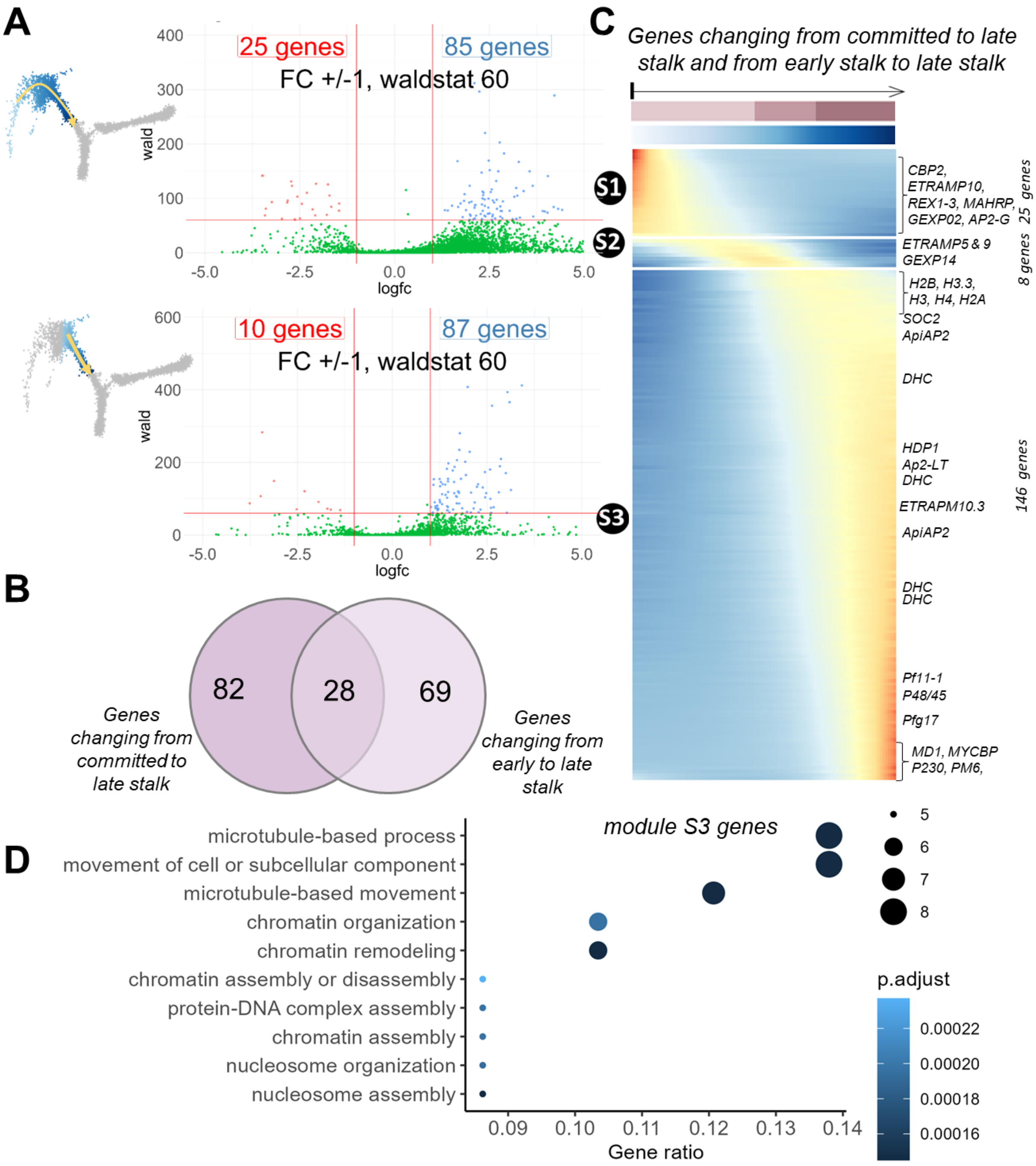


**Fig. S5. Patterns of gene expression from commitment to late stalk gametocytes.** ***(A)*** *Volcano plots showing the 110 and 97 genes, and* ***(B)*** *Venn diagram showing 179 genes in total, that change expression significantly (fold change > 1) from committed to late stalk, and early stalk to late stalk, respectively. tradeSeq's startVsEnd test was used to assess the differential expression between the start and end point of the lineages. UMAP panels on the left show the cells considered for comparison, gradient-colored according to pseudotime values. Results of these*

*analyses are presented in data S2.* ***(C)*** *The heatmap shows the smoothed expression profiles of the 179 genes, assigned to three distinct modules as follows: S1, comprising genes that decrease following commitment; S2, comprising genes that appear to be transiently expressed after commitment; S3, comprising genes that increase in expression along the stalk following commitment. Select genes mentioned in the main text are highlighted on the heatmap. Cells are ordered according to their pseudotime and colored row bars indicate life stage and pseudotime. Rank-two ellipse seriation (100) was used to rank genes based on a correlation matrix of the fitted values for each module's genes.* ***(D)*** *Module S3 comprising genes that start to transcribe within the stalk are enriched in factors necessary for motility and microtubule associated processes, and those involved chromatin remodeling and organization, as investigated by GO enrichment analysis. The top 10 significant results from the output were visualized with the "dotplot" function in the clusterProfiler package. The x-axis and y-axis represent the gene ratio, and GO terms, respectively. The color scale represents the adjusted p-value. Complete GO enrichment results are provided in data S2. No significant results (p-value 0.05) were obtained upon GO enrichment of modules S1 and S2.*

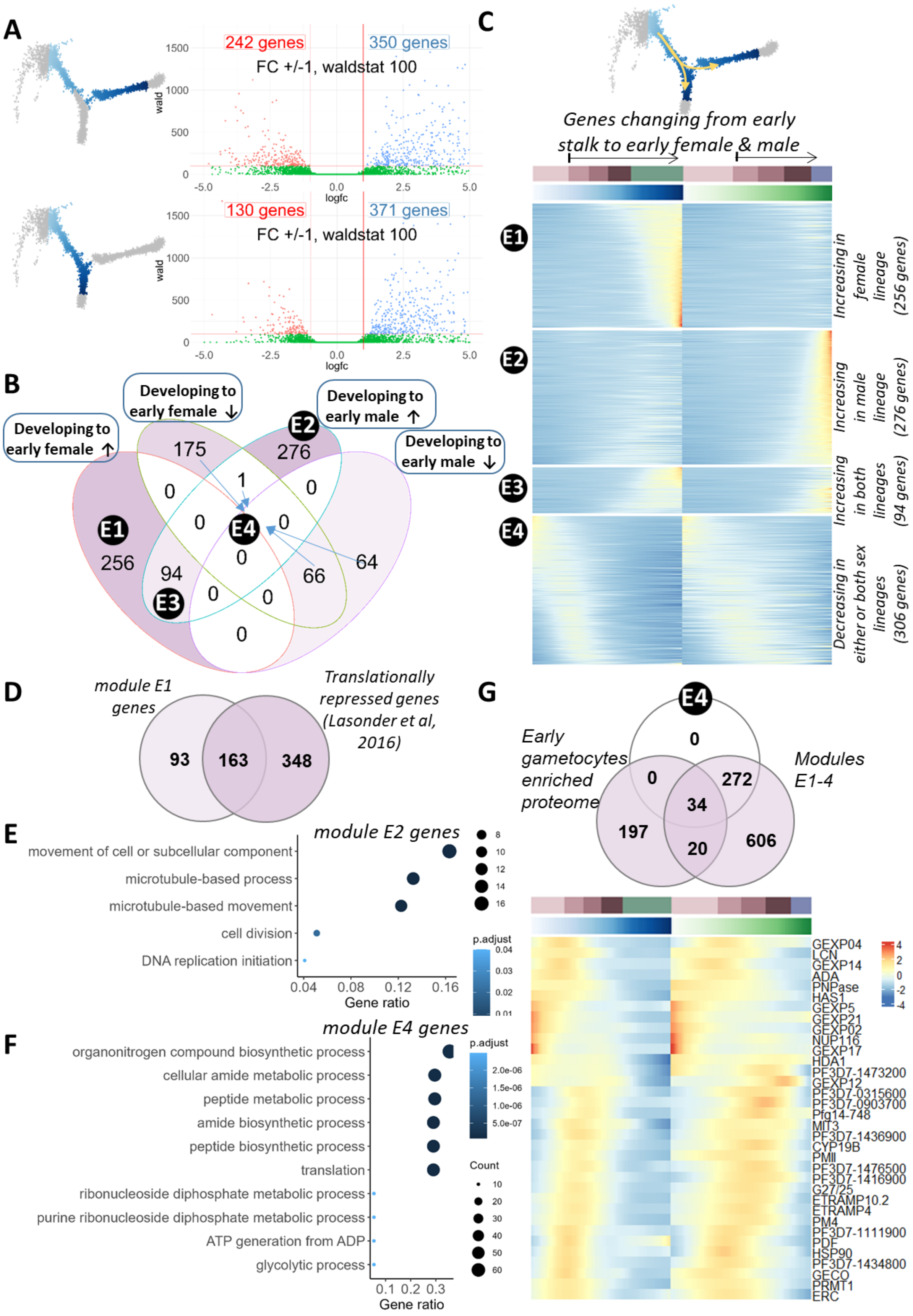

A
242 genes
350 genes
FC +/-1, waldstat 100
130 genes
371 genes
FC +/-1, waldstat 100
wald
logfc
B
Developing to early female ↑
Developing to early female ↓
Developing to early male ↑
Developing to early male ↓
E1
E2
E3
E4
256
175
276
94
66
64
1
C
Genes changing from early stalk to early female & male
Increasing in female lineage (256 genes)
Increasing in male lineage (276 genes)
Increasing in both lineages (94 genes)
Decreasing in either or both sex lineages (306 genes)
D
module E1 genes
Translationally repressed genes (Lasonder et al, 2016)
93
163
348
E
module E2 genes
movement of cell or subcellular component
microtubule-based process
microtubule-based movement
cell division
DNA replication initiation
Gene ratio
p.adjust
F
module E4 genes
organonitrogen compound biosynthetic process
cellular amide metabolic process
peptide metabolic process
amide biosynthetic process
peptide biosynthetic process
translation
ribonucleoside diphosphate metabolic process
purine ribonucleoside diphosphate metabolic process
ATP generation from ADP
glycolytic process
Gene ratio
p.adjust
Count
G
Early gametocytes enriched proteome
Modules E1-4
0
272
34
197
20
606
GEXP04
LCN
GEXP14
ADA
PNPase
HAS1
GEXP5
GEXP21
GEXP02
NUP116
GEXP17
HDA1
PF3D7-1473200
GEXP12
PF3D7-0315600
PF3D7-0903700
Pfg14-748
MIT3
PF3D7-1436900
CYP19B
PMII
PF3D7-1476500
PF3D7-1416900
G27/25
ETRAMP10.2
ETRAMP4
PM4
PF3D7-1111900
PDF
HSP90
PF3D7-1434800
GECO
PRMT1
ERC

**Fig. S6. Patterns of gene expression from early stalk gametocytes to early sex gametocytes.** ***(A)*** *Volcano plots showing the 592 and 501 genes (932 in total) that change expression significantly (fold change > 1) from early stalk gametocytes to early male and early female gametocytes. tradeSeq's startVsEnd test was used to assess the differential expression between the start and end point of the lineages. UMAP panels on the left show the cells considered for comparison gradient-colored according to pseudotime values. Results of these analyses are presented in data S2.* ***(B)*** *These 932 genes were assigned to four distinct modules as follows: E1, comprising genes that increase in the female lineage; E2, comprising genes that increase in male lineage; E3, comprising genes that increase in both lineages; E4, comprising genes that decrease in both lineages* ***(C)*** *The heatmap shows smoothed expression profiles of the 932 genes in total that show differential expression between the early stalk and the early male or female stages (cells colored in the UMAP, top panel). Cells are ordered according to their pseudotime and colored row bars indicate life stage and pseudotime. Rank-two ellipse seriation (100) was used to rank genes based on a correlation matrix of the fitted values for each module's genes. Results of these analyses are presented in supplementary file data S2.* ***(D)*** *Module E1 genes that increase in the female lineage have a significant overlap (p-value < 1.1e-66, hypergeometric test) with those found in a translationally repressed state in female gametocytes as described by (22).* ***(E)*** *Module E2 comprising genes increasing in male lineage is enriched in factors necessary for motility and microtubule associated processes, cell division and cell cycle progression as investigated by GO enrichment analysis. The 5 significant results (p-value < 0.05) from the output were visualized with the "dotplot" function in the clusterProfiler package.* ***(F)*** *Module E4 comprising genes that are decreasing in either or both sex lineages following sexual commitment contain metabolic processes as the top GO terms. The top 10 results were visualized with the "dotplot" function in the clusterProfiler package. The x-axis and y-axis represent the gene ratio, and GO terms, respectively. The color scale represents the adjusted p-value. GO enrichment analysis for module E3 resulted in a single GO term 'microtubule-based movement', while no significant results (p-value < 0.05) were obtained for module E1. GO enrichment results for genes in module E2, E3 and E4 are provided in supplementary file data S2.* ***(G)*** *Module E4 genes overlap with proteins that are putatively involved in erythrocyte remodeling enriched in the proteome of early gametocytes ( (16) (p-value <1.09e-08, hypergeometric test). The E4 module also contains several genes that are identified as markers of early gametocytes including Pfg14-748 and G27/25.*

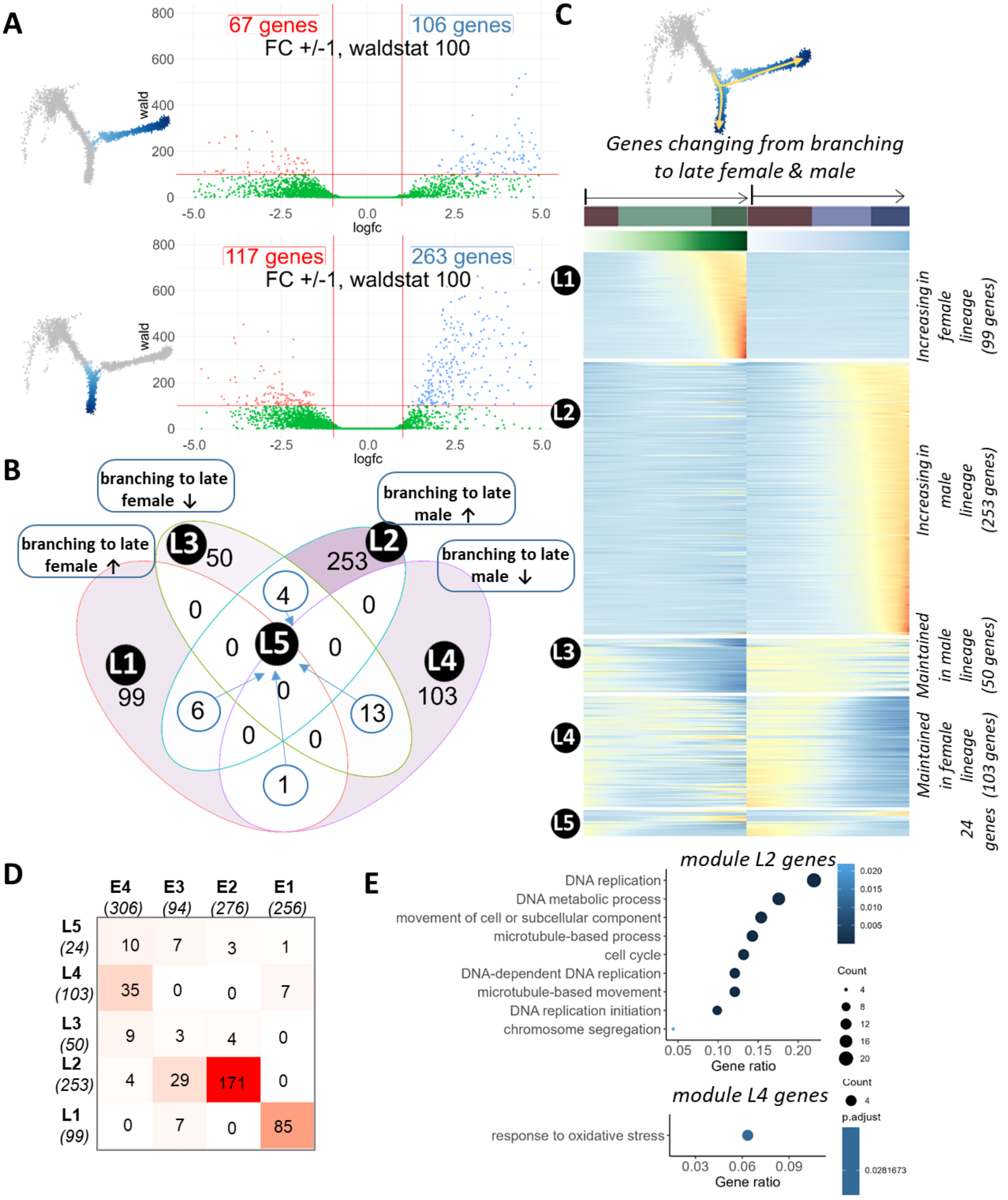


| | E4 (306) | E3 (94) | E2 (276) | E1 (256) |
|---|---|---|---|---|
| L5 (24) | 10 | 7 | 3 | 1 |
| L4 (103) | 35 | 0 | 0 | 7 |
| L3 (50) | 9 | 3 | 4 | 0 |
| L2 (253) | 4 | 29 | 171 | 0 |
| L1 (99) | 0 | 7 | 0 | 85 |

**Fig. S7. Patterns of gene expression from branching to late sex gametocytes.** *(A) Volcano plots showing the 173 and 380 genes that change expression significantly (fold change > 1) between branching and late female and male stages, respectively. tradeSeq's startVsEnd test was used to assess the differential expression between the start and end point of the lineages. UMAP panels on* *the left show the cells considered for comparison, gradient-colored according to their pseudotime values. These results are tabulated in data S2* ***(B)*** *Combined results from A (529 genes in total) were categorized into five modules based on their mode of expression in the male and female lineages as follows: L1, comprising genes that increase in female lineage; L2, comprising genes that increase in male lineage; L3, comprising genes that are maintained only in male lineage following 'branching' and decreasing in female lineage; L4, comprising genes that are maintained only in female lineage following 'branching' and decreasing in male lineage; L5, comprising the rest of the genes that includes: 13 genes that decrease in both lineages, 6 genes that increase in both lineages, 4 genes that increase in male lineage, but decrease in female lineage, and 1 gene that increases in female lineage and decreases in male lineage, following 'branching'.* ***(C)*** *The heatmap shows smoothed expression profiles of the 529 genes that show differential expression between the branching and the late male or female stages (cells colored in the UMAP, top panel), with expression modules L1-L5 as described above labeled. Cells are ordered according to their pseudotime and colored row bars indicate life stage and pseudotime. Rank-two ellipse seriation (100) was used to rank genes based on a correlation matrix of the fitted values for each module's genes. Results of these analyses are presented in supplementary file data S2.* ***(D)*** *Genes in module L1 and L2 show overlap with those in modules E1 and E2, respectively.* ***(E)*** *Module L2 comprising genes increasing only in the male lineage are enriched in factors necessary for motility and microtubule associated processes, and DNA replication processes, as investigated by GO enrichment analysis. These factors are necessary for the motility of male gametocytes and the ookinete and the rapid 3-fold genome replication that male gametocytes undergo following ingestion by a mosquito. The 9 significant results from the output were visualized with the "dotplot" function in the clusterProfiler package. GO enrichment analysis for module L4 resulted in a single GO term 'response to oxidative stress'. The x-axis and y-axis represent the gene ratio, and GO terms, respectively. The color scale represents the adjusted p value. No significant results (p-value < 0.05) were obtained for modules L1, L3 or L5. GO enrichment results for genes in module L2 and L4 are provided in supplementary file data S2.*

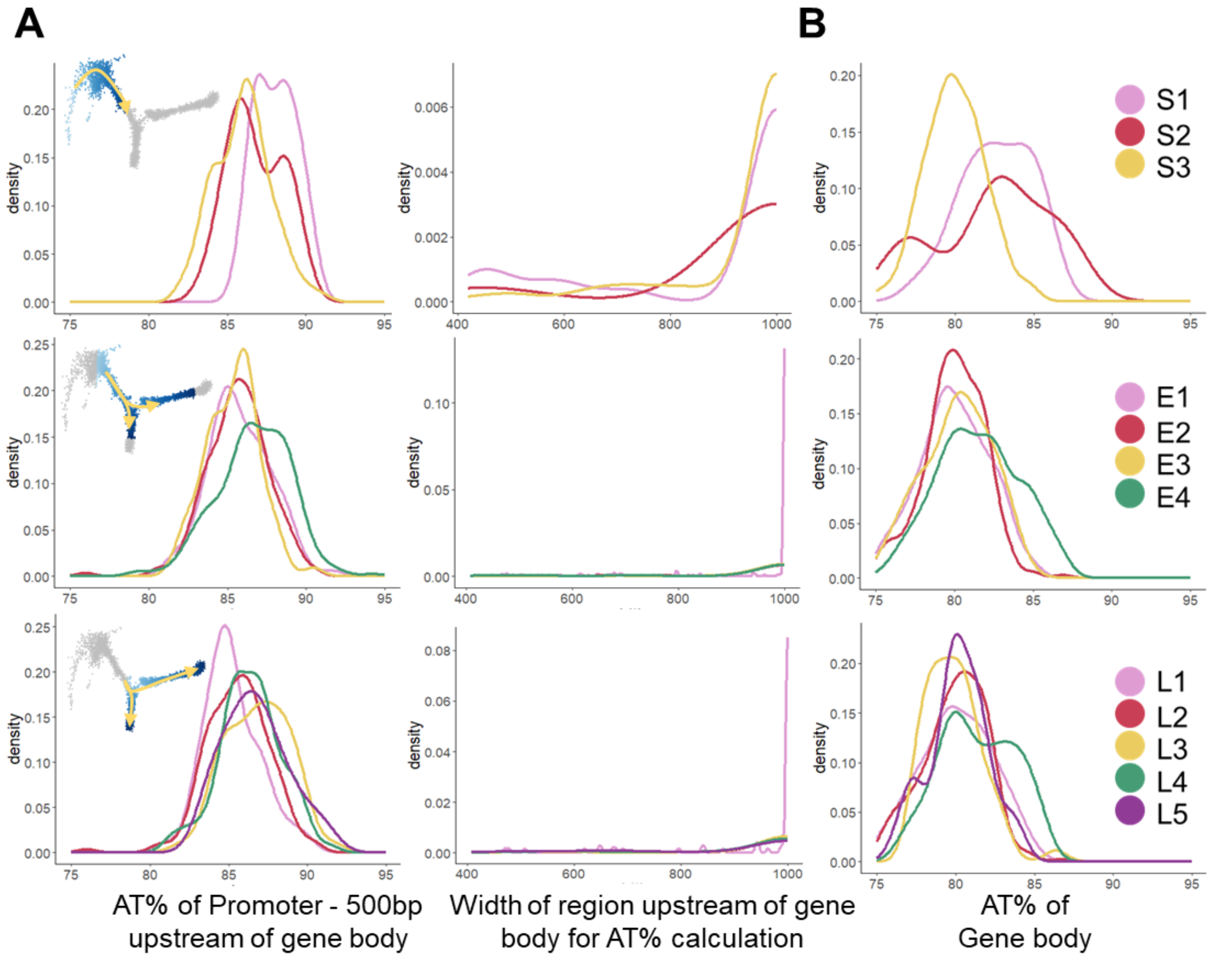


**Fig. S8. A slightly higher AT content of promoter region differentiates gene expression modules E4, L3, L4 and L5 that are transiently expressed along gametocyte development.** ***(A)*** *A recent study uncovered the role of an AT-rich interaction domain containing protein, PfARID, in regulating sexual development* (23)*. AT% of promoter regions across the modules was plotted by extracting 500 bp upstream of genes, excluding regions if they overlapped other genes (Methods). Modules with transient expression early in gametocyte development, such as E4 and L3 show a clear shift in the AT% peak, as do L2, L4 and L5 to a lesser extent. The width of the upstream regions (considering up to the presence of another gene closeby) used for the genes is presented in the right panel.* ***(B)*** *AT% of the gene body of genes is lower than that of the promoter regions and across the modules E1-E4 and L1-L5 display no clear differences except for module E4 and L4, which have a wider range. Modules S1 and S2 (containing 25 and 8 genes respectively) show a higher AT content in their gene bodies, with no differences in the AT% of the promoter regions.*

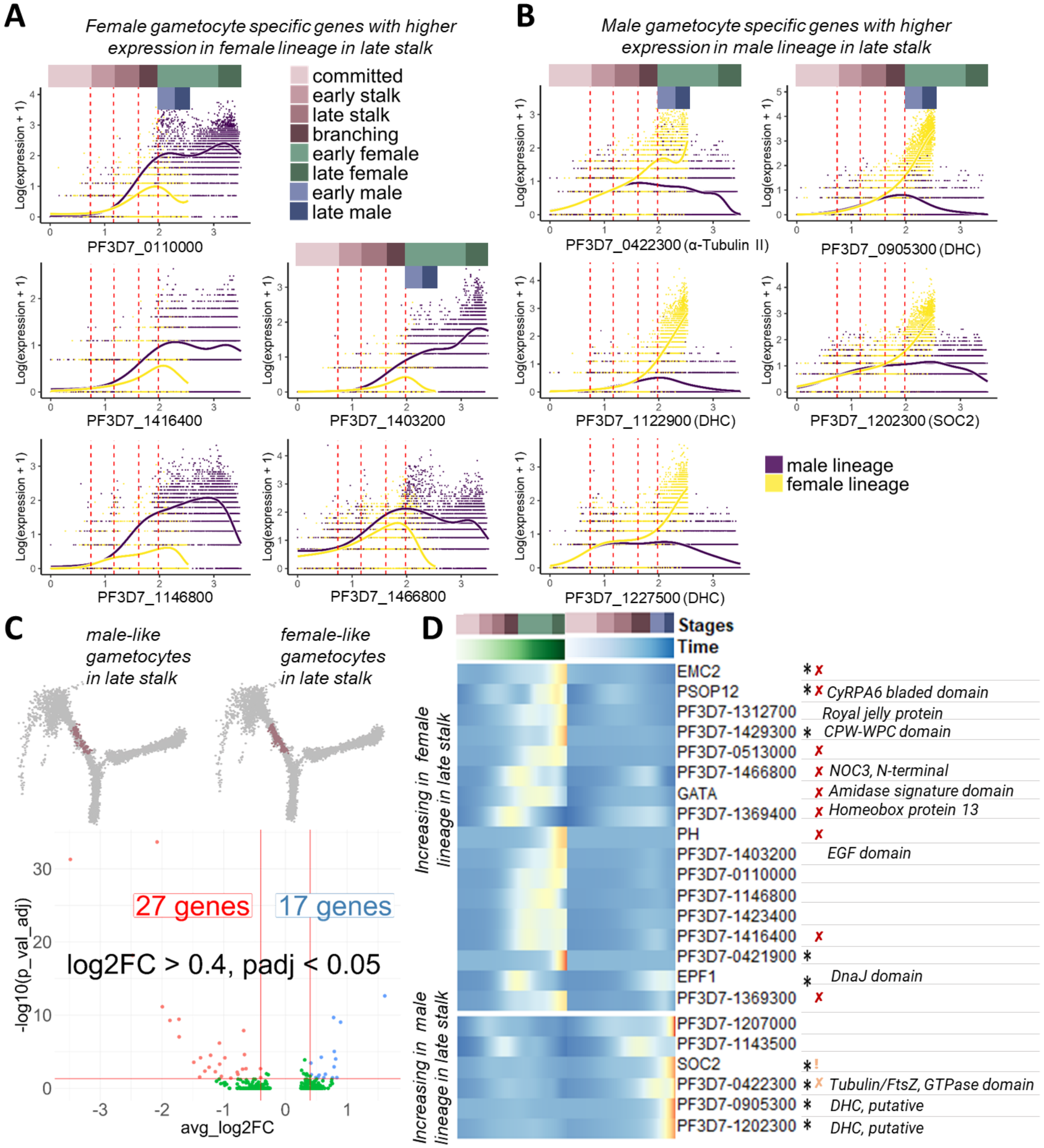


**Fig. S9. Differential expression analysis between male-like and female-like gametocytes within the late stalk gametocytes.** ***(A)*** *Five genes from module S3 as described in Fig. S5 (PF3D7_0110000, PF3D7_1146800, PF3D7_1416400, PF3D7_1403200, PF3D7_1466800) that show increased expression in the female lineage compared to male lineage within the late stalk.* ***(B)*** *Five other genes (PF3D7-0422300, PF3D7-0905300, PF3D7-1122900, PF3D7–1202300, PF3D7-1227500) that show increased expression in the male lineage compared to the female*

*lineage within the late stalk. A and B show tradeSeq smoother plots that represent expression of a gene over pseudotime along male (yellow) and female (violet) lineages by plotting log-transformed counts fitted by tradeSeq using a generalized additive model (GAM).* ***(C)*** *Genes in A and B start to show transcription within the stalk and continue to become stronger along sexual development, and these genes were used to define putative early male and female cells as follows: Cells within the late stalk (610 cells) were assigned as 'female-like' (133 cells, right UMAP) by summing the expression of the five genes in A across the 610 cells and selecting those with expression above the mean of these summed values. Similarly, 'male-like' (203 cells, left UMAP) were assigned by ranking the cells as above with the five genes from B and selecting unique cells, leaving 274 cells unclassified. Differential expression between the male-like and female-like groups of cells was performed using the FindMarkers function from seurat, using MAST DE test with a logFC of 0.4, the complete output of which is presented in data S3. 44 genes are differentially expressed along the male and female lineage within the late stalk.* ***(D)*** *Upon manual inspection of these 44 genes, 23 show a clear difference in expression between the lineages, as visualized by plotting the log-transformed counts fitted by tradeSeq using a GAM to represent expression over pseudotime using tradeSeq's smoother plots (data S4). The heatmap shows smoothed expression profiles of these 23 genes that show clear male-lineage specific increase (top panel) and female-lineage specific increase (bottom panel). Cells are ordered according to their pseudotime and colored row bars indicate life stage and pseudotime. Rank-two ellipse seriation (100) was used to rank genes based on a correlation matrix of the fitted values for each module's genes. Phenotypic information from Phenoplasm (28) is provided next to the gene ID/name. X indicates that the gene is refractory to deletion, while ! indicates an attenuated phenotype in the sexual or asexual life cycle stages. Lighter colored icons indicate data from orthologs. * on the gene row indicates that the gene is a likely target of AP2-G binding in stage I gametocytes (15).*

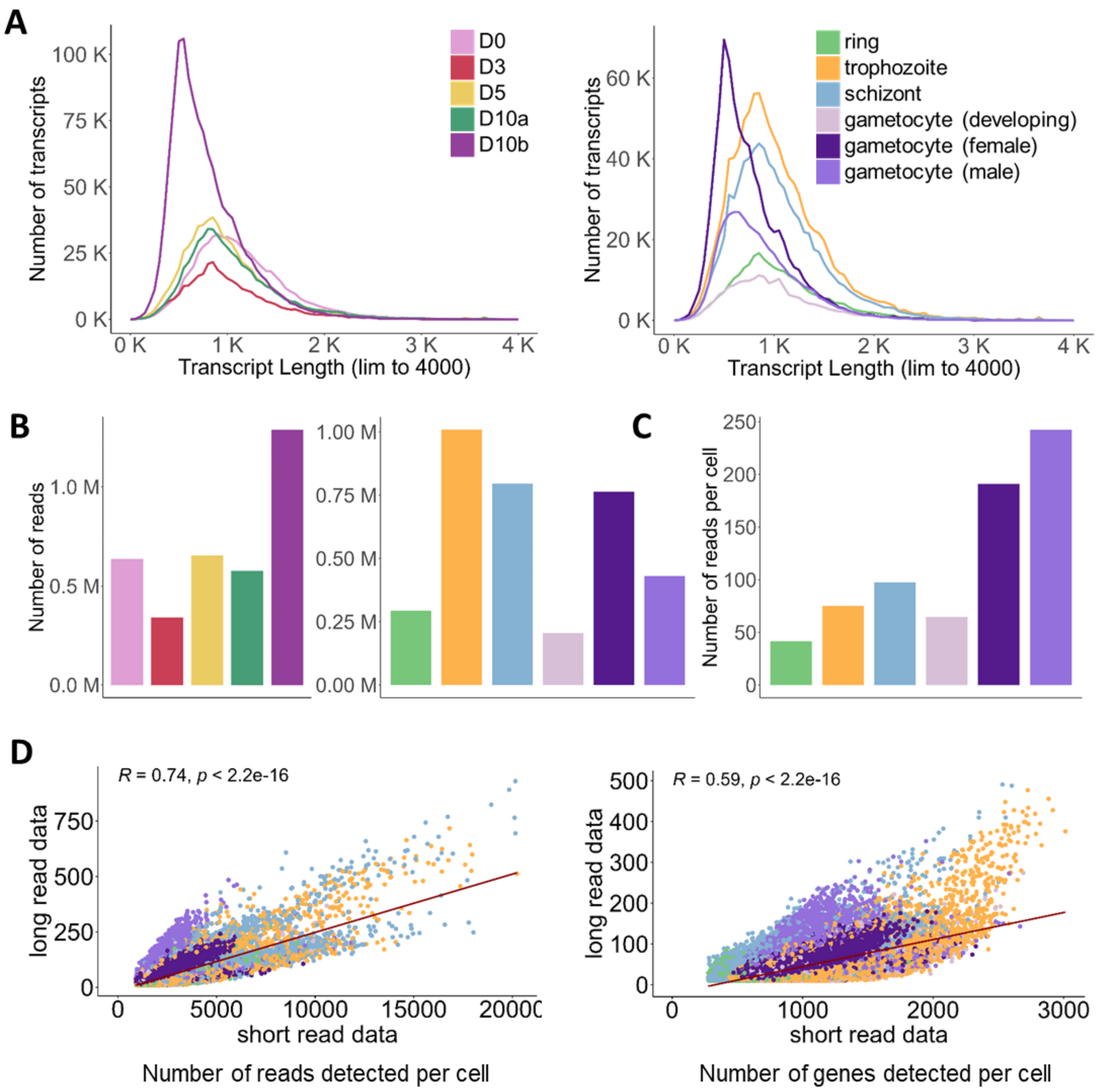


**Fig. S10. Long read sequencing output across the stages.** *For each sampling day, IsoSeq reads were QCed to remove primer overhangs, polyA tails, and artificial concatemers. UMI and Cell Barcode information on the long reads was used to cluster reads by the unique founder molecules.* ***(A)*** *Transcript length distribution based on long read data, grouped by sampling day (left) and assigned stage (right).* ***(B)*** *Number of long reads detected, grouped by sampling day (left) and assigned stage (right).* ***(C)*** *Mean number of long reads detected per cell, grouped by assigned stage.* ***(D)*** *Correlation of number of short reads and long reads detected per cell (left) and per gene (right).*

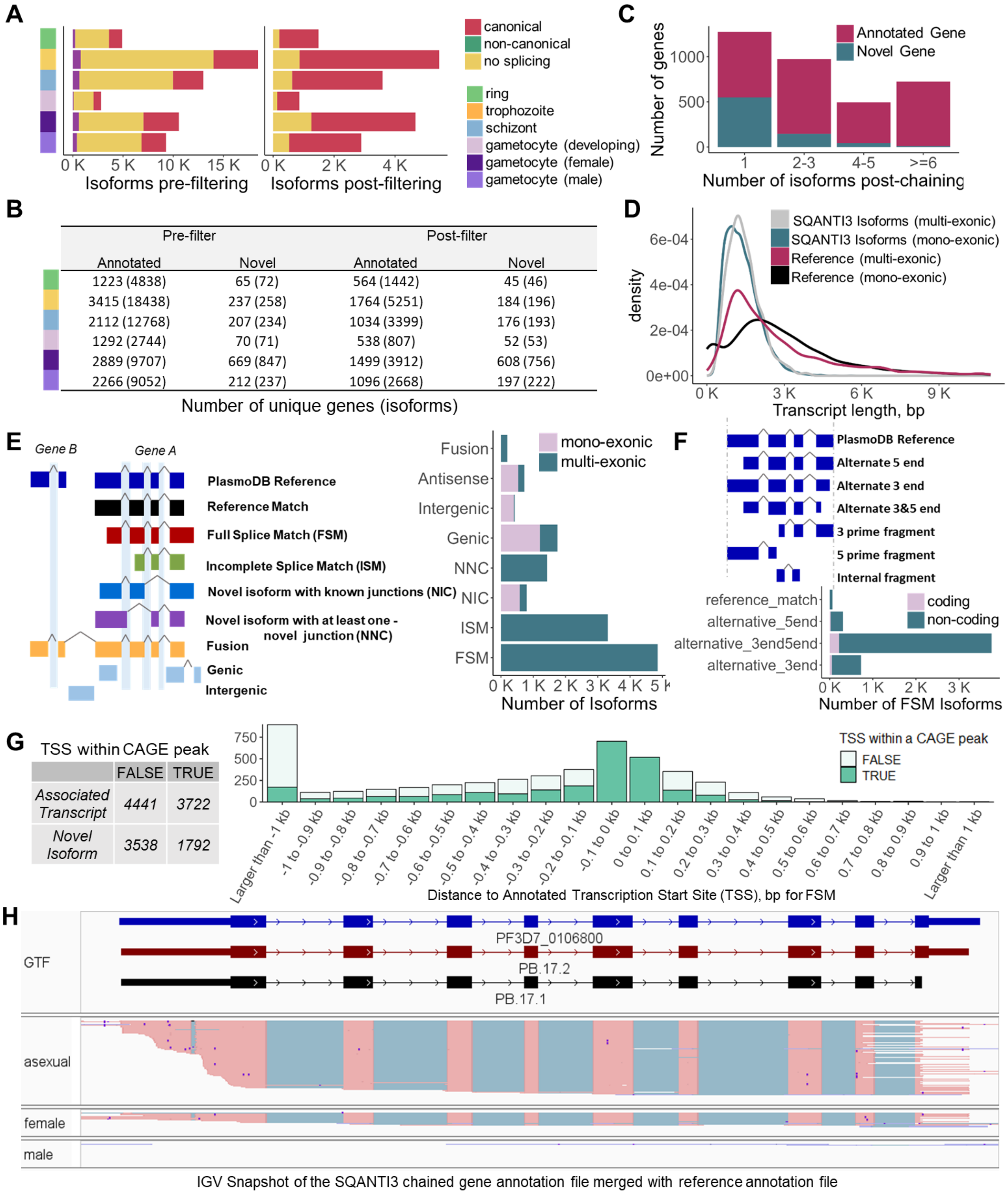


**Fig. S11. Stage labels from short read data allow generation of stage-specific isoforms.** ***(A)*** *Pre- and post-filtering summary of number of isoforms detected and breakdown of the splicing events detected for these isoforms for each life stage. Filtering was done to remove isoforms with > 90% adenines downstream of the transcription end site, those with non-canonical splicing junctions, and*

*transcripts with single exons (by using sqanti3_filter.py), while retaining transcripts containing full splice matches and novel isoforms.* ***(B)*** *Pre- and post-filtering summary of number of unique genes detected and the number of isoforms corresponding to these genes for each stage, classified according to whether they are identified as annotated or novel genes. Only the post-filtered isoforms are retained for further analysis by chaining to get non-redundant isoforms across all the stages.* ***(C)*** *Following chaining of the isoforms from all the cell types to obtain unique isoforms representing the full dataset, the plot shows the number unique genes binned according to the number of isoforms per gene, and categorized into whether they are novel genes or associated with annotated genes.* ***(D)*** *Transcript length distribution displayed as a kernel density plot for the mono-exonic and multi-exonic isoforms post-chaining, classified by SQANTI3 (mean length, mono-exonic = 1089 bp, multi-exonic = 1439 bp), compared to the 5791 reference transcripts currently annotated on PlasmoDB v59 (mono-exonic = 3144 bp, multi-exonic = 2920 bp).* ***(E)*** *SQANTI3 classification of sub-categories for the isoform events detected in the dataset as shown in the right panel (reproduced from https://isoseq.how/classification/categories.html, which has detailed description of these categories).* ***(F)*** *SQANTI3 classification of full splice matches into sub-categories, showing the number of coding and non-coding transcripts among them. Among full-splice matched isoforms (4828), the majority of events are alternate 5' and 3' ends.* ***(G)*** *Comparison of 5' ends of isoforms detected in this dataset, and the distance to annotated TSSs for full-splice match isoforms, highlighting their presence within the CAGE peaks as determined by long read sequencing of 5' capped capped mRNA from intra-erythrocytic asexual developmental stages (71).* ***(H)*** *Gene annotation files based on long reads from each cell type as well as the annotation chained by merging across the cell types is provided as a resource in data S7 and at Zenodo repository (DOI:10.5281/zenodo.8139823). This is an example region containing gene PF3D7_0106800, showing the detected isoforms PB.24.1 and PB.24.2 are classified as full splice matches, where PB.24.2 is sub-categorised as 'reference_match' and PB.24.1 as 'alternate_3end". The gene model from the reference annotation from PlasmoDB is shown in blue and SQANTI3 identified isoforms are colored maroon. Long reads from asexual (ring, trophozoite, schizont), late female and late male stages are shown in separate tracks.*

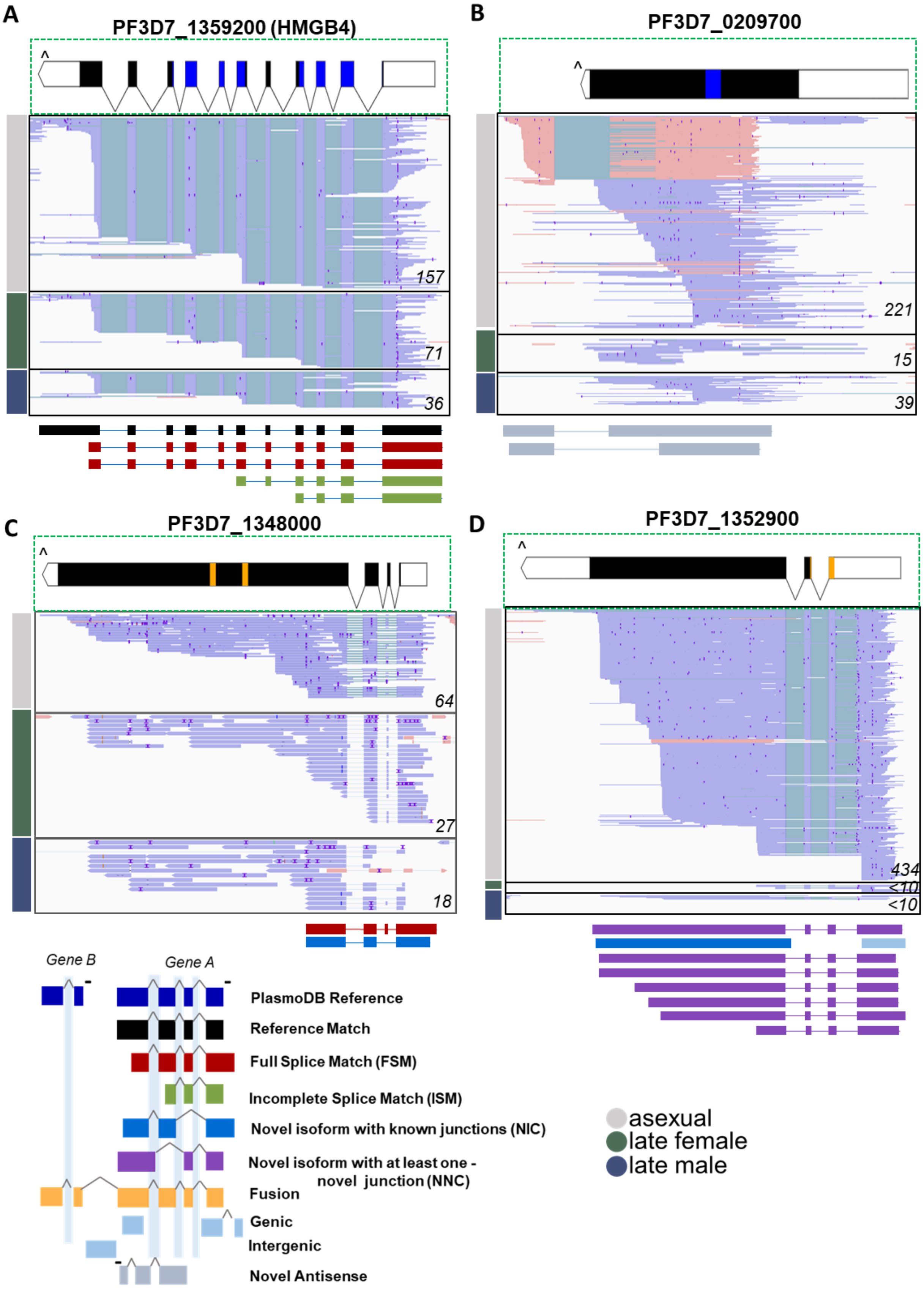
A
PF3D7_1359200 (HMGB4)
157
71
36
B
PF3D7_0209700
221
15
39
C
PF3D7_1348000
64
27
18
D
PF3D7_1352900
434
<10
<10
Gene B
Gene A
PlasmoDB Reference
Reference Match
Full Splice Match (FSM)
Incomplete Splice Match (ISM)
Novel isoform with known junctions (NIC)
Novel isoform with at least one - novel junction (NNC)
Fusion
Genic
Intergenic
Novel Antisense
asexual
late female
late male

**Fig. S12. Examples of structural categories among which SQANTI3 classifies the identified isoforms.** ***(A)*** *SQANTI3 classifies the long reads associated with HMGB4 into an exact reference match (that matches the 5' and 3' ends and splice sites), 2 full splice matches displaying both alternate 5'/3' ends, and 2 incomplete splice matches. While the full splice matches with a shorter 5' UTR, and the 2 incomplete splice matches could be real isoforms, we cannot exclude the possibility of them being products of 3' degradation.* ***(B)*** *Novel isoforms antisense to PF3D7_1209400 are detected and classified accordingly by SQANTI3.* ***(C)*** *A full splice match with a short 3' end and an novel isoform containing a combination of known splice sites are identified for PF3D7_1348000.* ***(D)*** *A novel splice site is identified in the first exon of PF3D7_1352900 within the 5' UTR and is classified among several isoforms as Novel isoforms with at least one novel junction. For each example above, IGV tracks of long read data from asexual (ring, trophozoite, schizont), late female and late male stages are shown in separate tracks, along with the PlasmoDB gene annotation (top panel) and SQANTI3 generated isoforms (bottom panel). Different cell types are indicated by coloured bars on the left. InterPro PFAM domain regions in the gene models (High mobility group box domain and Transcription factor CBF/NF-Y/archaeal histone domains of HMGB4, zf-C3HC4_2 domain of PF3D7_0209700) are displayed in blue. Transmembrane domains of PF3D7_1348000 and signal peptide of PF3D7_1352900 are displayed in orange. Reads in forward orientation are coloured rose and reads in reverse orientation in periwinkle. 3' ends of the gene models are denoted by a '^' sign.*

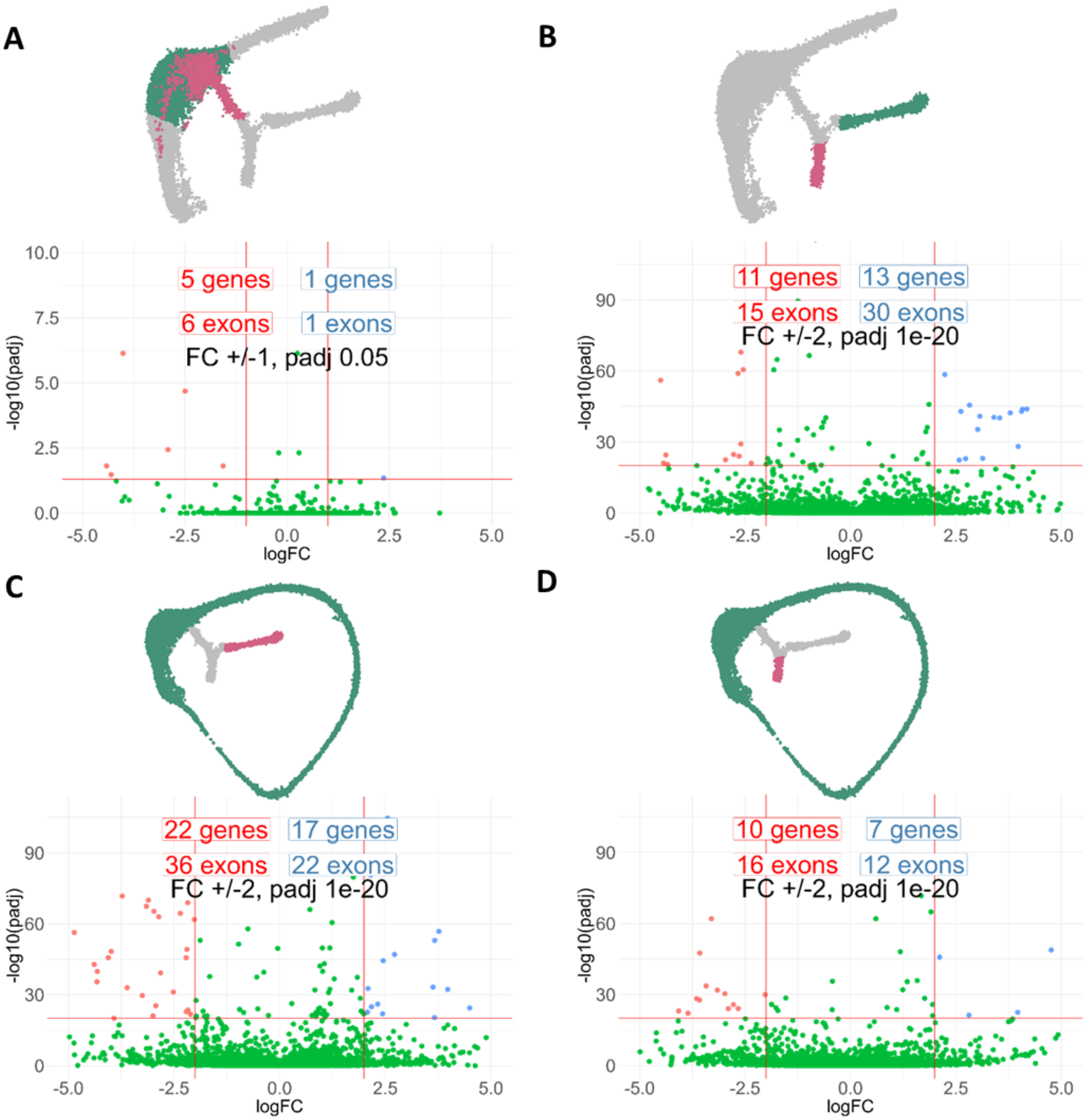


**Fig. S13. Differential exon usage between cell types was explored using DEXseq *(A)*** *Volcano plot showing the 6 genes that have exons that change expression significantly (fold change > 1, padj < 0.05) between committed and early trophozoites.* ***(B)*** *Volcano plot showing the 24 genes (23 unique) that have exons that change expression significantly (fold change > 2, padj < 1e-20) between the cells of the male and female branch.* ***(C)*** *Volcano plot showing the 39 genes (35 unique) that have exons that change expression significantly (fold change > 2, padj < 1e-20) between the cells of the female branch and asexual stages.* ***(D)*** *Volcano plot showing the 17 genes that have exons that change expression significantly (fold change > 2, padj < 1e-20)*

*between the cells of the male branch and asexual stages. Complete list of genes from DEXseq output for the above comparisons are provided in supplementary file data S8.*

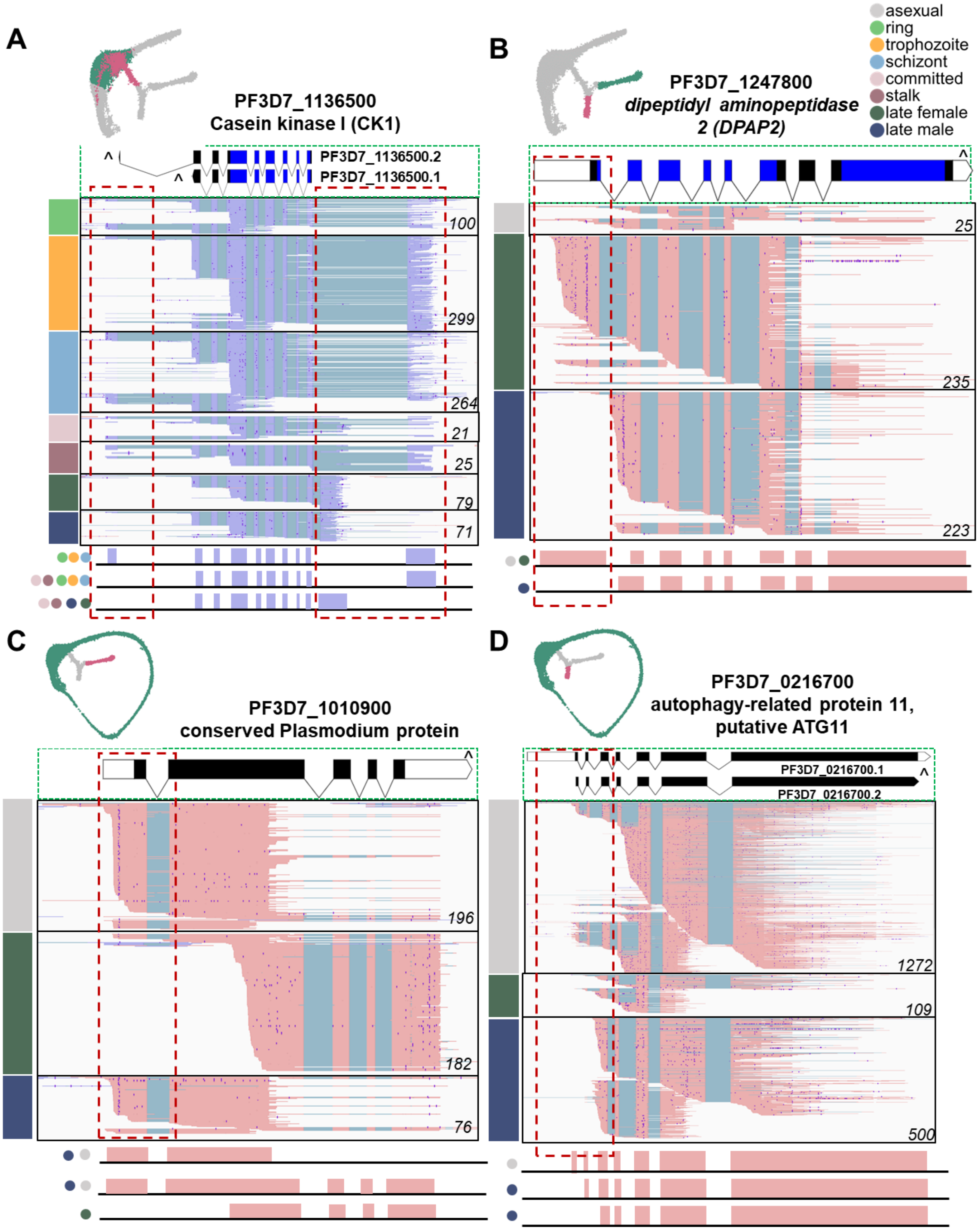


**Fig S14. Differential exon usage along sexual development.** *Here we show an example of differential exon usage for each comparison as represented in the UMAP plots and in fig. S13.* ***(A)*** *Early trophozoites vs early sexual stages (committed, stalk), where CK1 appears to transcribe*

*transcript .2 as the cells proceed to sexual development. We also observe a new gene model downstream of the annotated first exon (dashed box on the right).* ***(B)*** *Female (early female, late female) vs male (early male, late male) gametocytes, where male gametocytes lack usage of exon 1 of DPAP2.* ***(C)*** *Female (early female, late female) vs asexuals (ring, trophozoite, schizont), where PF3D7_1010900 displays distinct exon usage profile in females compared to both the asexuals and male gametocytes.* ***(D)*** *Male (early male, late male) vs asexuals (ring, trophozoite, schizont), where male gametocytes show no usage of the first 2 annotated exons. For each example (A-D), long reads spanning the gene in different cell types, as indicated by coloured bars on the left, are shown. IGV tracks of ring (early ring, late), trophozoite (early, late), schizont (early, late), asexual (ring, trophozoite, schizont), committed, stalk (early, late), and late female and late male stages are shown in separate tracks, along with PlasmoDB gene annotation (top panel). InterPro PFAM domain regions in the gene model (Pkinase Protein kinase domain for CK1, CathepsinC_exc Cathepsin C exclusion domain and Peptidase_C1 Peptidase C1A, papain C-terminal domain for DPAP2) are displayed in blue. Reads in forward orientation are coloured rose and reads in reverse orientation in periwinkle. 3' ends of the gene models are denoted by a '^' sign.*

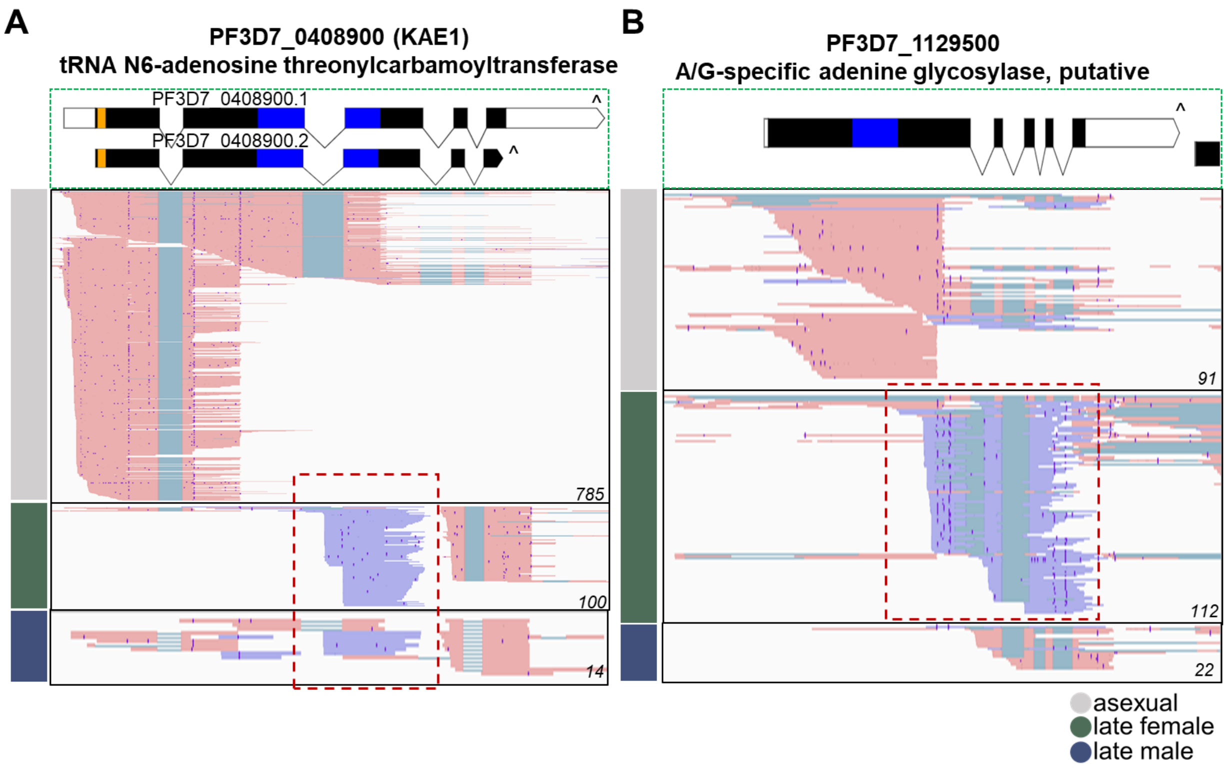


**Fig S15. Antisense expression, associated with decreased exon usage, was observed in several genes upon inspection of DEU candidates upon DEXseq analysis.** ***(A)*** *It is likely that KAE1 is in fact two genes and expression of antisense transcripts in the sexual stages is associated with decreased exon usage of the first 3 exons.* ***(B)*** *Presence of antisense transcripts in the female gametocytes is associated with decreased expression of the sense transcripts in PF3D7_1129500. Another such example is PF3D7_0213600, presented later in fig. S16. Reads in forward orientation are coloured rose and reads in reverse orientation in periwinkle. 3' ends of the gene models are denoted by a '^' sign. InterPro PFAM domain regions in the gene models (TsaD Gcp-like domain of KAE1 and HhH-GPD domain of PF3D7_1129500) are displayed in blue. Transmembrane domain of KAE1 is displayed in orange.*

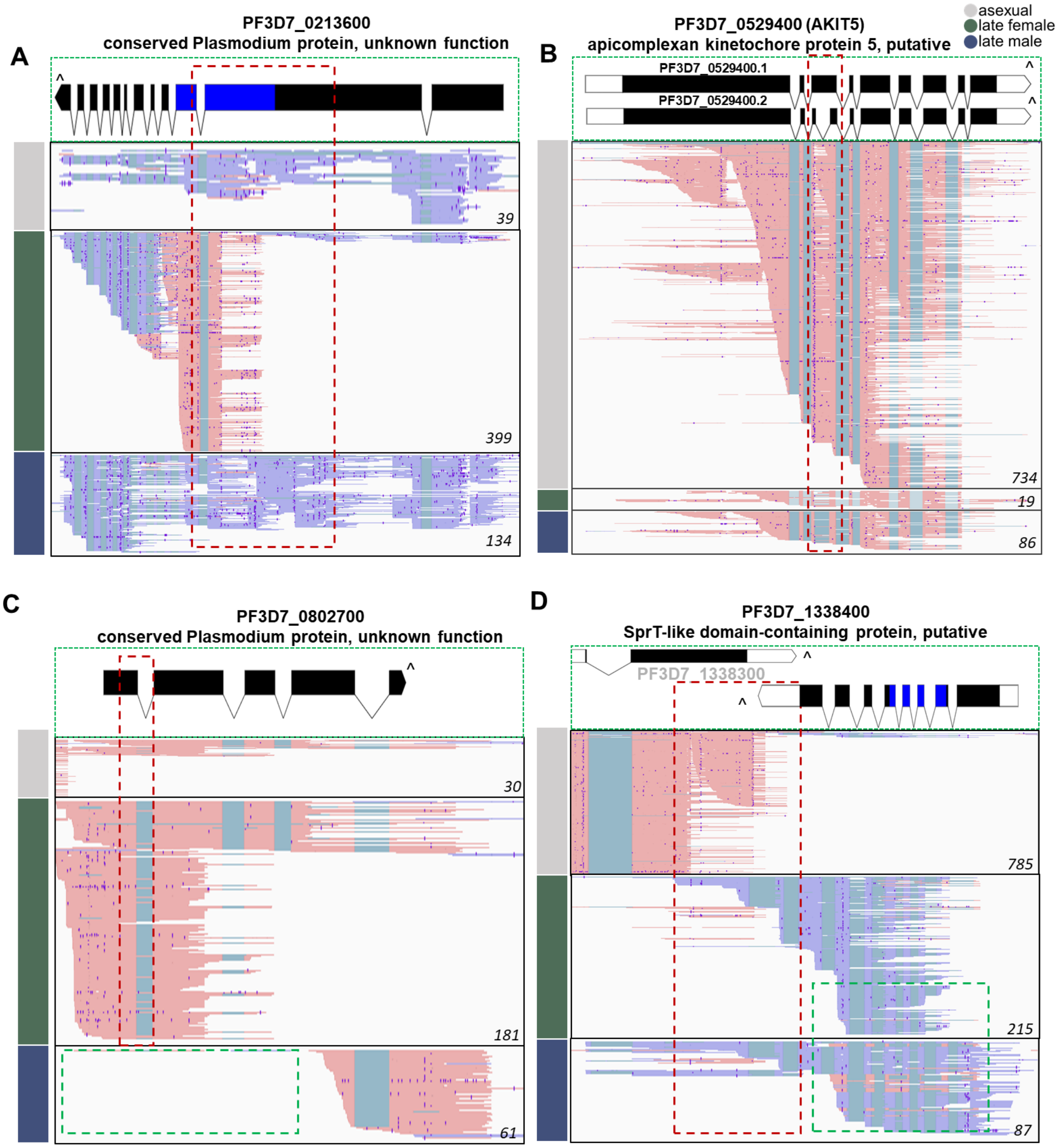


**Fig S16. Long read data helps identify novel cell-type specific splicing events and gene models.** *Following DEXseq comparisons, inspection of the significant hits revealed genes containing novel splice sites and gene models not present in the current annotations on PlasmoDB. Long reads are shown spanning the gene in different cell types as indicated by coloured bars on the left for each panel. IGV tracks of asexual (ring, trophozoite, schizont), late female, and late male stages are shown in separate tracks, along with the PlasmoDB gene annotation (top panel). InterPro domains (Panther domain LISH DOMAIN-CONTAINING PROTEIN FOPNL of PF3D7_0213600, and PFAM*

*domain SprT-like domain of PF3D7_1338400) are displayed in blue. Reads in forward orientation are coloured rose and reads in reverse orientation in periwinkle. 3' ends of the gene models are denoted by a '_' sign. Coverage values from IGV tracks are displayed in the lower right corner of each track.* ***(A)*** *A novel transcript antisense to the first four exons of PF3D7_0213600 is detected in the female gametocytes.* ***(B)*** *PF3D7_0529400 has two annotated isoforms on PlasmoDB. PF3D7_0529400.2 has an additional intron within exon 3, and this isoform appears to be specific to the sexual stages.* ***(C)*** *The first intron of PF3D7_0802700 is retained in most asexual stages and in some female gametocytes (red dashed box), while male gametocytes lack the first 3 exons of the current gene model (green dashed box).* ***(D)*** *New splice sites are observed in the last exon of PF3D7_1338400 (red dashed box). In addition, antisense transcripts with canonical splice sites are observed in the sexual stages (green dashed box).*

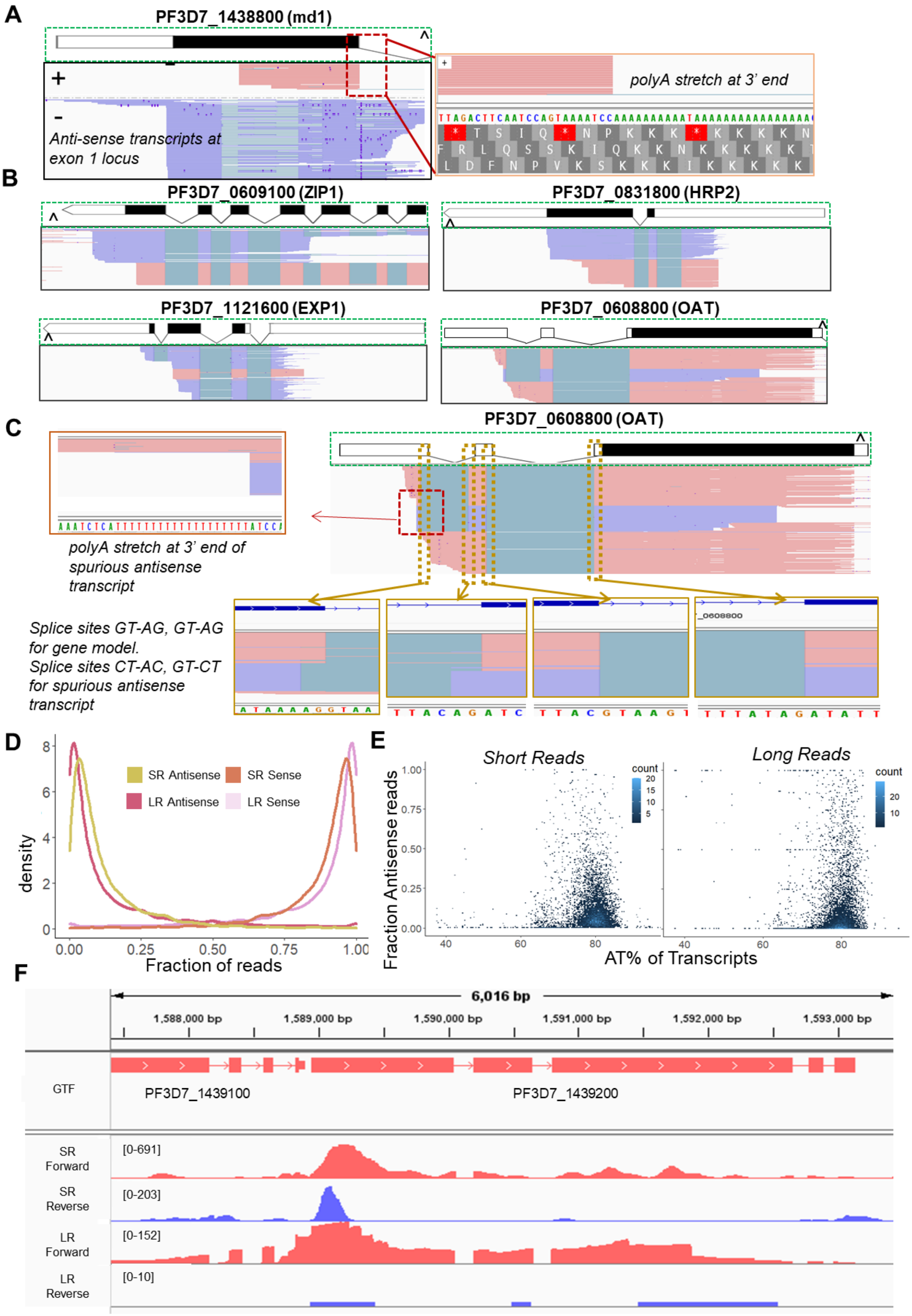
A
PF3D7_1438800 (md1)
+
-
Anti-sense transcripts at exon 1 locus
polyA stretch at 3' end
B
PF3D7_0609100 (ZIP1)
PF3D7_0831800 (HRP2)
PF3D7_1121600 (EXP1)
PF3D7_0608800 (OAT)
C
PF3D7_0608800 (OAT)
polyA stretch at 3' end of spurious antisense transcript
Splice sites GT-AG, GT-AG for gene model.
Splice sites CT-AC, GT-CT for spurious antisense transcript
D
SR Antisense
SR Sense
LR Antisense
LR Sense
density
Fraction of reads
E
Short Reads
Long Reads
count
Fraction Antisense reads
AT% of Transcripts
F
6,016 bp
1,588,000 bp
1,589,000 bp
1,590,000 bp
1,591,000 bp
1,592,000 bp
1,593,000 bp
GTF
PF3D7_1439100
PF3D7_1439200
SR Forward
[0-691]
SR Reverse
[0-203]
LR Forward
[0-152]
LR Reverse
[0-10]

**Fig. S17. As a result of the high AT% of P. falciparum, spurious transcripts may occur due to priming off poly-A stretches on first-strand cDNA.** *Examples include* ***(A)*** *The forward direction transcripts that appear in exon 1 of PF3D7_1438800 appear to be spurious transcripts, as evidenced by the polyA stretch at the 3' end as well as block-like appearance of these transcripts* ***(B)*** *Examples of other loci where these spurious transcripts appear.* ***(C)*** *We use PF3D7_0608800 as an example to show that the above transcripts don't have canonical splice sites and have a 20nt polyA stretch at the 3' end. The spurious transcripts in the other 3 examples similarly display long polyA stretches and contain non-canonical splice sites. Reads in forward orientation are coloured rose and reads in reverse orientation in periwinkle. 3' ends of the gene models are denoted by a '^' sign.* ***(D)*** *Proportion of reads mapping antisense to current annotated genes is slightly higher in short read data. Mapping reads to the current reference annotation shows that a proportion of reads map antisense to the annotated genes (median of 7.2% in short read data and 5.3% in long read data), with a slightly higher fraction in the short read dataset compared to the long read. SR - Short read, LR - Long read. Counts on genes (only those with non-zero reads in both datasets) were generated in the sense and antisense orientation using htseq-count, with "-m union --nonunique none" option. It is likely that a proportion of these antisense reads arise from real, unannotated coding and non-coding transcripts.* ***(E)*** *The proportion of reads mapping antisense to annotated transcripts was explored in relation to the AT% of the transcripts. The density of anti-sense mapping appears to be higher in short read data compared to long read data, as seen with the shift of blue density.* ***(F)*** *IGV snapshot showing the forward and reverse tracks of short and long reads from male gametocytes (early, late) of PF3D7_1439200. This gene is used as an example where the proportion of reads in the opposite orientation of the gene is more prominent in the short read dataset compared to the long read dataset.*

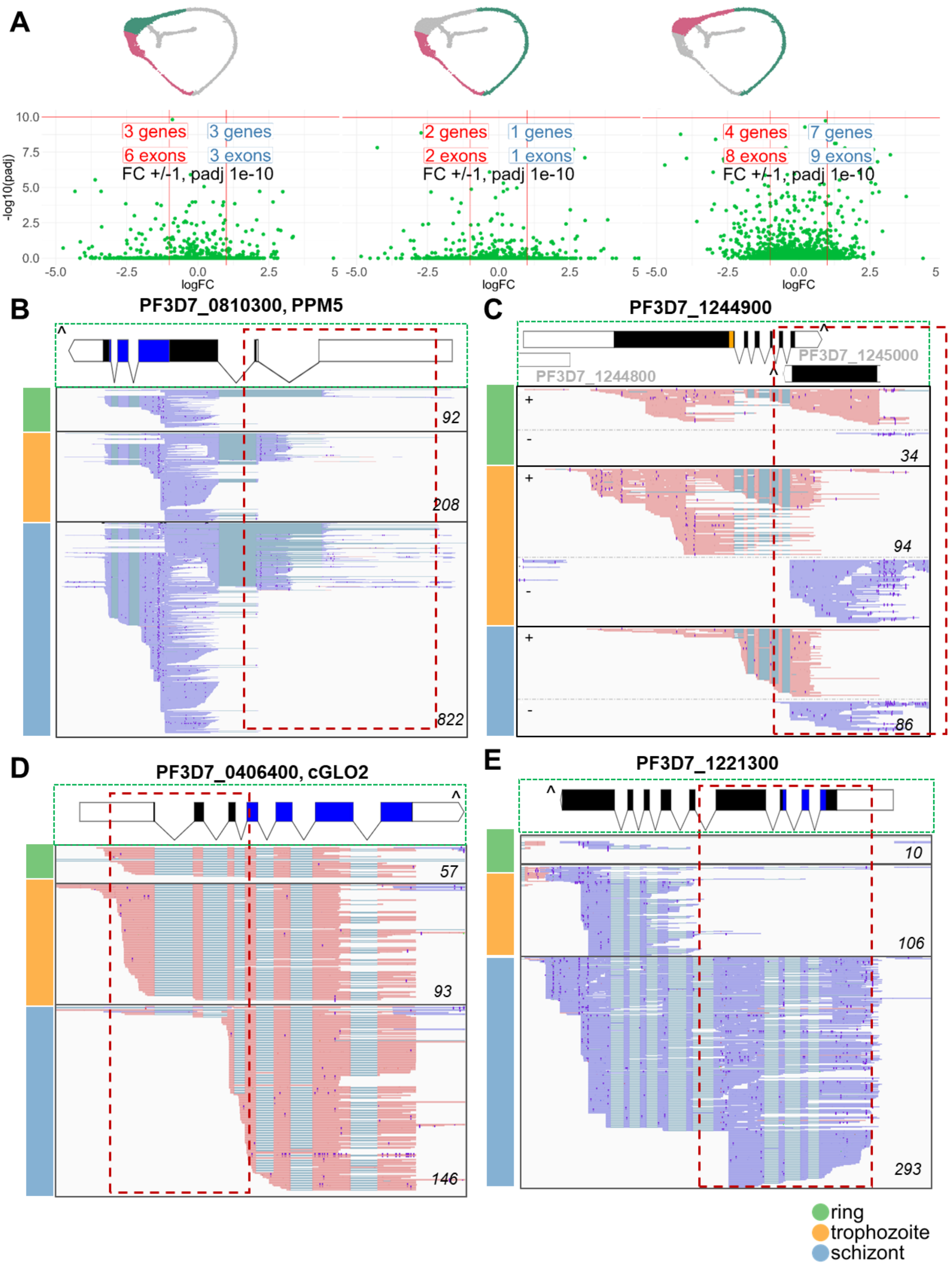

A
10.0
7.5
5.0
2.5
0.0
-log10(padj)
3 genes
6 exons
3 genes
3 exons
FC +/-1, padj 1e-10
2 genes
2 exons
1 genes
1 exons
FC +/-1, padj 1e-10
4 genes
8 exons
7 genes
9 exons
FC +/-1, padj 1e-10
-5.0
-2.5
0.0
2.5
5
logFC
B
PF3D7_0810300, PPM5
92
208
822
C
PF3D7_1244900
PF3D7_1245000
PF3D7_1244800
34
94
86
D
PF3D7_0406400, cGLO2
57
93
146
E
PF3D7_1221300
10
106
293
ring
trophozoite
schizont

**Fig S18. Long read data identifies novel stage-specific splicing events and gene models among asexual stages.** ***(A)*** *Volcano plots showing the output of differential exon usage analysis between ring vs trophozoite, ring vs schizont, and trophozoite vs schizont. Complete list of genes from DEXseq output for the above comparisons are provided in supplementary file data S8.* ***(B)*** *DEXseq comparison of ring stages with trophozoites revealed PPM5 to display different splicing patterns across the asexual stages in the loci of the first 2 exons.* ***(C)*** *PF3D7_1244900, a candidate gene upon comparison of rings with schizonts shows reads with extended 3' UTR in rings. In trophozoites and schizonts, a shorter UTR is accompanied by expression of PF3D7_1245000 on the reverse strand.* ***(D)*** *cGLO2, a candidate gene upon comparison of trophozoites with schizonts, lacks the first 2 exons in schizonts.* ***(E)*** *PF3D7_1221300, another candidate gene upon comparison of trophozoites with schizonts shows minimal usage of the first 4 exons in trophozoites. Long reads spanning the gene in different cell types as indicated by coloured bars on the left, are shown. IGV tracks of ring (early ring, late ring), trophozoite (early trophozoite, late trophozoite), and schizont (early schizont, late schizont) stages are shown in separate tracks, along with PlasmoDB gene annotation (top panel). InterPro domains (PPM-type phosphatase domain of PPM5, Lactamase_B Metallo-beta-lactamase and Hydroxyacylglutathione hydrolase C-terminal domain of cGLO2 and EF-hand domain pair of PF3D7_1221300) are displayed in blue. Reads in forward orientation are coloured rose and reads in reverse orientation in periwinkle. 3' ends of the gene models are denoted by a '^' sign. Coverage values from IGV tracks are displayed in the lower right corner of each track.*

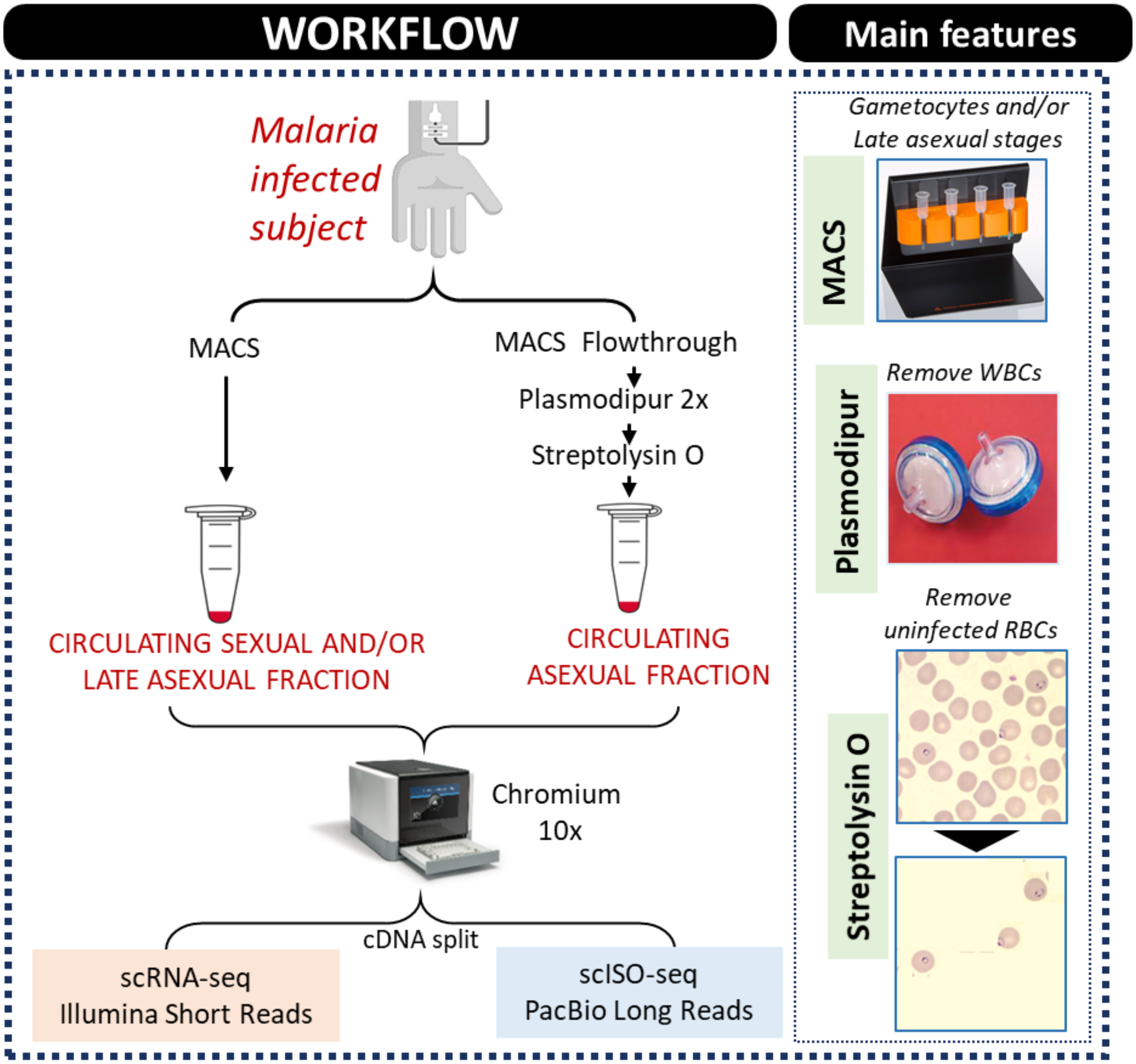


**Fig. S19. Schematic of the protocol for parasite enrichment.** *The schematic of the protocol for isolation of circulating intraerythrocytic stages of Plasmodium from malaria infected subjects. A detailed protocol is available at protocols.io (74)*

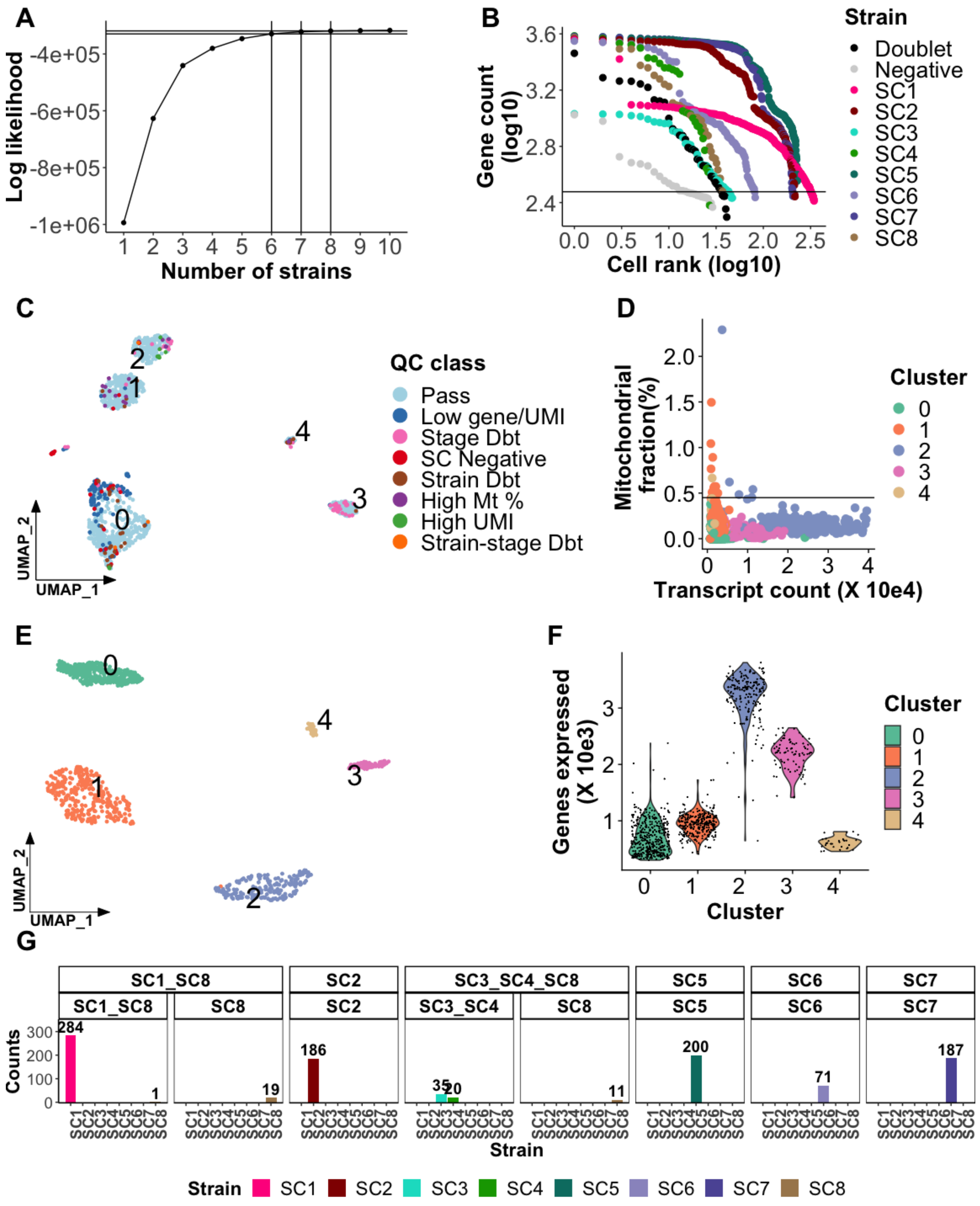


**Fig. S20. Genotype and gene expression profiling quality control (QC)** ***(A)*** *Elbow plot to identify the most likely number of strains among the parasites that pass cellranger QC (N=1258) using souporcell. The estimated total log likelihood is on the y axis and the number of strains (k) assumed*

*per iteration on the x axis. The number of strains just before the elbow plot plateaus is chosen as the optimum number of strains. Here, the log likelihood plateaus between k=6 and k=8 (Table S2). We used the upper classification of k=8 to get the most resolution.* ***(B)*** *Knee plot of the 1258 parasites coloured by the souporcell k=8 classification and ranked in decreasing order (x-axis) of their log transformed gene counts (y-axis). Doublets indicate droplets/barcodes with more than one parasite strain and negative are droplets/barcodes estimated by souporcell to contain mostly ambient RNA that leak into the cell suspension from cells that lyse during the cell preparation or fractions of cells that have lost most of their transcripts. Here all the cells identified to be doublets and negative by k=6, k=7 and k=8 souporcell iterations are included the 'Doublet' and 'Negative' classifications, respectively. These are removed as part of QC as described below. The other singlet parasites are coloured by their k=8 souporcell assignment. The horizontal line indicates a cut-off for removing cells with less than 300 genes, also removed as part of QC.* ***(C)*** *UMAP of gene expression (transcriptome) clusters of the 1258 parasites that pass cellranger QC. Here we highlight cells that were removed for various QC reasons; 'SC Negative' (Souporcell negative) cells, (N=29), also coloured gray in (B), 'High gene/UMI' cells, (N=9), with more than 40000 transcript (UMI) molecules that seem to be outliers, 'Low gene/UMI' cells, (N=107), expressing fewer than 300 genes or fewer than 700 transcript (UMI) molecules suspected to be of poor quality. The lower threshold of 700 was less stringent than that used for the V3 lab dataset (1000) because downstream processing indicated these to be good quality cells. We also removed outlier cells that had more than 0.45% mitochondrial gene expression relative to total expression (n=17). We finally removed doublets including; 'Strain Dbt' , inter-strain doublets identified by souporcell (N=29) also coloured black in (B), 'Stage Dbt' , inter-stage doublets (N=49) identified using doubletFinder with inter-strain doublets as the ground-truth and also using Scrublet and finally 'Stage-strain Dbt', intra-strain doublets that were also inter-stage (N=3).* ***(D)*** *Mitochondrial transcripts plotted as a percentage of the total transcripts (y axis) versus the number of total transcripts (x axis) with cells coloured by transcriptomic (gene expression) clusters determined by Seurat. All cells falling above the horizontal line at 0.45% fraction of mitochondrial reads were excluded from the analyses (N=17) as highlighted in QC above.* ***(E)*** *UMAP showing identified unsupervised clustering of 1014 parasites after filtration of poor quality cells described in the QC categories above.* ***(F)*** *Violin plot showing the distribution of gene expression (y axis) stratified and coloured by the seurat clusters (x axis) in the QCd parasites. These parasites were taken forward for lifecycle stage annotation.* ***(G)*** *Souporcell (k=6, k=7 and k=8) classification of genotypic clusters among QCd cells with k=6 (broadest classification) on the top facet label, k=7 (second most coarse strain classification) on the second facet label, and k=8 (the highest resolution classification) on the x axis. The counts of parasites in the different categories are on the top of each bar.*

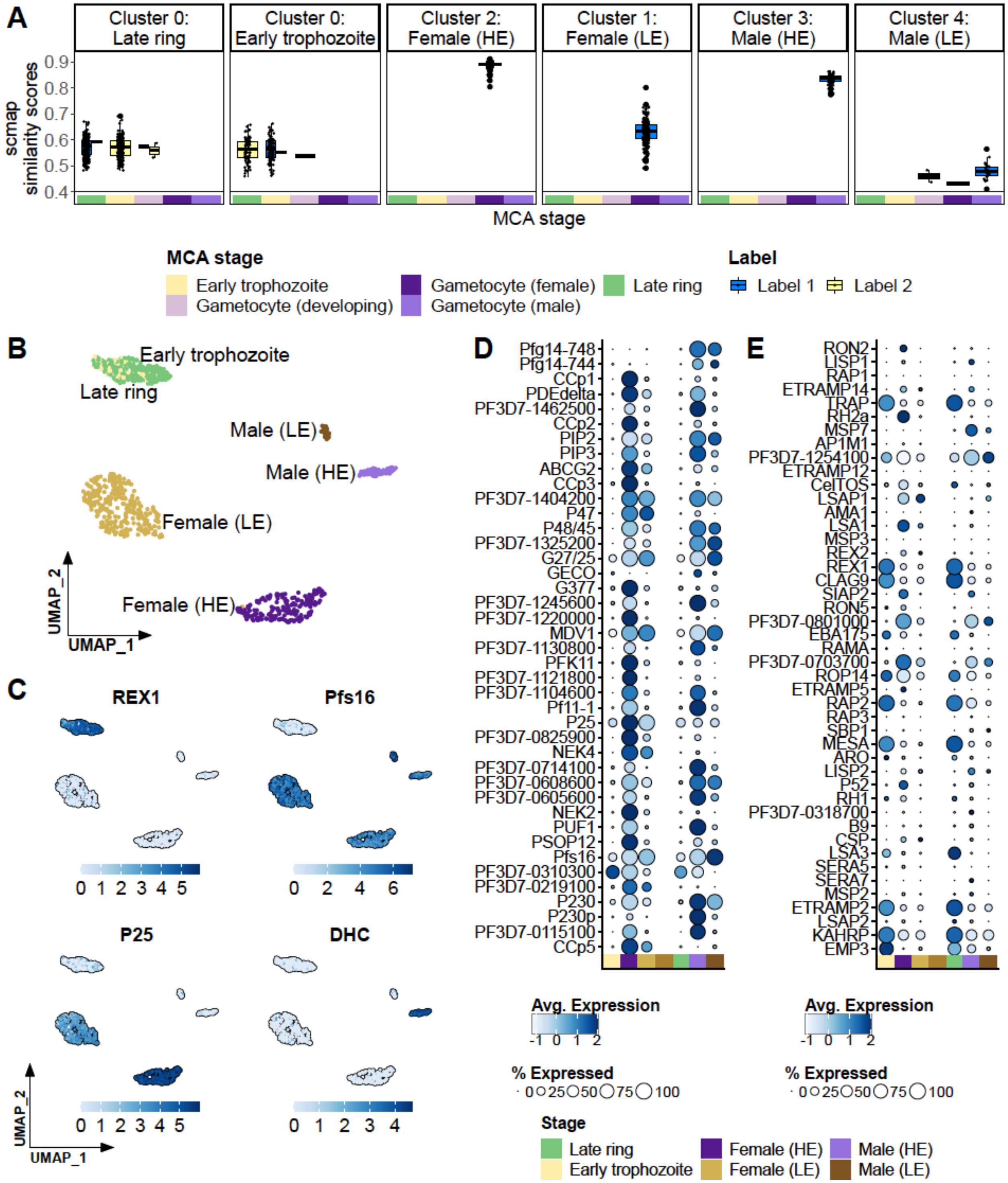


**Fig. S21. Annotation of donor parasites by label transfer from the MCA V3 reference *(A)*** *scmap label assignment for donor parasites with stage annotation used from here forward as the facet title, and the top (or the top two) scmap assignments from the MCA V3 reference on the x-axis. The similarity score of the scmap assignments is on the y-axis. The boxplots are coloured based on rank, with label1 as the top assignment and label2 as the second one. The MCA V3 reference*

*stages are represented by color blocks based on the legend below. Through this annotation process, we were able to assign stages to 1011 cells (3 cells that couldn't be annotated clearly were removed).* ***(B)*** *UMAP plot of QCd and annotated donor parasites clustered by gene expression and coloured by life-cycle stage annotation as assigned in (A).* ***(C)*** *UMAP plot similar to (B) but coloured by expression of canonical early asexual (REX1), gametocyte (Pfs16), female gametocyte (P25) and male gametocyte (Dynein heavy chain (DHC)) markers. In each plot, expression of the marker increases from light to dark blue as represented by the blue bar underneath the plot. Dotplot of asexual* ***(D)*** *and gametocyte* ***(E)*** *genes in donor parasites that have previously been reported as 'gold standard' markers from an integrated analysis of previously published P. falciparum datasets (101). Each of the marker genes is averaged across all cells per stage and indicated by a dot. The size of the dots indicates the proportion of cells within the stage that express the gene as represented by the '% Expressed' legend on the bottom right of the plot. The color intensity of the dot corresponds to the average level of expression across the cells of the particular stage increasing from light to dark blue as indicated in the color bar beneath each dot plot. The dotplot stages indicated by the color tiles on the x axis are denoted by the 'Stage' legend beneath the dotplots.*

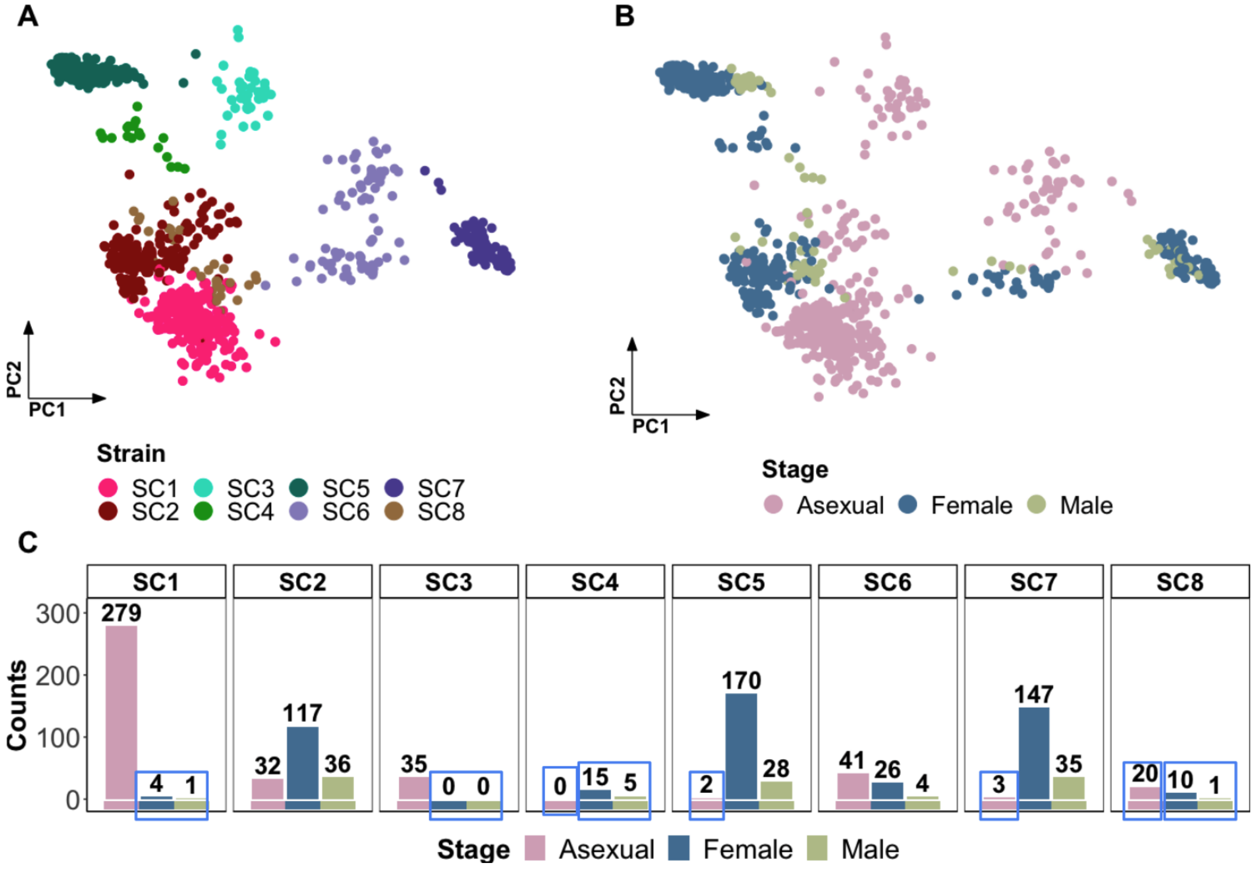


**Fig. S22. Field parasites strain and stage distribution** ***(A)*** *Genotype PCA clustering of field parasites coloured by souporcell genotype assignment.* ***(B)*** *Genotype PCA clustering of field parasites coloured by coarse stage (Late ring and early trophozoite are collapsed into Asexuals, Female (HE) and (LE) are grouped into Female, and Male (HE) and (LE) are grouped into Male assignments.* ***(C)*** *Bar plot of the counts of parasites grouped by strain and coloured by coarse-stage with the total per group indicated on top of each bar. Blue boxes indicate asexual or gametocyte (Female + Male) stage groups that contain fewer than 30 parasites and therefore removed from strain relatedness (Fig. S24,25) and differential expression (Fig.S29,30) analysis. We cannot confidently genotype stage/strain clusters with fewer than 30 cells and thus these are excluded from further analyses on IBD. The remaining groups in this bar plot were collapsed into asexual or gametocyte (Female + Male) pseudobulk groups within each strain and genotyped as independent samples.*

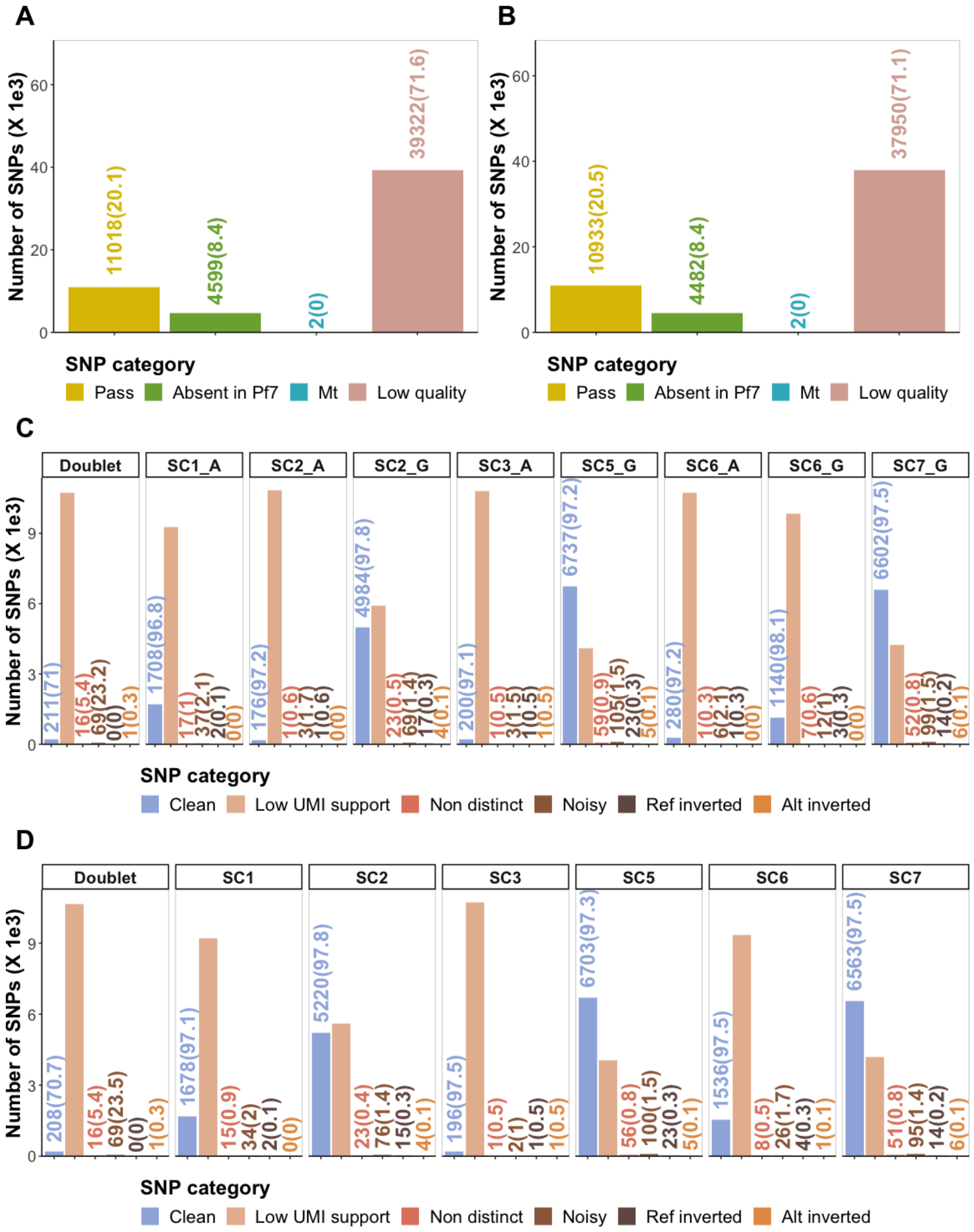


**Fig. S23. Quality control of SNPs from pseudobulk groups genotyping.** *Bar plots showing the number and proportion of SNPs coloured by different QC categories when genotyping was*

*completed on pseudo-bulks of strain and coarse-stage (asexual/gametocyte) groups (A&C) or just at the strain level (B&D). **(A&B).** Categories where the cut-off was indiscriminately applied to all variants/SNPs regardless of pseudobulk group. These include three QC filters that were applied to the initial set of biallelic SNPs (54941 for strain+coarse-stage & 53367 for strain classification) from freebayes; 1) removal of 'Low quality SNPs with genotyping quality of 75 or less & depth of 40 or less, 2) 'Mt' mitochondrial SNPs , 3) removal of 'Absent in Pf7' SNPs, i.e those are missing or failed QC among Malian variants cataloged in the pf7 dataset (85). The 'Pass' SNPs are those that qualify from this QC stage and are subjected to further QC below. C&D. Further QC with categories where the cut-offs were selectively applied to genotype calls in specific pseudobulk clusters. These included five QC filters 1) 'Low UMI support' genotypes for SNPs in pseudobulk groups where neither the reference nor the alternate allele are supported by more than 8 unique transcripts (UMIs). The numbers for this category are not shown or included in the estimation of the proportions as they are too large so as to allow focus on the other categories. 2) 'Noisy' genotype calls where the difference between the number of UMIs supporting the alternate allele and those supporting the reference allele was less than eight in the same pseudobulk group. 3) 'Non distinct' where both the alternate allele and reference allele were each supported by more than 15 UMIs in the same pseudobulk cluster. We suspected that these were RNA edits, de novo mutations or pseudo-heterozygotes (102). 4) 'Alt inverted' where freebayes calls the genotype to be alternate but vartrix estimates more UMIs to be supporting the reference allele. 5) Ref inverted' where freebayes calls the genotype to be reference but vartrix estimates more UMIs to be supporting the alternate allele. The pseudobulk group comprising cells classified as 'Doublet' by souporcell had more than 10 times the number of 'Noisy' genotypes and more than 5 times 'Non-distinct' genotypes than any other pseudobulk group for both strain+coarse-stage **(C)** and for strain **(D)** genotyping. Ignoring the 'Low UMI support', the clean genotypes comprised more than 96% in all the pseudo-bulks categories except doublets. As shown in fig.S24F below, IBD results were similar when we did a less stringent QC and filtered only 'Low quality', 'Mt' and 'Absent in Pf7' SNPs but included all the genotype calls in the other QC categories shown in the bar plots (C&D).*

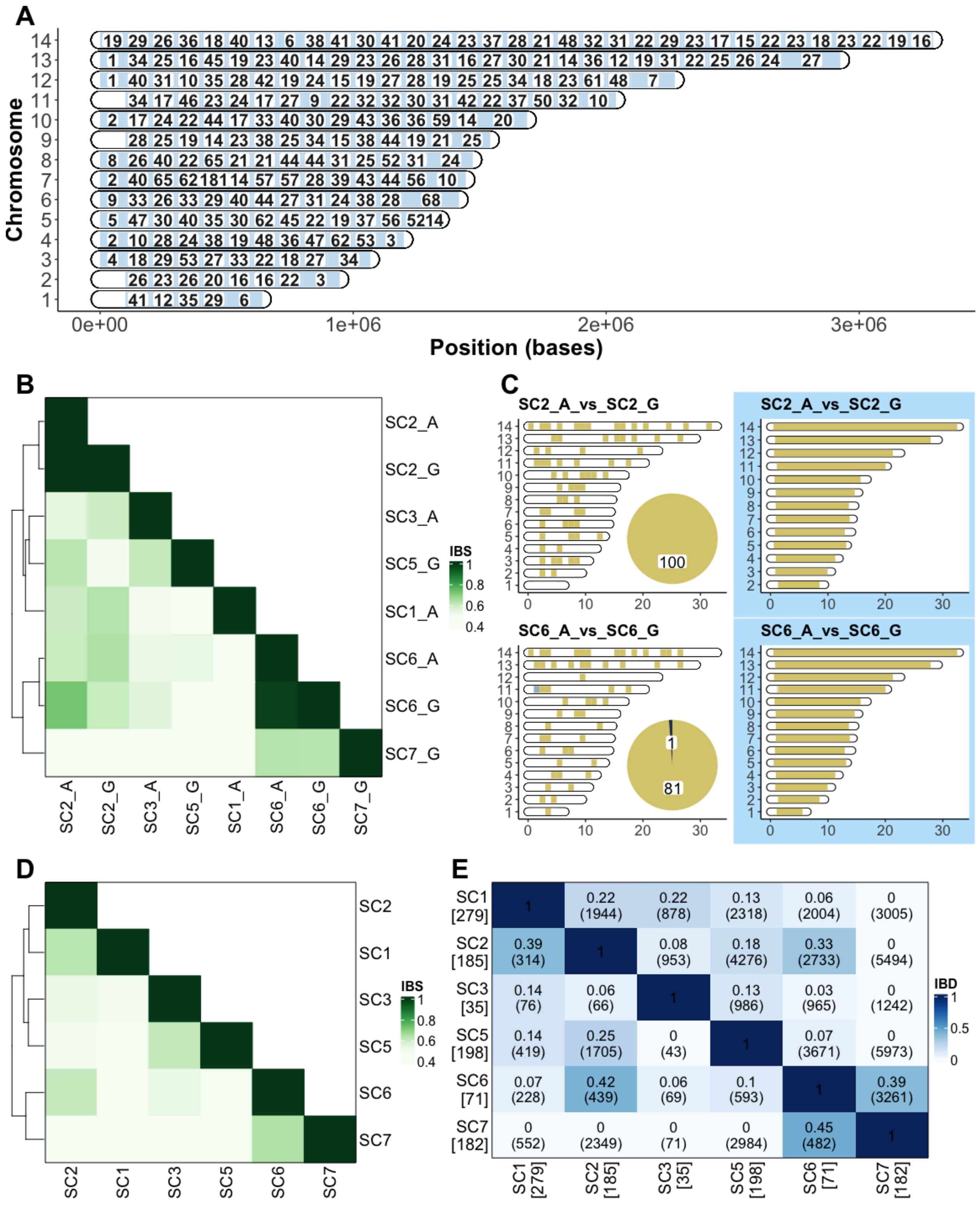


**Fig. S24. QCd genotypes of pseudo-bulked parasites grouped by stage + strain and relatedness estimates** ***(A)*** *Chromosome painting indicates the distribution of all QCd SNPs across the 14*

*chromosomes (y axis) arranged from the largest to the smallest with the position (in bases) on the x axis. Each chromosome is divided into 50 kb windows but some windows may be larger at the ends. The number of SNPs in each window is indicated (white segments have no SNPs).* ***(B)*** *Heatmap of the genome fractions that are identical by state (IBS) across all pairwise comparisons of strain/coarse-stage pseudo-bulk groups increasing from white to dark green as computed by snpRelate. The groups that had >95% of their genomes shared by IBS, highlighted in blue borders, were collapsed into one strain category. These fully recapitulate souporcell assignments.* ***(C)*** *Chromosome paint of pairwise comparison of the asexual (A) and gametocyte (G) (female + male) stages in strains that had both of these stages, i.e SC2 and SC6. For each strain comparison, the left plot with the white background is of the manual comparison of alleles within 50 kb windows across each chromosome with the segments coloured yellow if all the contained alleles are similar between the pairwise comparison, grey if some alleles are similar and others different and white if no QCd SNPs were detected. The pie chart at the bottom right of the left chromosome paint indicates the number of SNPs compared across the entire chromosome. The right chromosome paint plot with blue background shows similar pairwise comparison as on the right but with a different method, hmmIBD, where the yellow stretch indicates regions detected to be identical by descent (IBD). The hmmIBD p-values for the comparison SC2A vs SC2G and SC6A vs SC6G was 1.38e-14 and 1.43e-11, respectively.* ***(D)*** *IBS metrics increasing from white to green for pairwise comparisons between strains with genotyping done for all parasites within each strain as a pseudo-bulk.* ***(E)*** *Pairwise IBD, from hmmIBD, between strains increasing from white to blue. For each comparison in the cells, the top number indicates the IBD and the bottom number in the brackets is the number of sites (SNPs) factored in the comparison. The number of parasites per strain is indicated in square brackets together with the strain label in the row and column annotations. Bottom left contains IBD results from SNPs that have been subjected to the stringent QC steps highlighted in the methods and Fig S23 adobe. Top right quadrant contains the same comparisons with IBD estimated without stringent QC filtering of SNPs retaining those in the 'Noisy', 'Non-distinct', 'Ref inverted' and 'Alt inverted' categories.*

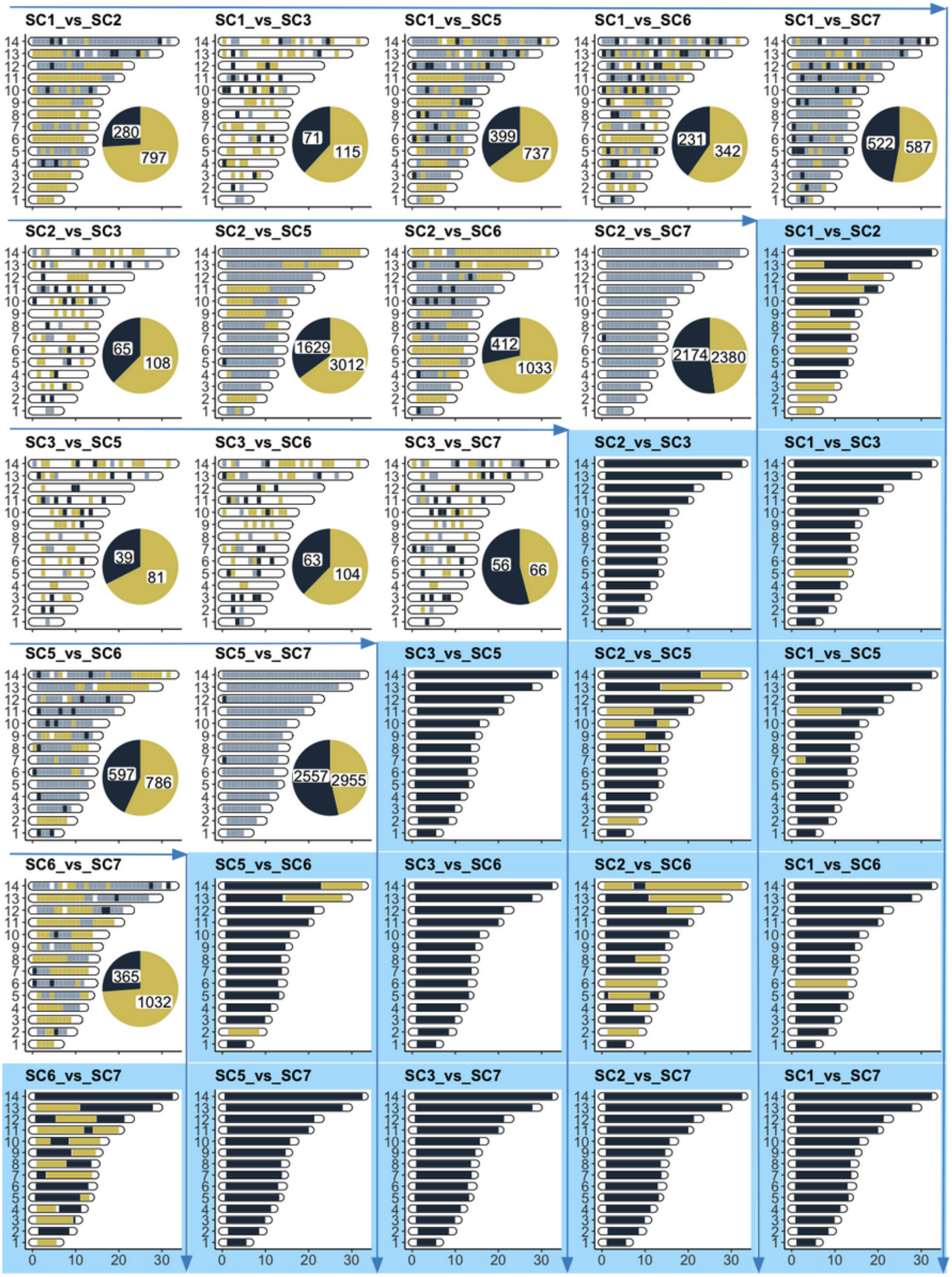
SC1_vs_SC2
280
797
SC1_vs_SC3
71
115
SC1_vs_SC5
399
737
SC1_vs_SC6
231
342
SC1_vs_SC7
522
587
SC2_vs_SC3
65
108
SC2_vs_SC5
1629
3012
SC2_vs_SC6
412
1033
SC2_vs_SC7
2174
2380
SC3_vs_SC5
39
81
SC3_vs_SC6
63
104
SC3_vs_SC7
56
66
SC5_vs_SC6
597
786
SC5_vs_SC7
2557
2955
SC6_vs_SC7
365
1032
0 10 20 30

**Fig. S25. Pairwise strain relatedness for strains SC1, SC2, SC3, SC5, SC6, and SC7.** *In each plot, the 14 P. falciparum chromosomes are on the y-axis with genomic position in bases (x1e5) on the x-axis. The top left half of the plot with the white background shows manual chromosome paint plots for all pairwise comparisons between six strains. The bottom right half with blue background shows the hmmIBD chromosome paint plots. For the manual chromosome paint plot each chromosome is divided into 100 kb windows where overlapping SNPs between each pair of strains are compared. A window is coloured yellow if all the alleles between the compared strains are identical, black if all are different, grey if some are identical and some are different and finally, white if there are no overlapping SNPs. In the pie chart at the bottom right of each plot the proportion of identical SNPs are indicated with the yellow shade and that of different alleles with navy blue shade with the total numbers included. In the hmmIBD chromosome paint plots, yellow segments indicate regions estimated to be identical by descent (IBD) and navy blue regions are not IBD. The first row and last column has the pairwise comparison between SC1 and all other strains, the second row and second last column compares SC2 with all the other strains except SC1 and so on. The blue arrows are for easy visualization of the plots indicating the manual chromosome paints going across and their counterpart hmmIBD plots going down.*

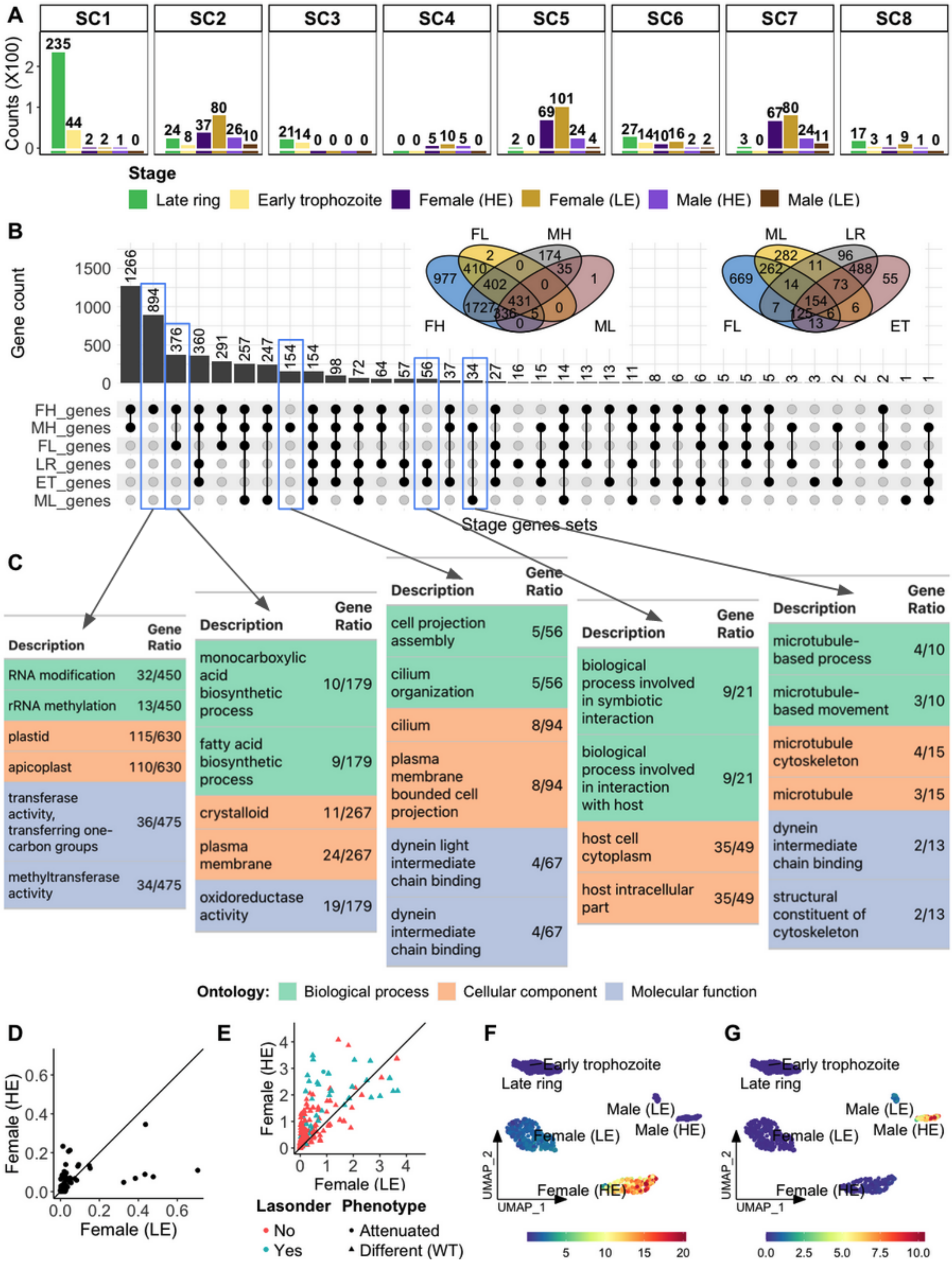


**Fig. S26. Female (HE) versus female (LE).** ***(A)*** *Barplot of the distribution of donor parasites across*

*strain and stage grouped by strain in the facets and coloured by stage at the highest resolution with the total numbers indicated for each.* ***(B)*** *UpSet plot and Venn diagrams showing overlap of 'core genes' expressed in the different stages. A gene is considered to be 'core' if it is detected in at least 25% of the cells in that stage (FH = female high, FL = female low, LR=late ring, ET=early trophozoite). On the right, the first Venn diagram indicates the overlap of all expressed genes in the gametocytes and the second Venn diagram indicates the overlap of the core genes in the LE gametocytes and asexuals. The same set in the UpSet plot and Venn diagram will not have the same number of 'core genes' because the genes are apportioned across all six stages in the former but only four stages in the latter.* ***(C)*** *Top 2 significant (adjusted p < 0.05) GO enrichment hits per ontology (cellular component, biological process and molecular function) for the five core gene sets surrounded with the blue boxes. The groups include core genes only expressed in female (HE), in both female (HE) and female (LE) which are the hallmark female genes, only in male (HE), only in asexual and in both male (HE) and male (LE), which are the hallmark male genes. In addition to the GO terms, we also include the number of genes within the GO term that are also in the gene list. The background color of the rows correspond to the GO ontology group (Biological process = green, cellular component = orange, molecular function = blue).* ***(D)*** *Correlation plot showing the ratio of unspliced to spliced read counts in female (HE) (y axis) and female (LE) (x axis) for the genes that are detected in both female (HE) and female (LE) and are also among the set of genes described to be translationally repressed in female gametocytes (22). Female (HE) ratios are higher compared to female (LE); Wilcoxon p-value = 0.028. The diagonal line cuts through x=y points.* ***(E)*** *Correlation plot showing HE (y axis) vs LE (x axis) female gene expression of genes that result in a detrimental phenotype in the mosquito stages when disrupted as reported in phenoplasm (28). Each dot represents a gene and for each gene the expression levels on the axis are the average of seurat log-normalized transcript counts in that stage. The color of the dots denote whether the genes are present or absent in the published set of transcriptionally repressed genes (22) and the shape denotes the phenotype in the mosquito stages i.e whether growth is attenuated or the parasites are different from the wild type (WT) (F) Donor parasite UMAP plot of the average expression of the raw counts of 376 genes unique to both female (HE) and female (LE) and (G) 34 genes unique to both male (HE) and male (LE). The clusters are labeled by stages with the average expression in each cell increasing from blue to red. This shows that the genes detected in either the male or female gametocyte clusters are more highly expressed in the HE clusters.*

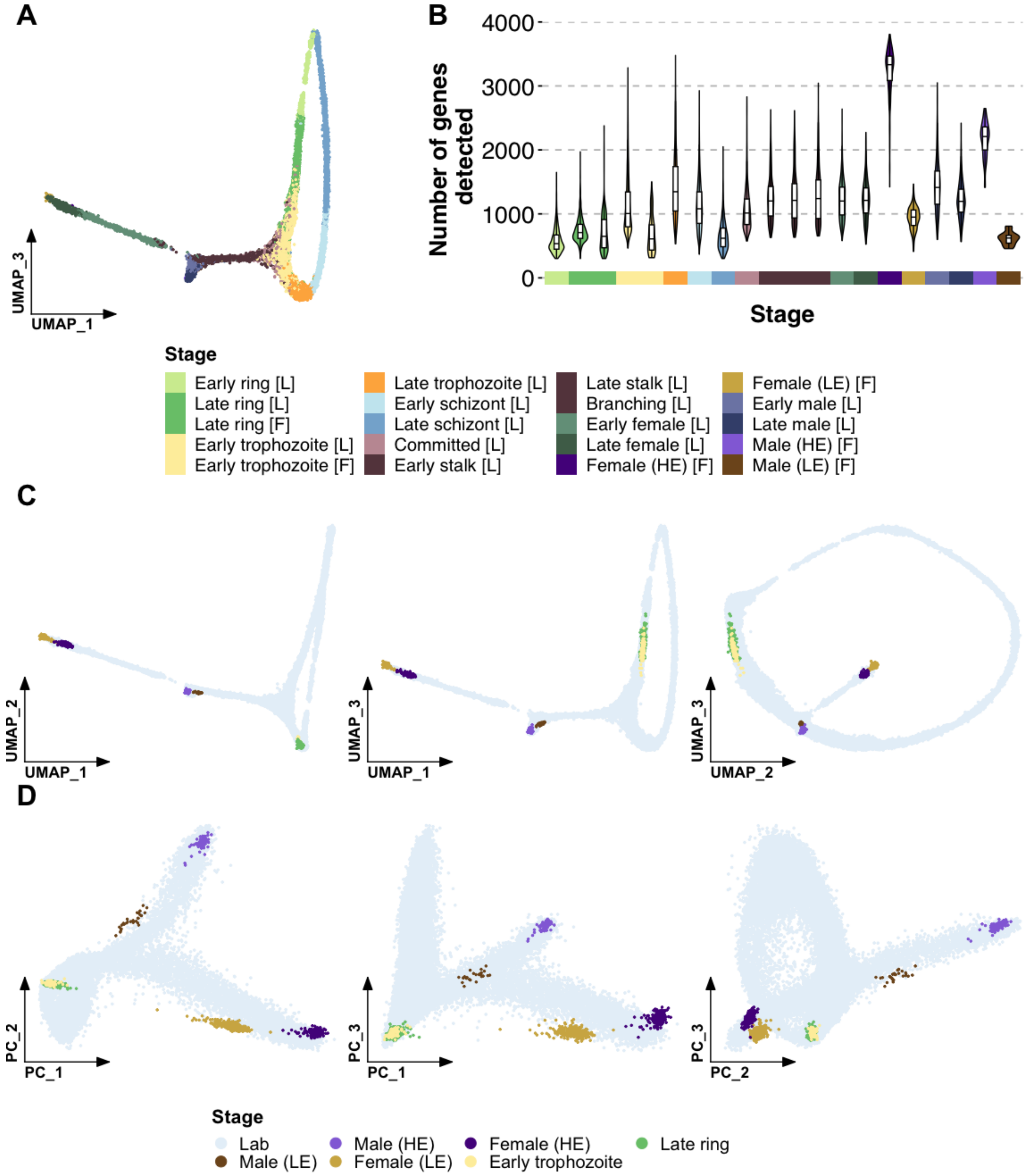


**Fig. S27. Integration of field and lab data.** ***(A)*** *UMAP plot of integrated donor parasites and MCA V3 NF54 lab parasites clustered based on gene expression with the parasites coloured by stage.* ***(B)*** *Violin plots of the distribution of genes (y axis) expressed in the parasite stages (x axis) in the integrated object. Boxplots are overlaid on the violin plots. The legend below A&B plots denote the stage colors with the letter in the square brackets indicating whether the stage is for the field, 'F' or*

*lab 'L' parasites.* ***(C)*** *Different UMAP dimensions of the same field and lab integrated object in A but showing just the field parasites.* ***(D)*** *Different dimensions of the first 3 PCA metrics for the same field and lab integrated object in A. In both the UMAP and PCA visualizations, the field parasites are overlaid on top of the lab parasites and coloured by their distinct colors and all lab parasites coloured in light blue as shown by the legend at the bottom.*

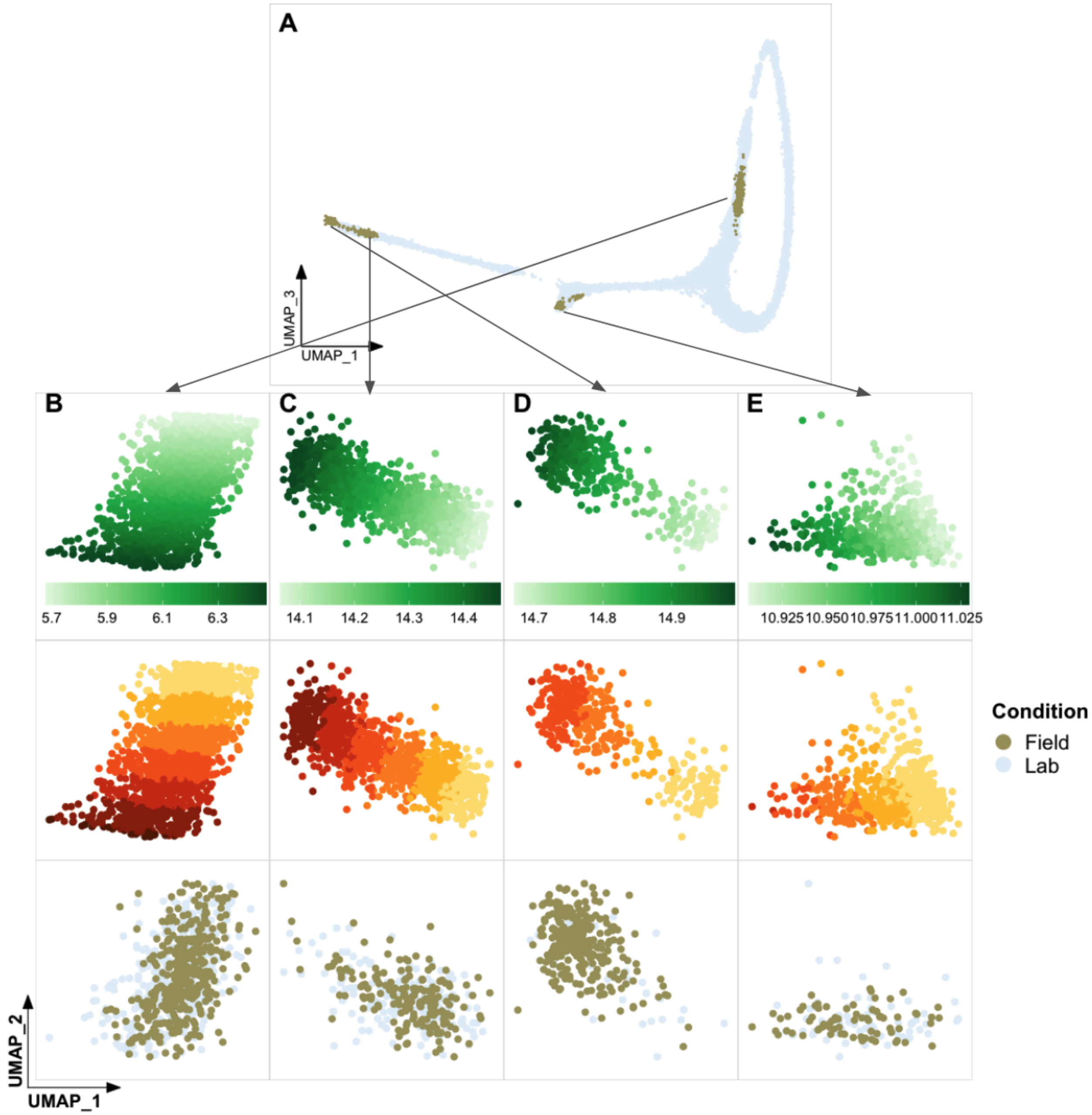


**Fig. S28. Subsetting cells for the lab vs field differential expression (DE) analysis.** ***(A)*** *Subsetting sections of integrated UMAP that contain donor parasites interspersed with lab cells colored by field and lab categories. UMAP plots of subsetted* ***(B)*** *asexual,* ***(C)*** *female (HE),* ***(D)*** *female (LE), and E) male (HE), with cells coloured by pseudotime progression (increasing from light to dark green). In the second row the cells are coloured by pseudotime chunks within which we downsample the lab parasites to be equal to the field parasites. In the third row the balanced cells with the lab downsampled are shown coloured by field and lab stage. The legend on the right indicates the color of the lab and field strains.*

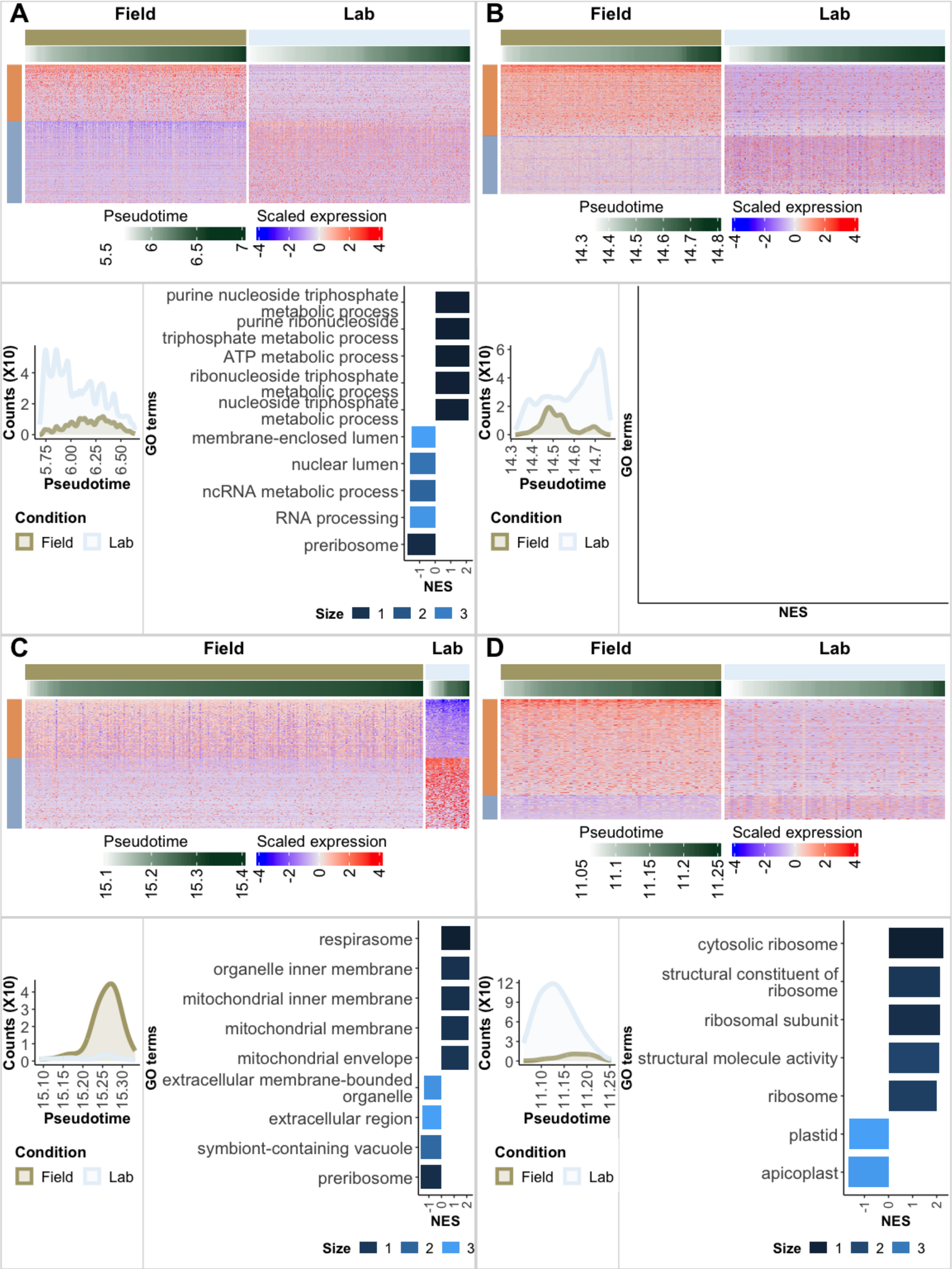
A
Field
Lab
Pseudotime
Scaled expression
B
C
D
Counts (X10)
Pseudotime
Condition
Field
Lab
GO terms
NES
Size
purine nucleoside triphosphate metabolic process
purine ribonucleoside triphosphate metabolic process
ATP metabolic process
ribonucleoside triphosphate metabolic process
nucleoside triphosphate metabolic process
membrane-enclosed lumen
nuclear lumen
ncRNA metabolic process
RNA processing
preribosome
respirasome
organelle inner membrane
mitochondrial inner membrane
mitochondrial membrane
mitochondrial envelope
extracellular membrane-bounded organelle
extracellular region
symbiont-containing vacuole
cytosolic ribosome
structural constituent of ribosome
ribosomal subunit
structural molecule activity
ribosome
plastid
apicoplast

**Fig. S29. Differential expression (DE) between lab and field parasites.** ***(A)*** *DE between field and lab parasites in asexuals [N=1054],* ***(B)*** *female (HE) [N=2836],* ***(C)*** *female (LE) [N=896], and* ***(D)*** *male (HE) [N=859] while accounting for pseudotime. In each panel the heatmap (top) details the DE, the bar plot (bottom right) shows the gene set enrichment analysis (GSEA) for GO terms (Genes highly expressed in field female (HE) parasites as compared to the lab female parasites did not yield any significant GO terms in the GSEA analysis), and the density plot (bottom left) shows the distribution of parasites along pseudotime for lab (blue) and field cells (brown). In the heatmaps, the top bar indicates the field (brown) and lab (blue) groups of parasites. Each column of the matrix within these groups represents single parasites that are arranged based on pseudotime developmental progression which is indicated by the second top bar increasing from white to dark green from left to right. Each row of the matrix is a gene and these are grouped based on the direction of expression in the field versus lab comparison as indicated on the left bar. Genes are coloured orange on the Field_vs_Lab bar if they are upregulated in the field and blue if they are upregulated in the lab. Lab parasites have been downsampled to be equal to the number of field parasites in the DE analysis. In the heatmaps, we show 1055, 2838, 903, and 836 differentially expressed genes (DEGs) in asexual, female (HE), female (LE), and male (HE), respectively. These were selected based on achieving significance in 2 statistical tests; Seurat MAST (Adjusted p-value <0.05) and tradeseq (waldStat >10). The Pearson correlation between these tests was 0.77, 0.16, 0.81 ,0.50 for asexual, female (HE), female (LE) and male (HE), respectively (P<2.2e-16). Many genes expressed in the female (HE) are not present in the lab females and may explain the low concordance in the tests. The GO terms were obtained from gene set enrichment analysis (GSEA) of the DE results of the field parasites vs the lab parasites including all detected genes. The length of the bar represents the normalized enrichment scores (NES) and the color represents the 'Size' i.e. the number of genes under the respective GO term.*

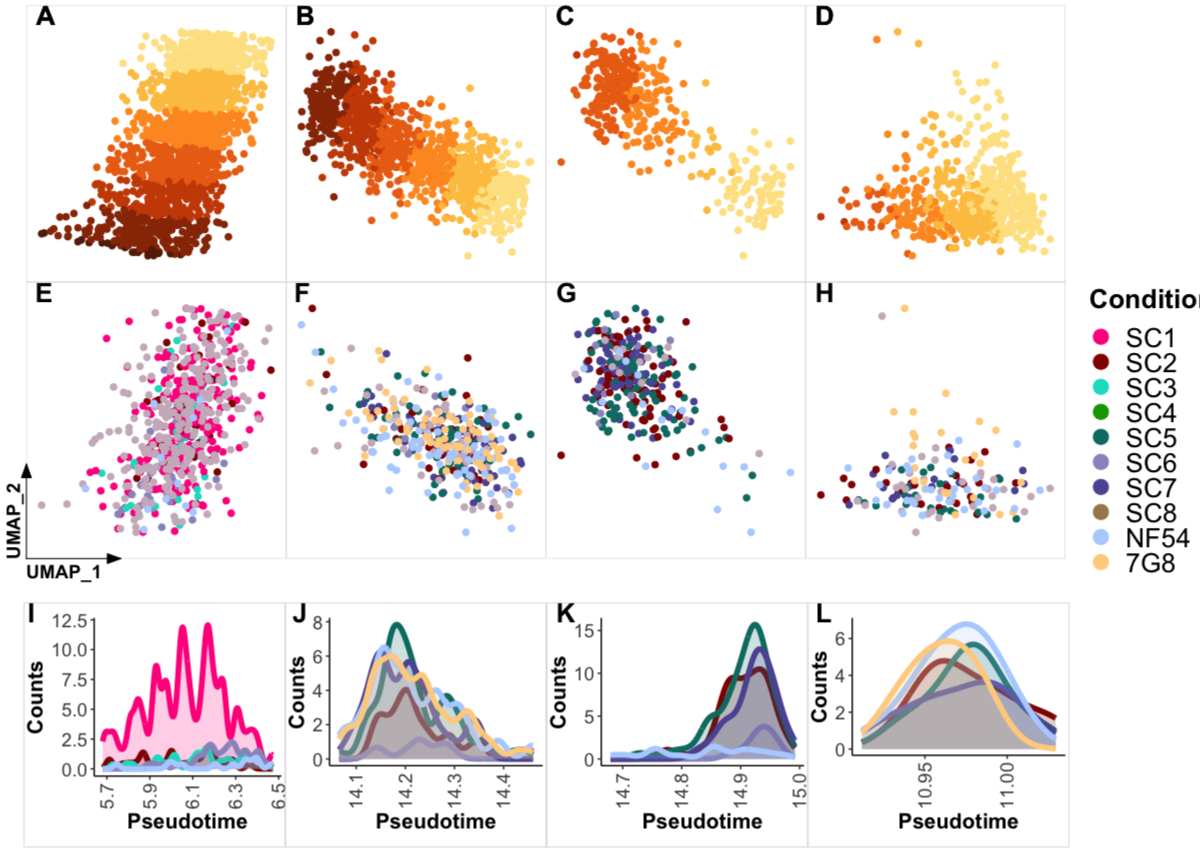


**Fig. S30. Downsampling lab parasites for comparisons of DE levels between field and lab strains.** *UMAP scatter plots of **(A)** asexual, **(B)** female (HE), **(C)** female (LE), and **(D)** male (HE) subsets coloured by pseudotime chunks within which the lab parasite strains (NF54 & 7G8) are each downsampled to be equal to the most abundant field strain. UMAP scatter plots of the distribution of the different field and lab strains after downsampling of each lab strain to be equal to the most abundant field strain showing subsets of **(E)** asexual, **(F)** female (HE), **(G)** female (LE), and **(H)** male (HE) coloured by field and lab strains. Density plots of the distribution of each strain along pseudotime after downsampling of lab parasites showing **(I)** asexual, **(J)** female (HE), **(K)** female (LE), and **(L)** male (HE) subsets coloured by strains. Asexual and female (LE) subsets had very few lab NF54 and 7G8 strains to allow DE between them so were omitted from the DE step below in which they are compared to lab strains.*

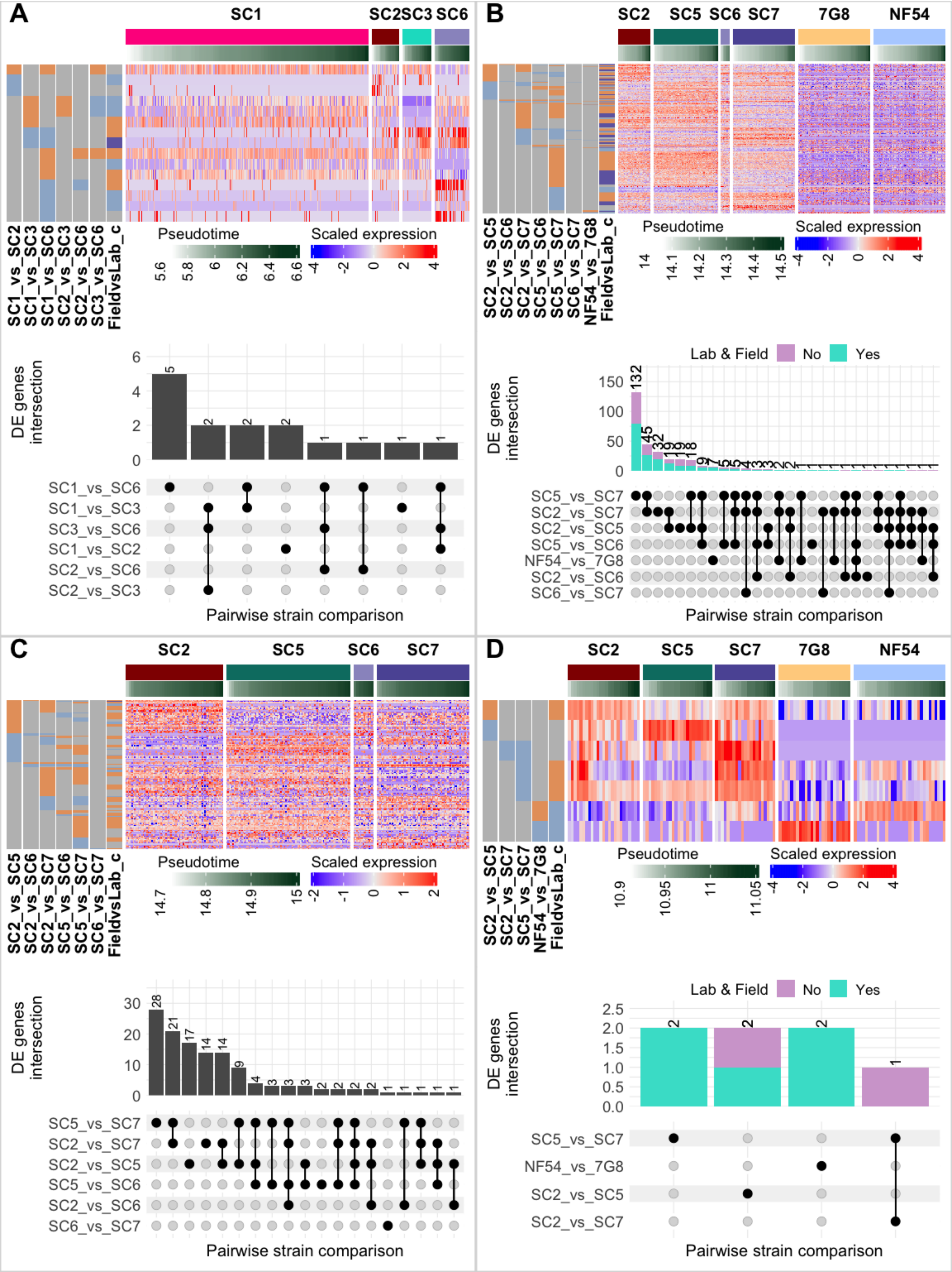
A
SC1
SC2
SC3
SC6
SC1_vs_SC2
SC1_vs_SC3
SC1_vs_SC6
SC2_vs_SC3
SC2_vs_SC6
SC3_vs_SC6
FieldvsLab_c
Pseudotime
5.6 5.8 6 6.2 6.4 6.6
Scaled expression
-4 -2 0 2 4
DE genes intersection
0 2 4 6
5 2 2 2 1 1 1 1
SC1_vs_SC6
SC1_vs_SC3
SC3_vs_SC6
SC1_vs_SC2
SC2_vs_SC6
SC2_vs_SC3
Pairwise strain comparison
B
SC2 SC5 SC6 SC7 7G8 NF54
SC2_vs_SC5
SC2_vs_SC6
SC2_vs_SC7
SC5_vs_SC6
SC5_vs_SC7
SC6_vs_SC7
NF54_vs_7G8
FieldvsLab_c
Pseudotime
14 14.1 14.2 14.3 14.4 14.5
Scaled expression
-4 -2 0 2 4
Lab & Field No Yes
DE genes intersection
0 50 100 150
132 45
SC5_vs_SC7
SC2_vs_SC7
SC2_vs_SC5
SC5_vs_SC6
NF54_vs_7G8
SC2_vs_SC6
SC6_vs_SC7
Pairwise strain comparison
C
SC2 SC5 SC6 SC7
SC2_vs_SC5
SC2_vs_SC6
SC2_vs_SC7
SC5_vs_SC6
SC5_vs_SC7
SC6_vs_SC7
FieldvsLab_c
Pseudotime
14.7 14.8 14.9 15
Scaled expression
-2 -1 0 1 2
DE genes intersection
0 10 20 30
28 21 17 14 14 9 4 3 3 3 2 2 2 2 1 1 1 1 1
SC5_vs_SC7
SC2_vs_SC7
SC2_vs_SC5
SC5_vs_SC6
SC2_vs_SC6
SC6_vs_SC7
Pairwise strain comparison
D
SC2 SC5 SC7 7G8 NF54
SC2_vs_SC5
SC2_vs_SC7
SC5_vs_SC7
NF54_vs_7G8
FieldvsLab_c
Pseudotime
10.9 10.95 11 11.05
Scaled expression
-4 -2 0 2 4
Lab & Field No Yes
DE genes intersection
0.0 0.5 1.0 1.5 2.0 2.5
2 2 2 1
SC5_vs_SC7
NF54_vs_7G8
SC2_vs_SC5
SC2_vs_SC7
Pairwise strain comparison

**Fig. S31. Differential expression (DE) between parasite strains circulating in the same host compared & between lab strains treated with the same conditions.** *Differentially expressed genes (DEGs) identified by pairwise comparisons between different parasite strains in **(A)** asexuals, **(B)** female (HE), **(C)** female (LE), and **(D)** male (HE). Donor strain (SC1,2,3,5,6 & 7) comparisons were done in different combinations in all stages but the lab strain (7G8 & NF54) comparison was only done in female (HE) (B) and male (HE) (D). This resulted in a total of 15, 318, 129 and 7 DEGs in asexuals, female (HE), female (LE), and male (HE), respectively. In each panel, the heatmap (top) shows the DE and the UpSet plot (bottom) shows the intersection of detected genes across the pairwise strain comparisons. In the heatmaps, the top bar indicates the different field and lab parasite strains. Each column of the matrix within these groups represents single parasites that are ordered by pseudotime, which is indicated by the second top bar increasing from from white to dark green from left to right. Each row of the matrix is a gene and these are grouped based on the direction of expression in the field versus lab comparison as indicated in the left bar. Genes are coloured orange on the vertical left bars if upregulated in the first named strain (eg SC1 in SC1_vs_SC6) and blue if upregulated in the second strain. On the last vertical bar on the left of each heatmap, the genes are also coloured based on whether they are differentially expressed in aggregate field versus lab comparisons as explained and determined in fig S29. In the UpSet plots, the numbers of DE genes from the different comparison sets (bottom panel) are indicated by the bars. For the female (HE) (B) and male (HE) (D) stages where we include lab strain (7G8 & NF54) comparisons, the section of bars coloured green indicate the number of genes detected in 25% of both the lab and field parasites in the comparison sets and purple indicates those failing this criteria. The DE genes are selected based on achieving significance in 2 statistical tests; Seurat MAST (Adjusted p-value $<0.05$) and tradeseq (waldStat $>10$). The pearson correlation between these tests was 0.63, 0.91, 0.89 ,0.41 for asexual, female (HE), female (LE) and male (HE, respectively ($P<2.2e-16$).*

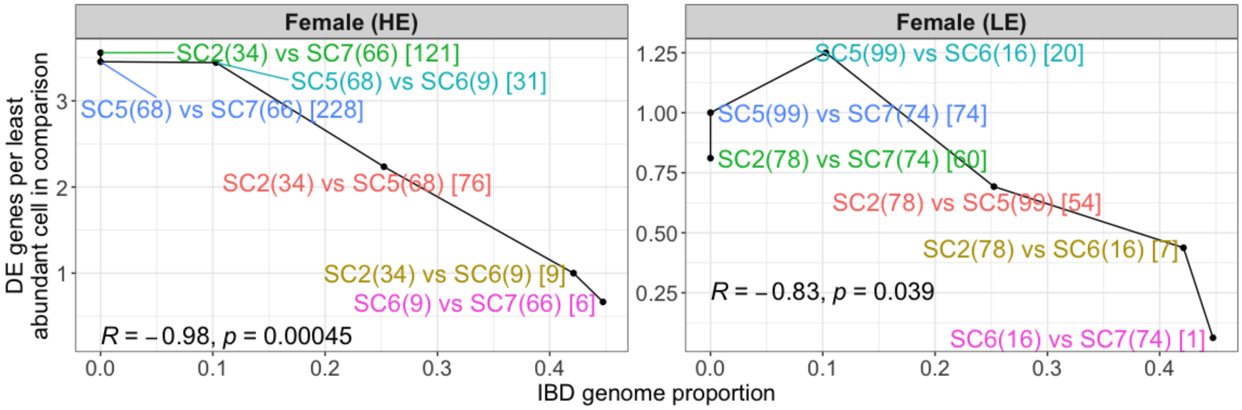


**Fig. S32. Correlation between the identity by descent (IBD) and number of DE genes.** *Here we show the relationship between the genomic relatedness distance (x axis) and the number of genes that are DE (y axis) between pairs of strains in Female (HE) and Female (LE), adjusted for the number of parasites compared by dividing by number of cells in the least abundant strain in each comparison. Each point in the plots represents a pairwise strain comparison, with the different pairs indicated and coloured differently. The number of parasites for each strain is in brackets while the total count of DE genes is in square brackets. Pearson correlation (R) together with P value (p) is indicated on the bottom left of each plot. The correlation in female (HE) is stronger probably because a lot of genes are expressed in this stage and the correlation is more robust to the fluctuation in the number of parasites being compared per strain pair. On the other hand, fewer genes are expressed in female (LE) and the correlation may not be as robust to fluctuating parasite numbers in the pairwise comparisons. This manifests in the example of SC5 vs SC6 which is the only outlier in the negative correlation trend. Here, even though the SC5 vs SC6 comparison yields fewer DE genes (n=20), normalization through dividing by 16, gives it more weight than the SC2 vs SC7 comparison which has more DE genes (n=60) and is normalized through dividing by 74.*

**A**

| ZMWs input | Polymerase Reads | Polymerase Read Length (mean) | Polymerase Read N50 | Longest Subread Length (mean) | Longest Subread N50 | HiFi Reads | HiFi Yield (bp) | HiFi Read Length (mean, bp) | HiFi Number of Passes (mean) | Non-concatamer reads with 5 and 3 primers | FLNC (Non-concatamer reads with 5' and 3' Primers and Poly-A Tail) | Number of non-redundant transcripts | Number of non-redundant transcripts, mapped to *P. falciparum* genome |
|---|---|---|---|---|---|---|---|---|---|---|---|---|---|
| 4,264,918 | 4,265,467 | 69,491 | 143,250 | 3,327 | 12,250 | 3,306,173 | 3,502,039,051 | 1,059 | 22 | 2,668,358 | 2,467,080 | 1,954,770 | 805,199 |

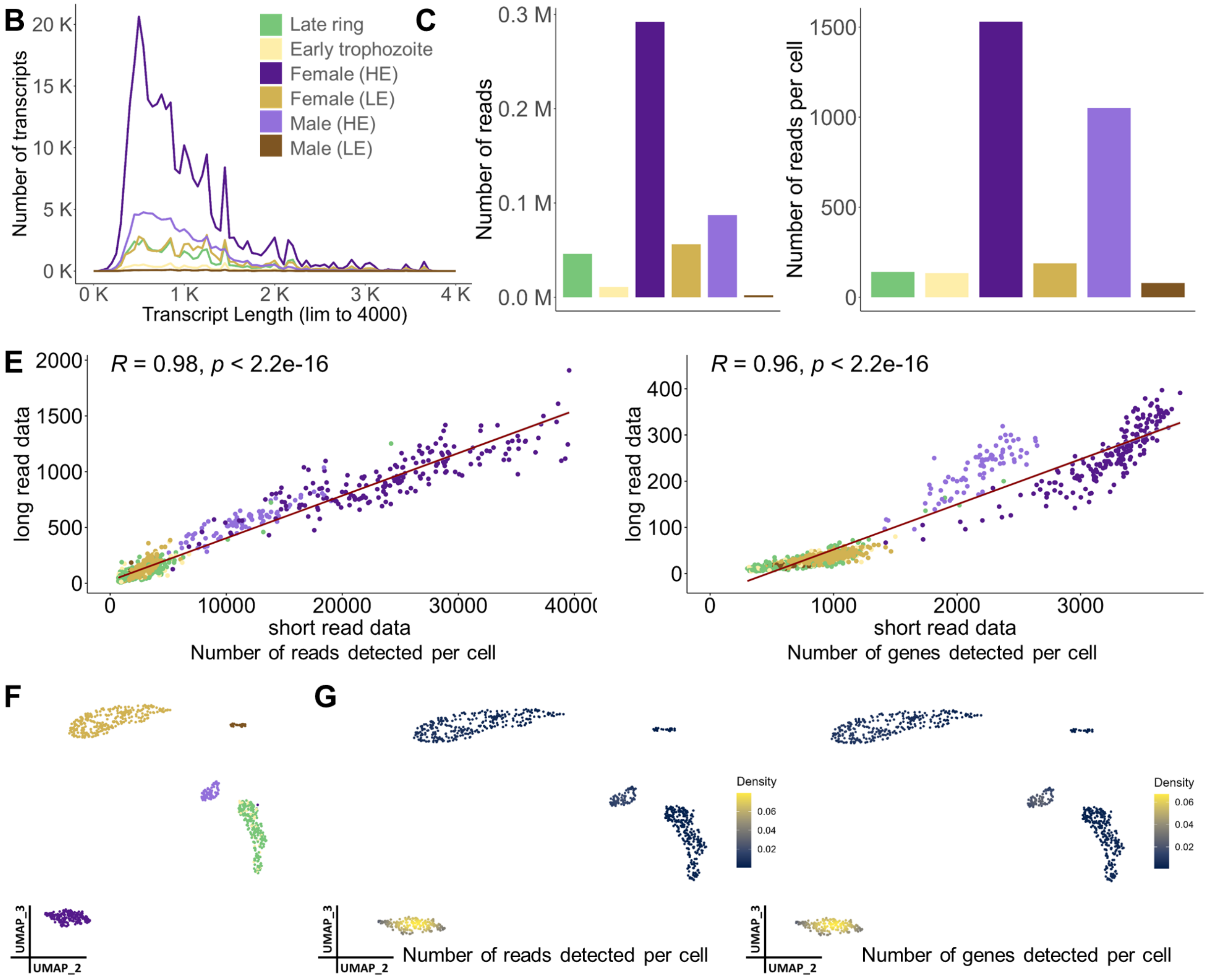


**Fig. S33. Long read sequencing output of the field sample across the detected stages.** ***(A)*** *CCS reads of donor parasites were QCd to remove primer overhangs, polyA tails, and artificial concatemers. UMI and Cell Barcode information on the long reads was used to cluster reads by the unique founder molecules, and associated metadata was assigned using short read data, including life stage assignments. Starting from ~4.2 million reads, a yield of ~2 million non-redundant reads were obtained, of which ~800,000 mapped to Plasmodium reference genome, and the rest are derived from transcripts arising from red blood cells in the sample loaded for 10x emulsion.* ***(B)*** *Transcript length distribution based on long read data, grouped by stage information assigned by*

*using short read data.* ***(C)*** *Number of long reads detected, grouped by assigned stage.* ***(D)*** *Number of long reads detected per cell, grouped by assigned stage.* ***(E)*** *Correlation of number of short reads and long reads detected per cell (left panel) and per gene (right panel).* ***(F)*** *UMAP embedding of single-cell gene expression using the long-read data of the 854 cells (which have corresponding barcodes in the filtered short read dataset), with at least 5 reads and 5 genes detected per cell. Cells are annotated with the cell identities determined from the short-read data.* ***(G)*** *Density distribution of the number of reads and number of genes detected per cell, displayed on the UMAP embedding*

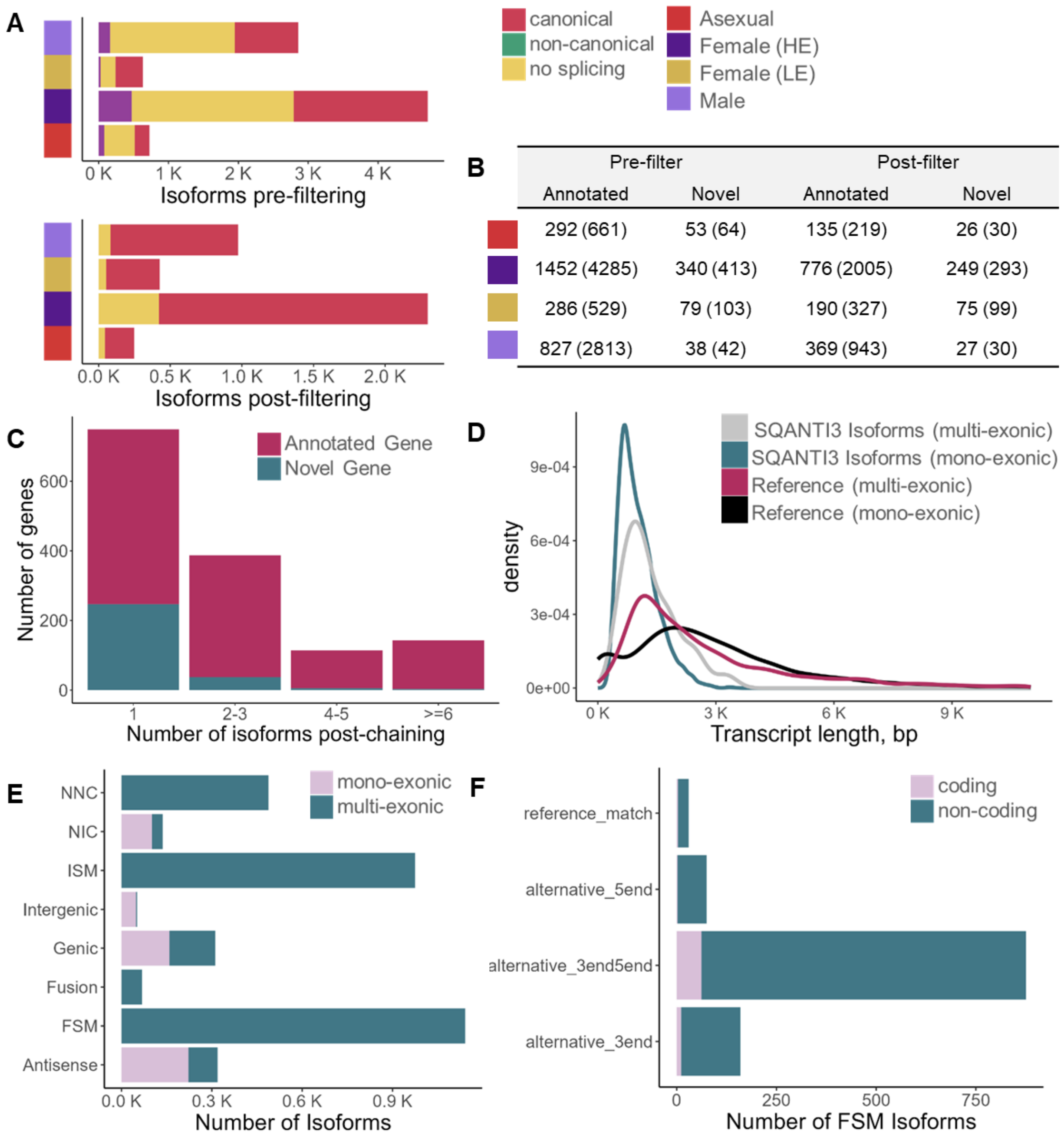


| | Pre-filter | | Post-filter | |
|---|---|---|---|---|
| | Annotated | Novel | Annotated | Novel |
| | 292 (661) | 53 (64) | 135 (219) | 26 (30) |
| | 1452 (4285) | 340 (413) | 776 (2005) | 249 (293) |
| | 286 (529) | 79 (103) | 190 (327) | 75 (99) |
| | 827 (2813) | 38 (42) | 369 (943) | 27 (30) |

**Fig. S34. Long read sequencing and corresponding stage labels from short read data allow generation of stage-specific isoforms in the field data set.** ***(A)*** *Cell type pre- and post-filtering summary of number of isoforms detected and breakdown of the splicing events detected for these isoforms. Filtering was done, similar to the laboratory datasets, to remove isoforms with > 90% adenines downstream of the transcription end site, those with non-canonical splicing junctions, and transcripts with single exons (by using sqanti3_filter.py), while retaining transcripts containing full splice matches and novel isoforms.* ***(B)*** *Cell type pre- and post-filtering summary of number of unique genes detected and the number of isoforms corresponding to these genes, classified* *according to whether they are identified as annotated or novel genes. Only the post-filtered isoforms are retained for further analysis by chaining to get non-redundant isoforms across all the stages.* ***(C)*** *Following chaining of the isoforms from all the cell types to obtain unique isoforms representing the full dataset, the plot shows the number unique genes binned according to the number of isoforms per gene, and categorised into whether they are novel genes or associated to annotated genes.* ***(D)*** *Transcript length distribution displayed as a kernel density plot for the mono-exonic and multi-exonic isoforms post-chaining, classified by SQANTI3 (mono-exonic = 1348 bp, multi-exonic = 1028 bp), compared to the 5791 reference transcripts currently annotated on PlasmoDB v59 (mono-exonic = 3144 bp, multi-exonic = 2920 bp).* ***(E)*** *SQANTI3 classification of sub-categories for the isoform events detected in the dataset post-chaining, as shown in the right panel (reproduced from https://isoseq.how/classification/categories.html, which has detailed description of these categories).* ***(F)*** *SQANTI3 classification of full splice matches into sub-categories post-chaining, showing the number of coding and non-coding transcripts among them. Among full-splice matched isoforms (1143), the majority of events are alternate 5' and 3' ends. Gene annotation files based on long reads from each cell type as well as the annotation chained by merging across the cell types is provided as a resource in data S14 and at Zenodo repository (DOI:10.5281/zenodo.8139823).*

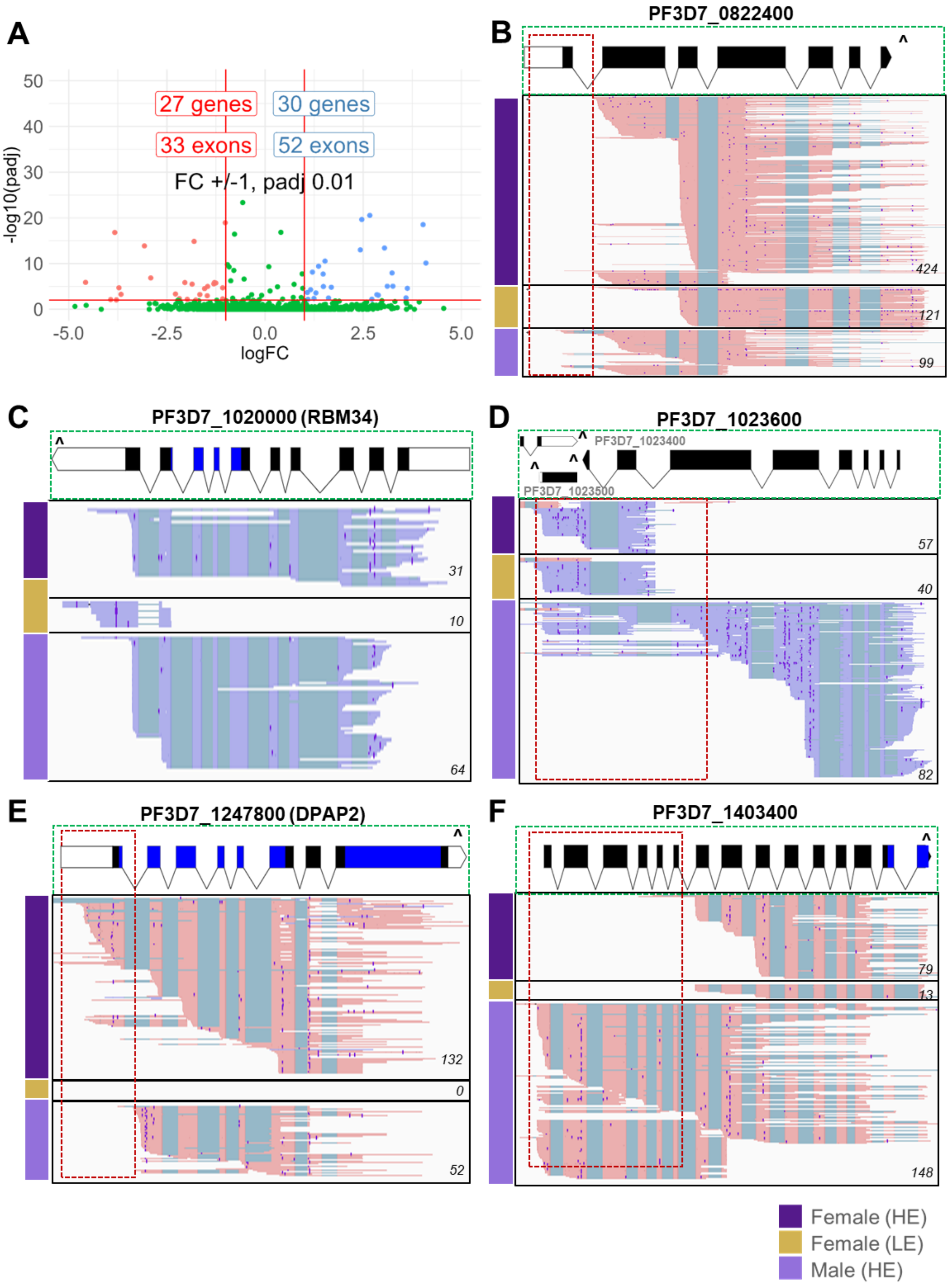
A
27 genes
30 genes
33 exons
52 exons
FC +/-1, padj 0.01
-log10(padj)
logFC
B
PF3D7_0822400
424
121
99
C
PF3D7_1020000 (RBM34)
31
10
64
D
PF3D7_1023600
PF3D7_1023400
PF3D7_1023500
57
40
82
E
PF3D7_1247800 (DPAP2)
132
0
52
F
PF3D7_1403400
79
13
148
Female (HE)
Female (LE)
Male (HE)

**Fig. S35. Differential exon usage in donor field parasites.** *Differential exon usage between female (HE) and male (HE) cells was explored using DEXseq.* ***(A)*** *The volcano plot shows the 62 genes that have exons that change expression significantly (fold change > 1, padj 0.01) between these stages. Here we show a few examples of differential exon usage. These profiles are consistent with the observations from the male and female gametocytes in the lab dataset.* ***(B)*** *PF3D7_0822400 presents exon usage profiles distinct from the gene model. A shorter isoform in female (HE) and male (HE) lacking the last 3 exons is observed, within which the female (HE) gametocytes lack usage of the first exon.* ***(C)*** *Male and female gametocytes display distinct exon usage profiles, and the annotated gene model PF3D7_1010900 could indeed be 2 genes.* ***(D)*** *For PF3D7_1023600, female gametocytes express a shorter fragment comprising the last 2 exons, while the male gametocytes show usage of all the exons in the gene model, along with the shorter fragment.* ***(E)*** *Male gametocytes lack usage of the first exon of DPAP2.* ***(F)*** *Female gametocytes lack usage of the first 6 exons of PF3D7_1403400. IGV tracks of long reads of Female (HE), Female (LE) and Male (HE) stages are shown in separate tracks, along with PlasmoDB gene annotation (top panel). Reads in forward orientation are coloured rose and reads in reverse orientation in periwinkle. 3' ends of the gene models are denoted by a '^' sign. Coverage values from IGV tracks are displayed in the lower right corner of each track. InterPro PFAM domain regions in the gene model (Papain C-terminal domain for DPAP2, OHA OST-HTH associated domain and OST-HTH/LOTUS domains of MD1) are displayed in blue. Complete list of genes from DEXseq output for the above comparison is provided in supplementary file data S15.*

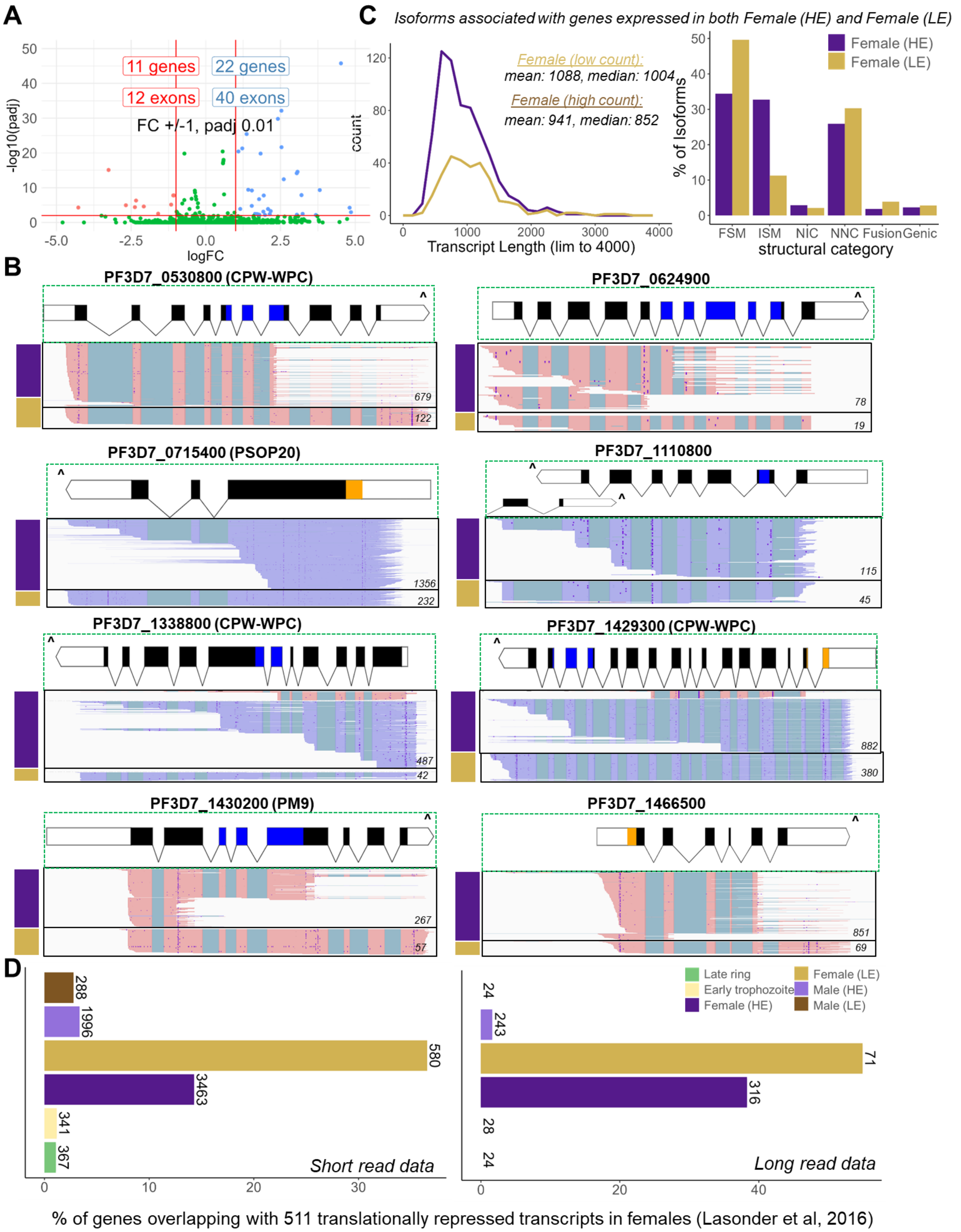

A
11 genes
22 genes
12 exons
40 exons
FC +/-1, padj 0.01
-log10(padj)
logFC
C
Isoforms associated with genes expressed in both Female (HE) and Female (LE)
Female (low count):
mean: 1088, median: 1004
Female (high count):
mean: 941, median: 852
count
Transcript Length (lim to 4000)
% of Isoforms
Female (HE)
Female (LE)
FSM
ISM
NIC
NNC
Fusion
Genic
structural category
B
PF3D7_0530800 (CPW-WPC)
PF3D7_0624900
PF3D7_0715400 (PSOP20)
PF3D7_1110800
PF3D7_1338800 (CPW-WPC)
PF3D7_1429300 (CPW-WPC)
PF3D7_1430200 (PM9)
PF3D7_1466500
D
Late ring
Early trophozoite
Female (HE)
Female (LE)
Male (HE)
Male (LE)
Short read data
Long read data
% of genes overlapping with 511 translationally repressed transcripts in females (Lasonder et al, 2016)

**Fig. S36. Female (LE) gametocytes retain longer transcripts of a subset of genes.** ***(A)*** *Differential exon usage between Female (HE) and Female (LE) was explored using DEXseq. The volcano plot shows the 11 and 22 genes that have exons that change expression significantly (fold change < -1 & > +1 respectively, padj 0.01) between these stages. Complete list of genes from DEXseq output for the above comparison is provided in supplementary file data S15.* ***(B)*** *15 of the 28 unique genes among the above show clear exon usage differences between the HE and LE females, with longer transcripts in the LE Females. We present a few examples here, which include genes translationally repressed in female gametocytes such as PSOP20 and genes of the CPW-WPC family. IGV tracks of long reads of Female (HE) and Female (LE) stages are shown in separate tracks, along with PlasmoDB gene annotation (top panel). Reads in forward orientation are coloured rose and reads in reverse orientation in periwinkle. 3' ends of the gene models are denoted by a '^' sign. Coverage values from IGV tracks are displayed in the lower right corner of each track. InterPro PFAM domain regions in the gene model (CPW-WPC domain of PF3D7_0530800, PF3D7_1338800, PF3D7_1429300, Growth arrest-specific protein 8 domain of PF3D7_0624900, Asp Peptidase family A1 domain of PM9) are displayed in blue. Signal peptide region is displayed in orange.* ***(C)*** *Concordant with the above observation, length of isoforms generated by SQANTI3 is higher for Female (LE) gametocytes (mean length: 1088 bp) compared to Female (HE) gametocytes (mean length: 941 bp). Among classifications of isoforms into structural categories by SQANTI3, Female (LE) gametocytes also have a higher percentage of FSM isoforms (50%) than ISM isoforms (11 %) compared to Female (HE) gametocytes (34% FSM and 32% ISM).* ***(D)*** *Female (LE) gametocytes are enriched in the set of genes described to be translationally repressed in female gametocytes (22). The plot for short read data shows the percentage overlap between genes detected in at least 50% of cells in each stage (number displayed on top of each bar) and the 511 translationally repressed transcripts from Lasonder et al., 2016. Similarly, the plot for long read data shows the percentage overlap between genes detected in at least 10% of cells in each stage (number displayed on top of each bar) and the 511 translationally repressed transcripts. A lower threshold of 10% is used for long read data due to the sparsity of the data.*

| | *D0* | *D3* | *D5* | *D10a* | *D10b* |
|---|---|---|---|---|---|
| *ZMWs input* | *2,393,746* | *2,441,922* | *2,688,353* | *2,701,479* | *5,674,531* |
| *Polymerase Reads* | *2,397,443* | *2,444,187* | *2,691,046* | *2,703,617* | *6,739,941* |
| *Polymerase Read Length (mean)* | *58,183* | *52,266* | *56,962* | *51,372* | *48,375* |
| *Polymerase Read N50* | *133,750* | *128,250* | *135,250* | *128,750* | *103,250* |
| *Longest Subread Length (mean)* | *6,912* | *8,421* | *5,881* | *4,827* | *7,873* |
| *Longest Subread N50* | *48,750* | *55,750* | *41,250* | *30,750* | *39,750* |
| *HiFi Reads* | *1,510,175* | *1,358,776* | *1,690,001* | *1,666,660* | *3,865,555* |
| *HiFi Yield (bp)* | *2,020,623,510* | *1,576,082,981* | *2,020,377,775* | *2,090,179,930* | *3,607,302,577* |
| *HiFi Read Length (mean, bp)* | *1,338* | *1,159* | *1,195* | *1,254* | *933* |
| *HiFi Number of Passes (mean)* | *21* | *21* | *21* | *20* | *22* |
| *Non-concatamer reads with 5' & 3' primers* | *1,363,750* | *1,194,182* | *1,481,605* | *1,487,368* | *1,809,375* |
| *FLNC (Non-concatamer reads with 5' & 3' Primers and Poly-A Tail)* | *1,173,386* | *1,002,039* | *1,275,605* | *1,269,429* | *1,624,466* |
| *Number of non-redundant transcripts* | *1,070,789* | *986,647* | *1,238,805* | *1,234,672* | *1,621,235* |

**Table S1. Output of long read sequencing from the 5 lab samples.**

| Strain_k6 | Strain_k7 | Strain_k8 | Counts |
|---|---|---|---|
| Doublet | Doublet | Doublet | 17 |
| Doublet | Doublet | Negative | 1 |
| Doublet | Doublet | SC3 | 2 |
| Doublet | Doublet | SC4 | 1 |
| Doublet | G6 | Doublet | 1 |
| Doublet | G6 | Negative | 1 |
| Doublet | G6 | SC8 | 10 |
| G1 | G5 | Doublet | 1 |
| G1 | G5 | SC5 | 229 |
| G2 | Doublet | Doublet | 1 |
| G2 | G6 | SC8 | 13 |
| G2 | G7 | SC3 | 47 |
| G2 | G7 | SC4 | 27 |
| G2 | Negative | Negative | 1 |
| G3 | Doublet | Doublet | 3 |
| G3 | Doublet | SC8 | 1 |
| G3 | G3 | Doublet | 1 |
| G3 | G3 | Negative | 1 |
| G3 | G3 | SC1 | 343 |
| G3 | G3 | SC8 | 1 |
| G3 | G6 | SC1 | 1 |
| G3 | G6 | SC8 | 23 |
| G3 | Negative | Negative | 6 |
| G3 | Negative | SC8 | 1 |
| G4 | G1 | SC6 | 82 |
| G5 | G2 | Negative | 2 |
| G5 | G2 | SC2 | 216 |
| G6 | G4 | Doublet | 1 |
| G6 | G4 | SC7 | 206 |
| Negative | Doublet | Doublet | 1 |
| Negative | G3 | SC1 | 2 |
| Negative | G6 | Doublet | 1 |
| Negative | G6 | SC8 | 6 |
| Negative | Negative | Negative | 8 |

**Table S2. Overlap of strain assignments from souporcell assuming 6 (Strain_k6), 7 (Strain_k7), and 8 (Strain_k8), strains.** *The highlighted rows include cells that are consistently assigned to the same strain cluster with the 3 iterations. The rest of the clusters are split when as the assumed number of strains is increased from 6 to 7 then 8*

| Package | Version | Built | | Package | Version | Built |
|---|---|---|---|---|---|---|
| Biostrings | 2.62.0 | R 4.1.1 | | RColorBrewer | 1.1-3 | R 4.1.3 |
| clusterProfiler | 4.2.2 | R 4.1.2 | | readr | 2.1.2 | R 4.1.3 |
| ComplexHeatmap | 2.10.0 | R 4.1.1 | | readxl | 1.4.0 | R 4.1.3 |
| data.table | 1.14.2 | R 4.1.2 | | reshape2 | 1.4.4 | R 4.1.2 |
| DEXSeq | 1.40.0 | R 4.1.1 | | rmarkdown | 2.14 | R 4.1.3 |
| dittoSeq | 1.7.0 | R 4.1.2 | | Rsubread | 2.8.2 | R 4.1.3 |
| dplyr | 1.0.8 | R 4.1.2 | | rtracklayer | 1.54.0 | R 4.1.1 |
| forcats | 0.5.1 | R 4.1.3 | | scales | 1.2.0 | R 4.1.2 |
| GenomicRanges | 1.46.1 | R 4.1.2 | | scmap | 1.16.0 | R 4.1.1 |
| Gviz | 1.38.4 | R 4.1.3 | | seriation | 1.3.6 | R 4.1.3 |
| gg3D | 0.0.0.9000 | R 4.1.2 | | Seurat | 4.1.0 | R 4.1.2 |
| ggfun | 0.0.6 | R 4.1.3 | | SingleCellExperiment | 1.16.0 | R 4.1.1 |
| ggplot2 | 3.4.2 | R 4.1.3 | | slingshot | 2.2.1 | R 4.1.3 |
| ggpubr | 0.4.0 | R 4.1.3 | | stringi | 1.7.6 | R 4.1.2 |
| ggthemes | 4.2.4 | R 4.1.3 | | stringr | 1.4.0 | R 4.1.2 |
| ggVennDiagram | 1.2.1 | R 4.1.2 | | swissknife | 0.38 | R 4.1.2 |
| gplots | 3.1.3 | R 4.1.3 | | tibble | 3.1.6 | R 4.1.2 |
| knitr | 1.39 | R 4.1.3 | | tidyr | 1.2.0 | R 4.1.2 |
| MetBrewer | 0.2.0 | R 4.1.3 | | tidyverse | 1.3.1 | R 4.1.3 |
| openxlsx | 4.2.5 | R 4.1.3 | | tradeSeq | 1.8.0 | R 4.1.1 |
| org.Pf.plasmo.db | 3.14.0 | R 4.1.2 | | viridisLite | 0.4.0 | R 4.1.2 |
| plotly | 4.10.0 | R 4.1.2 | | grDevices | 4.1.2 | R 4.1.2 |
| purrr | 0.3.4 | R 4.1.2 | | Matrix | 1.3-4 | R 4.1.2 |

**Table S3. List of software and versions used**

#### Supplementary Materials

**Data S1** (separate file) *Differential expression analysis between early trophozoites and early gametocytes (committed, stalk).* MAST package was used to run the DE testing, via *FindMarkers* function in Seurat version 4.1.

**Data S2** (separate file) *Outputs from tradeseq tests.* Spreadsheets displaying the complete output from tradeseq tests within the sexual stages along male and female lineages. Sheet 1 includes the description of the sheets and the figures that these tables correspond to. Sheets "developing stalk (heatmap)", "early stalk>early sex (heatmap)" and "branch > late (heatmap)" comprise the lists of genes in the heatmaps of Fig. S5, S6 and S7 respectively, in the order that they appear in the heatmap. Gene info from PlasmoDB v56 (description, genename, Ortholog group, protein and CDS length) was added for each gene in all the sheets.

**Data S3** (separate file) *Differential expression analysis between male-like and female-like gametocytes in the late stalk.* MAST package was used to run the DE testing, via *FindMarkers* function in Seurat version 4.1. Sheet 1 includes the description of the sheets and the figures that these tables correspond to. Sheet 2 contains the complete table of output from running the differential expression analysis. Sheet 3 contains the list of genes that pass thresholds of avg_log2FC > |0.4| and p_val_adj < 0.01). Gene info from PlasmoDB v56 (description, genename, Ortholog group, protein and CDS length) was added for each gene in both sheets 2 and 3.

**Data S4** (separate file) *Smoother plots, estimated by tradeSeq, for genes differentially expressed between male-like and female-like gametocytes in the late stalk.* Plots of log-transformed counts and the fitted values along male (yellow) and female lineages (violet) for genes that pass logFC and p_val_adj threshold from data S3.

**Data S5** (separate file) *UMAP gene clusters table.* Complete table of 5087 genes clustered based on their expression in single cells using kMeans clustering on the predicted UMAP dimensions, with k=15, and implemented by the scikit-learn package in python. Sheet 1 provides the legend with description of the sheets and the column names. Information in Sheet 2 includes UMAP dimensions and cluster assignment along with summary statistics from PlasmoDB (transcript ID, gene description, gene type, gene names, gene fitness scores). GO analysis output on the clusters using enrichGO package is provided in Sheet 3.

**Data S6** (separate file) *GENIE3 GRN Ranked Connections.* Complete output from Gene Regulatory Networks (GRN) training using the GENIE3 algorithm, on transcription factors from the ApicoTFdb database. The regulators were ranked after the total number of connections in the top 1.5 % percentile.

**Data S7** (separate file) *SQANTI Isoform Classification.* Sheet 1 contains the description of the sheets and the columns within each sheet. Sheet 2 contains the complete table of classification of the isoforms obtained following chaining of the filtered stage-wise SQANTI3 identified isoforms. Sheet3 contains the genome annotation file in GTF format for the chained isoforms. Field 2 contains the isoform identifier (PacBio for isoforms identified in this study). Field 9 includes a color code for the isoforms from this study, colored according to the level of support from long-read RNA-seq data (darker, more support). The color is based on the abundance values calculated from the counts for each isoform using cupcake's color_bed12_post_sqanti.py script.

**Data S8** (separate file) *DEXseq outputs.* Spreadsheets displaying the complete output from differential exon usage comparisons using DEXseq tests, along with identification of those that pass padj and logFC thresholds in the last column. Sheet 1 includes the description of the sheets and the description of the columns of the DEXseq outputs.

**Data S9** (separate file) *Strain and coarse-stage SNPs including QC categories.* Spreadsheets displaying the complete set of SNPs that come up after genotyping the different strain+coarse-stage pseudobulk and also strain pseudobulk clusters with freebayes including the different filter categories. Sheet 1 includes the description of the subsequent sheets with different pseudobulk classifications and the description of the columns of each sheet. Sheet 2 includes the entire set of strain pseudobulk SNPs together with QC metrics, Sheet 3 includes the clean subset of SNPs in sheet 2 used in IBS relatedness analysis. Sheet 4 includes the entire set of strain+coarse-stage pseudobulk SNPs together with QC metrics, Sheet 4 includes the clean subset of SNPs in sheet 4 used in relatedness analysis IBS and IBD.

**Data S10** (separate file) *Strain pairwise IBD.* Spreadsheets displaying the IBD results from hmmIBD. Sheet 1 includes the description of the subsequent sheets and the description of the columns of each sheet. Sheet 2 includes a summary of the pairwise relatedness between strains and sheet 3 details the IBD segment along the genomes for each pairwise comparison of strains.

**Data S11** (separate file) *Donor parasite stage genes, GO, FHE vs FLE unspliced_spliced ratios and average expression.* Spreadsheets displaying the genes that are expressed in each stage together with GO analysis, using clusterprofiler, for a set of these genes. Sheet 1 includes the description of the subsequent sheets. Sheet 2 indicates genes expressed in each stage with a gene considered expressed in a particular stage if it is detected in 25% of all the parasites in that stage. Sheet 3 to sheet 7 include the GO analysis for the genes only detected in female (HE), female (LE), female (HE + LE), male (HE + LE) and asexual stages respectively. Sheet 8 has the unspliced to spliced mRNA counts ratio for female (HE) and female (LE). Sheet 9 has the average expression of seurat log normalised genes in the different stages and proportion of cell in a stage expressing the gene.

**Data S12** (separate file) *Field versus lab MAST & tradeseq DE & GSEA.* Differential expression analysis between field vs laboratory cells using Seurat MAST and tradeSeq and corresponding GSEA. Sheet 1 includes the description of all subsequent sheets. Sheet 2 to 5 includes the DE for asexual, female (HE), female (LE) and male (HE) respectively. Sheet 6 to 9 includes the GSEA for asexual, female (HE), female (LE) and male (HE) respectively.

**Data S13** (separate file) *Field strains MAST & tradeseq DE.* Differential expression analysis between pairwise comparisons of field strains using MAST and tradeSeq. Sheet 1 includes the description of all subsequent sheets. Sheet 2 to 5 includes the DE for asexual, female (HE), female (LE) and male (HE) respectively.

**Data S14** (separate file) *SQANTI Isoform Classification for field parasites.* Sheet 1 contains the description of the sheets and the columns within each sheet. Sheet 2 contains the complete table of classification of the isoforms obtained following chaining of the filtered stage-wise SQANTI3 identified isoforms. Sheet3 contains the genome annotation file in GTF format for the chained isoforms, merged with Plasmodium falciparum 3D7 reference annotation from PlasmoDB v52. Field 2 contains the isoform identifier (PacBio for isoforms identified in this study, and VEuPathDB for annotated genes from PlasmoDB v52). Field 9 includes a color code for the isoforms from this study, colored according to the level of support from long-read RNA-seq data (darker, more support). The color is based on the abundance values calculated from the counts for each isoform using cupcake's color_bed12_post_sqanti.py script.

**Data S15** (separate file) *DEXseq outputs within field parasite stages.* Spreadsheets displaying the complete output from differential exon usage comparisons using DEXseq tests, along with identification of those that pass padj and logFC thresholds in the last column. Sheet 1 includes the description of the sheets and the description of the columns of the DEXseq outputs.
